# CiliaIO: Machine learning reveals spatial patterns of cilia beating dynamics in the zebrafish spinal cord

**DOI:** 10.64898/2025.12.10.693099

**Authors:** Ece Atayeter, Jason Ho, Talon G. Blottin, Ilyena B. Joe, Ron Sistrunk, Bo Zhang, Lilianna Solnica Krezel, Andreas Gerstlauer, John B. Wallingford, Ryan S. Gray

## Abstract

Motile cilia generate fluid flows that are essential for normal development and physiology. Cilia display diverse beating waveforms, and while pronounced defects are strongly associated with motile ciliopathies, subtler alterations also influence disease manifestations. Finer quantification of ciliary dynamics is therefore critical for understanding ciliopathies, but the heterogeneity of cilia beating makes accurate and robust characterization challenging. Here, we present CiliaIO, a machine learning-based tool for quantification of motile cilia morphodynamics. Using this platform, we discovered subtle but highly significant regional differences in ciliary waveforms in the zebrafish spinal cord. To demonstrate the tool’s efficacy, we used CiliaIO to capture and quantify subtle ciliary defects in a novel *bbs2* allele that causes a late-onset scoliosis phenotype. These results provide a workflow for additional fine-scale analyses of ciliary morphodynamics that will be important for understanding motile ciliopathy.

**SUMMARY STATEMENT:** Accurate and automatized cilia quantification is a challenging task. CiliaIO overcomes these challenges using a machine learning-based framework that automatically detects and quantifies cilia morphodynamics.

## INTRODUCTION

Motile cilia are microtubule-based organelles that drive directional fluid flow through precise, coordinated beating, and are essential for the normal development and function of several organ systems. Defects in cilia dynamics underlie motile ciliopathies, a group of disorders characterized by respiratory dysfunction, infertility, and left-right patterning defects (Praveen et al., 2015). Notably, many disease-causing ciliary gene variants produce no overt changes in beat frequency but instead manifest as subtle deviations in beating waveform and downstream fluid flow (Blanchon et al., 2020; Chioccioli et al., 2019; Knowles et al., 2014; Wohlgemuth et al., 2024). These differences are subtle but are functionally significant and remain incompletely understood.

Accurately characterizing such defects is thus an important challenge, and it is further complicated by the intrinsic heterogeneity in cilia behaviors, even in normal cilia (Boggon et al., 2026; Geyer et al., 2022; Juan et al., 2020). Such heterogeneity arises from a complex interplay of environmental, spatial, structural and molecular factors. At the structural level, motile cilia are built around the axoneme, a conserved structure consisting of nine peripheral microtubule doublets (Praveen et al., 2015; Satir et al., 2014). The presence or absence of a central microtubule pair defines two architecturally distinct configurations with markedly different beating patterns: cilia lacking the central pair (9+0) beat rotationally, whereas cilia containing a central pair (9+2) exhibit a planar waveform consisting of alternating power and recovery strokes (Afzelius, 1976; Nonaka et al., 1998).

Even within the 9+2 configuration, distinct waveforms can emerge from differences in the spatial regulation of dynein-driven microtubule sliding, as demonstrated in *Chlamydomonas* (Dutcher, 2020). Indeed, heterogeneity in normal cilia beating in healthy cells has been commonly noted, and linked not just to biological factors such as cilia maturation (Oltean et al., 2018) but also to technical factors in the analysis pipeline (Kempeneers et al., 2017). At the molecular level, the specific composition and axonemal positioning of accessory protein complexes further tune the precise characteristics of ciliary motion (Satir et al., 2014). For instance, disruption of radial spokes and other ciliary components can cause cilia to adopt a rotary rather than power-stroke beating form, including in zebrafish (Castleman et al., 2009; Knowles et al., 2014). Together, these structural and molecular changes can generate substantial variation in motile cilia behavior, making rigorous characterization of ciliary motility inherently challenging. Furthermore, quantification of comprehensive cilia morphodynamics in vivo is complicated by the fact that cilia are often imaged in close proximity to one another and to neighboring structures, making it difficult to isolate individual cilia and study their morphodynamic properties.

A number of computational tools have been developed to address these challenges, with accurate segmentation of cilia from the background serving as a foundational step common to all approaches. One class of methods uses classical computer vision methods such as intensity thresholding and edge-based segmentation strategies, that when optimized per dataset, achieve high segmentation accuracy (Hansen et al., 2021, 2018). However, these tools typically require manual intervention to crop individual cilia and exclude non-cilia structures, limiting their scalability and throughput. A second class of methods leverages pixel-based frequency analysis to distinguish motile cilia from stationary background components, exploiting the periodicity of ciliary beating as a discriminating signal (Jeong et al., 2022; Smith et al., 2012; Thouvenin et al., 2021). While these frequency-based approaches enable reliable cilia identification and beat frequency quantification, they do not produce frame-by-frame segmentation masks and skeletonization of individual cilia, which is a limitation that restricts fine-grained analysis of cilia morphology and motion dynamics.

Machine learning and deep neural networks have recently emerged as powerful tools for object detection and segmentation, capable of automating complex feature extraction from visual data without hand-crafted rules (O’Mahony et al., 2020). Specifically, You-Only-Look-Once (YOLO) is an efficient neural network that performs bounding box object detection (Redmon et al., 2016) and Segment-Anything Model (SAM) is a vision transformer capable of producing high quality segmentations without task-specific tuning (Kirillov et al., 2023). Building on these machine learning tools, we introduce CiliaIO, a machine learning-based cilia quantification pipeline that integrates YOLO-driven detection with SAM-based segmentation to automatically identify and delineate individual cilia in fluorescence video data. Segmentations are propagated across frames to continuously track each cilium, enabling downstream morphodynamics analysis including full skeleton tracking, beat frequency analysis, waveform characterization, and motion trajectory, with minimal manual intervention.

To demonstrate CiliaIO, we characterized the morphodynamics of Foxj1a+ motile monocilia lining the zebrafish spinal cord central canal (Ribeiro et al., 2017). In zebrafish, central canal cilia interact closely with key structures governing straight-axis development, including cerebrospinal fluid and the Reissner fiber (Bagnat and Gray, 2020). Disruption of these ciliated cells has been linked to impaired Reissner fiber assembly, abnormal cerebrospinal fluid flow, and scoliosis, yet underlying mechanisms remain poorly understood (Cantaut-Belarif et al., 2018; Grimes et al., 2016; Troutwine et al., 2020; Wang et al., 2022). To dissect these mechanisms, we applied CiliaIO to *bbs2^stl438^,* a recessive late-onset scoliosis mutant. Bbs2 is a member of the BBSome complex, which functions as an adapter for intraflagellar transport (Lechtreck et al., 2009; Nishimura et al., 2001). Applying CiliaIO to analyze both wild-type and *bbs2^stl438^* mutant animals, we uncovered distinct regional differences between dorsal and ventral central canal cilia and identified dynamic defects in *bbs2^stl438^* mutants invisible to visual inspection alone, directly linking specific cilia dynamics to Bbs2 function. Taken together, CiliaIO provides a robust framework for quantification of motile cilia morphodynamics through high-quality, frame-by-frame segmentations. Designed with the dual goals of automation and accuracy, CiliaIO is released as open-source software trained on zebrafish spinal cord motile cilia, with underlying models that can be fine-tuned to accommodate other cilium types, imaging modalities, and organisms.

## RESULTS

### CiliaIO combines a fine-tuned YOLO model and zero-shot SAM to accurately detect and segment spinal cord cilia

We sought to develop a pipeline that would rapidly identify and segment beating cilia *in vivo* in a way that allows fine-grained analysis of cilia morphology and dynamics (Fig. 1). We therefore looked to deep learning-based methods, which have advanced rapidly in recent years, offering automated feature analysis and potential advantages over classical computer vision approaches in accuracy (O’Mahony et al., 2020). Our new tool, CiliaIO, uses pattern recognition to identify and quantify cilia from confocal video based on their morphology. Specifically, CiliaIO uses a fine-tuned single-shot YOLOv11 (Jocher and Qiu, 2024; Redmon et al., 2016) object detection model to localize individual cilia within bounding boxes (Fig. 1B-C, G). These bounding boxes serve as region-of-interest prompts for the Segment Anything Model (SAM) (Kirillov et al., 2023), which produces high-quality segmentation masks for each detected cilium (Fig. 1C-D, H). By chaining YOLO detections as inputs to SAM within a unified ML pipeline, SAM operates only within regions where YOLO has identified a cilium, substantially reducing computation resources and speeding up inference.

**Figure 1.** CiliaIO produces fast and accurate quantification of cilia morphodynamics in zebrafish using state-of-the-art machine learning. **(A)** Schematic of central canal motile cilia imaging in zebrafish. **(B–F)** Overview of the CiliaIO pipeline from raw image input to morphodynamic metric output. **(B)** CiliaIO accepts raw confocal image sequences or videos in common formats (e.g., .mp4, .tif). **(C)** The YOLOv11 model detects individual cilia and generates a bounding box around each cilium. **(D)** The Segment Anything Model (SAM) ViT-H generates a segmentation mask for each cilium, using the bounding box from (C) as a spatial prompt. **(E)** Cilia skeletons are extracted using a morphological thinning algorithm. **(F)** Per-cilium tracking of a representative skeletal point and Fourier frequency analysis enable quantification of morphodynamic metrics. **(G)** The YOLOv11 detection model divides each frame into an S×S grid. For each cell, bounding boxes are generated alongside a class probability map derived from the training dataset, yielding high-confidence cilia bounding boxes. **(H)** The SAM ViT-H model processes each YOLO bounding box as a spatial prompt, encoding it alongside the image through an encoder-decoder architecture to produce a segmentation mask for each cilium. **(I)** Skeletonization is performed via morphological thinning, and the resulting skeleton is fit to an N-point spline curve. **(J)** Frequency-based motility metrics are extracted by applying the Fourier transform to the PCA major-axis displacement of a single skeletal midpoint for each cilium, converting the time-domain signal into a frequency power spectrum. **(K)** Representative time-domain signals and frequency power spectra for two individual cilia from CiliaIO analysis illustrating a cilium with low spectral entropy (top) and higher spectral entropy (bottom). The black dashed line indicates the dominant frequency while the blue dashed line indicates the centroid frequency.

Segmented cilia are then skeletonized via morphological thinning and smoothed by fitting a curve along their length (Fig. 1D-E, I; see Methods: CiliaIO Pipeline for details). Individual cilia are then tracked across frames using the ByteTrack (Zhang et al., 2022) multi-object tracking algorithm. This enables consistent cilia IDs to be maintained for YOLO bounding boxes across frames by first using high-confidence detections with existing tracks, then recovering temporarily lost tracks (including overlapping or occluded cilia) using lower-confidence detections.

To further improve stable cilia identification across frames, we developed a custom post-processing methodology grounded in the presumption that individual cilia remain anchored at the basal body. In the first frame, each detected cilium is assigned a unique numerical identifier (0 through N-1). In subsequent frames, cilia are re-identified by matching each new detection to the spatially closest object detection (cilium) from prior frames. Detections displaced more than a user-defined number of pixels from any prior position are assigned new identities, as this exceeded the expected inter-frame displacement of an anchored cilium (we generally have used 3.2 µm here).

To account for brief detection failures caused by focal drift, motion blur, or transient occlusion, a cilium absent from detection is retained in the tracking record for up to a user-defined number of consecutive frames before removal, preventing short gaps from fragmenting a cilium’s tracking history (we have generally used 5 frames here). Finally, to prevent the same cilium from being counted more than once, any two detections whose bounding boxes overlapped more than 50% were merged into a single ID, with the older track taking precedence. As imaging was performed in 2D, this merging step provides a practical solution that can be tuned by the user for managing closely spaced cilia within the focal plane, though some ambiguity may remain in regions of particularly high cilia density (see Methods: CiliaIO Pipeline for details).

With a consistent identity bounding box prompt passed from YOLO to SAM, CiliaIO can decouple motile cilia that beat in proximity or cross paths. These events pose a significant challenge for classical computer vision algorithms, which struggle in noisy environments due to their reliance on clear boundaries for accurate segmentation (Xu et al., 2024). The ML-based formulation underlying CiliaIO combines positional cues from tracking with per-frame pattern recognition to accurately identify and distinguish individual cilia, even when partially overlapping (Video S1 and S2). In cases where YOLO generates partially overlapping bounding boxes, SAM executes independent inference passes for each box. This decouples contiguous foreground objects by forcing the model to complete the most probable segmentation boundary within each box, even when cilia share similar intensity gradients compared to the background.

### CiliaIO uses skeleton-based tracking to quantify ciliary beating dynamics

With cilia accurately detected and segmented, CiliaIO then skeletonizes each cilium and tracks individual skeletal points over time, enabling quantification of both morphology and beating dynamics. Users can choose to track either a single representative point or the full skeleton, offering flexibility for different analytical needs (Fig. 1F, I and Fig. 2A, B). To capture frequency-domain information, CiliaIO computes the principal component analysis (PCA) major-axis displacement of skeletal X-Y points over time, constructing a time-domain signal that reflects the physical motion of the cilium. Frequency metrics are derived from these time domain signals, capturing the representative beating dynamics along its length. Similar to existing cilia quantification methods (Hansen et al., 2018; Jeong et al., 2022; Smith et al., 2012; Thouvenin et al., 2021), the Fourier transform is applied to time-domain signals to extract frequency-domain metrics including dominant frequency, centroid frequency, and spectral entropy (Fig.1F, J-K, Fig. 2A’-B’’, see Methods: Quantification of Morphodynamic Metrics).

**Figure 2.** Ciliary motion reveals distinct oscillatory behaviors between low- and high-entropy cilia. **(A–A″)** Low-entropy ciliary motion. **(A)** Temporal skeletal overlays of ciliary trajectories, color-coded across a 2-second video, showing a stereotyped and highly coordinated beat pattern. Color gradient represents time. **(A′)** Time-domain signal of ciliary displacement showing regular, periodic oscillations with low variability. **(A″)** Frequency-domain power spectrum revealing a single sharp, dominant spectral peak, indicative of highly rhythmic beating. **(B–B″)** High-entropy ciliary motion. **(B)** Temporal skeletal overlays color-coded across a 2-second video. **(B′)** Time-domain signal showing irregular oscillations with greater variability. **(B″)** Frequency-domain power spectrum displaying a broad, distributed spectral profile, reflecting dysrhythmic beating with higher entropy fluctuations compared with the lower entropy, rhythmic beating of the cilium in (A). Scale bars: 1 µm.

For each analyzed video, CiliaIO generates three primary outputs. First, an annotated video displaying, for each frame, YOLO-detected bounding boxes with unique cilia object identifiers, SAM-generated segmentation masks, extracted skeletons, and temporarily tracked midlines. Second, a comma-separated values file cataloguing each cilium identifier alongside its corresponding morphodynamic measurements, including beat frequency, amplitude, and additional derived metrics. Third, Python data structures encoding each cilium identifier to its associated bounding box centers, skeleton midpoints, and per-frame full skeletons, supporting further downstream computational analyses.

CiliaIO achieves efficient runtime performance: on consumer-grade hardware (AMD Ryzen 5800X3D, NVIDIA GTX 1080 Ti, 32 GB RAM), a 250-frame zebrafish spinal cord microscopy video with 29 cilia per frame was processed in 11.6 minutes; on server-class hardware (AMD EPYC 7763, NVIDIA A100, 256 GB RAM), the same file was processed in 4.3 minutes.

### CiliaIO achieves robust cilia detection and segmentation across diverse zebrafish samples

We then trained CiliaIO for a specific real-world application: accurately identify and localize individual motile cilia in zebrafish central canal confocal microscopy images (Fig. S1). CiliaIO uses a pre-trained YOLOv11m model (Jocher and Qiu, 2024; Redmon et al., 2016) that was further fine-tuned on a cilia-specific dataset. To construct this dataset, confocal microscopy videos of zebrafish central canal motile cilia from 7 animals with different genotypes were annotated frame-by-frame in Label Studio (Tkachenko et al., 2020), with bounding boxes manually drawn around each visible cilium. In total 1,344 video frames were labeled, yielding 32,167 cilia annotations across the full dataset (A full description of the dataset is included in the methods).

The dataset was divided into training, validation, and test subsets at the video level, ensuring that all frames from a given video were assigned exclusively to one subset. This design prevents data leakage by ensuring the model is never evaluated on frames from videos used during training. The training set optimizes the cilia detection model. The validation set monitors model performance during training to determine when to stop training, reducing overfitting to the training data. The test set provides a final assessment of model performance on data entirely absent from both prior subsets. Of the 7 videos, 1 video was reserved as the test set, while the remaining 6 videos were split 80/20 into the training set (4 videos) and a validation set (2 videos). The validation set included 2 different genetic backgrounds to prevent overfitting to a single genotype.

The YOLOv11m model was fine-tuned for 300 epochs with its training and validation losses demonstrating continual learning with no overfitting (Fig. S1B-G; see Methods). On the test set, the model demonstrated strong detection performance. Accuracy was assessed using an F1 score, a combined measure of precision (the fraction of model detections corresponding to true cilia) and recall (the fraction of true cilia successfully detected). A cilium is identified correctly by the model when at least 50% of the total area of the predicted bounding box overlaps with the manually drawn bounding box.

This model achieved an F1 score of 86% at a confidence threshold of 0.63 (Fig. S1L). An F1 score of 86% indicates that the model reliably detects the vast majority of cilia while maintaining a low false detection rate. The model also achieved a mean average precision of 91% at 50% overlap (mAP@50; Fig. S1M), another metric used to evaluate object identifier models. On the test set with a confidence threshold of 0.25, 1,403 cilia were correctly identified, with 104 false negatives and 359 false positives (Fig. S1N). The large number of false positives comes from the low confidence threshold cutoff. Together, these results confirm that the fine-tuned model is well suited for automated cilia detection in the spinal cord central canal from confocal microscopy images.

To further assess robustness and generalizability, we performed stratified 3-fold cross validation for the YOLO fine-tuning procedure. As before, dataset splits were performed at the video level to prevent data leakage. In each fold, 4 videos were used for training and 2 videos were reserved for validation. The same test set was used for final evaluation across all folds. For each fold, F1-confidence and precision-recall curves were computed, and the mean and standard deviation were reported across folds (Fig. 3A-D). Across validation folds, the model achieved a peak F1-score of 78% at a confidence threshold of 0.24, and a mAP@50 of 80.5% (Fig. 3A, B). The variability observed across validation folds likely reflects biological heterogeneity between the samples and suggests that a larger training dataset would improve consistency. On the test set, model performance was higher and more stable across folds with a peak F1-score of 89% at a confidence threshold of 0.45 and mAP@50 of 93% (Fig. 3C, D). Together, these results demonstrate reliable detection of cilia on unseen biological samples and suggest that expanding the training dataset would further reduce variability and improve generalization. Any further results in the paper are performed using the fine-tuned YOLOv11 model described before the cross-validation experiment.

**Figure 3.** Training performance and segmentation accuracy of the fine-tuned YOLO model, and SAM-generated masks compared against manual ground truth. **(A)** F1-confidence curve for the validation set, showing F1 score as a function of confidence threshold across three cross-validation folds with the mean curve (±1 SD shading). Peak mean F1 score: 78% at a confidence threshold of 0.24. **(B)** Precision-recall curve for the validation set. Mean area under the curve (mAP@50): 80.5%. **(C)** F1-confidence curve for the test set across three folds. Peak mean F1 score: 89% at a confidence threshold of 0.45. **(D)** Precision-recall curve for the test set. Mean mAP@50: 93%. **(E)** Distribution of Intersection over Union (IoU) scores for all segmented cilia (N = 1,269; mean = 0.669 ± 0.136). Dashed line indicates the mean. **(F)** Distribution of Dice–Sørensen coefficient (DICE) scores for all segmented cilia (N = 1,269; mean = 0.793 ± 0.11). Black dashed line indicates DICE = 0.7. Red dashed line indicated the mean. Analysis in (E–F) collected from 7 animals comprising 1,269 cilia masks. **(G)** Representative confocal image showing SAM-predicted segmentation masks (magenta) overlaid on manual ground-truth masks (green) for individual cilia. White bounding boxes indicate manually labeled regions used as prompts to SAM; DICE scores are displayed for each object. Scale bar: 2.5 µm.

Having established reliable cilia detection, we next evaluated whether SAM could segment cilia accurately without task-specific training. Unlike supervised models that require task-specific labeled training data, SAM operates in a zero-shot manner, meaning it can generalize to novel objects by leveraging visual patterns learned from a large and diverse image dataset (Kirillov et al., 2023). To evaluate SAM’s zero-shot segmentation performance on cilia, we generated a ground truth dataset by manually labeling segmentation masks for 10 randomly selected frames per video from the YOLO datasets described above, yielding 1269 hand-labeled cilia masks. We utilized the existing bounding boxes from the dataset to define the regions of interest prompts for SAM (Fig. 3G). Segmentation accuracy was quantified using two standard metrics: Intersection over Union (IoU) and the Dice–Sørensen coefficient (DICE) (Kocak et al., 2025). IoU provides a strict measure of accuracy by calculating the ratio between the area of overlap and the total area of both the predicted and manual masks. IoU measures the ratio of the intersection to the union of predicted and reference mask areas, providing a strict assessment that penalizes false positive pixels (Kocak et al., 2025). DICE calculates the ratio of twice the intersection area to the sum of total pixels in both masks, and is widely used in biological imaging for its sensitivity to true positive overlap (Kocak et al., 2025). Zero-shot SAM achieved a mean IoU score of 0.67 ± 0.14 and mean DICE score of 0.79 ± 0.11 (Fig. 3E, F). Across the segmentation dataset, 84.6% of cilia (1,074 of 1,269) met or exceeded the broadly recognized benchmark DICE score of 0.7 for high segmentation accuracy in biological imaging (Boehringer et al., 2023; Zijdenbos et al., 1994). These results, corroborated by visual inspection, confirm that SAM provides accurate cilia segmentation of cilia without any dataset-specific fine-tuning.

### CiliaIO length and frequency outputs are consistent with pixel-based frequency quantification and manual length quantification

To assess the end-to-end accuracy of CiliaIO, we benchmarked its outputs against dominant/main frequency quantification using an established pixel-based Fast Fourier Transform (FFT) methodology (Thouvenin et al., 2021) and manual length quantification in ImageJ (Schindelin et al., 2012). Accuracy was quantified using Mean Absolute Percentage Error (MAPE), which measures the average absolute percentage deviation between predicted and observed values. In this analysis, the manual length annotations and pixel-based FFT methodologies served as ground-truth benchmarks for their respective comparisons.

For frequency validation, CiliaIO outputs were compared against cilia scripts adapted from a pixel-based FFT methodology, developed by Thouvenin et al., 2021 (Fig. 4A, B). A cohort of 3 animals consisting of 49 individual cilia was utilized for frequency validation. CiliaIO dominant frequency measurements yielded a mean absolute deviation of 0.5 Hz, corresponding to a MAPE of 8.46% relative to the pixel-based FFT baseline (Fig. 4C, D). The majority of this error was concentrated in three cases with deviations exceeding 75% error (Fig. 4D). Excluding these outliers, agreement between the two methods was strong across the remainder of the dataset.

**Figure 4.** Benchmarking of CiliaIO against pixel-based FFT beat frequency estimation and manual cilia length quantification. **(A)** Representative CiliaIO output showing individual cilia detected and segmented from a confocal image. Scale bar: 1 µm. **(B)** Corresponding output from Thouvenin et al. pixel-based FFT analysis for the same field of view; colors encode beat frequency (scale: 0–25 Hz). **(C)** Histogram of absolute error in dominant/main beat frequency between CiliaIO and Thouvenin et al. estimates; the majority of cilia show errors near 0 Hz, with a mean absolute error of 0.50 Hz. **(D)** Histogram of mean absolute percentage error (MAPE) in dominant/main beat frequency; MAPE = 8.46%. Beat frequency comparison performed on 3 animals (2 wild-type, 1 *bbs2^stl438^*) comprising 49 cilia. **(E)** Histogram of absolute error in cilia length between CiliaIO and manual annotations; mean absolute error = 0.13 µm. **(F)** Histogram of MAPE in cilia length; MAPE = 3.57%. Length comparison performed on 3 animals (2 wild-type, 1 *bbs2^stl438^*) comprising 47 cilia.

Closer inspection showed that these high-error cilia exhibited complex beat dynamics inconsistent with simple periodicity. Rather than displaying a singular, dominant frequency, these cilia alternated stochastically between different beat frequencies, showcasing the high level of heterogeneity within the same animal. Because CiliaIO tracks discrete skeletal points of a cilium over time, it is sensitive to phase discontinuities that arise from non-stationary signals. Pixel-based FFT methods, which capture global intensity shifts across spatial regions, are less susceptible to such phase effects; however, their spatial averaging can obscure beat stochasticity. In both cases, a single dominant frequency does not fully represent the temporal complexity of the signal. Ultimately, when examining complex dynamics, it is important to examine the entire Power Spectral Density (PSD) from the Fourier transform output (Fig. 2A’, B’) rather than rely solely on single-frequency summary statistics, regardless of the quantification method used.

For manual length validation, a separate, non-overlapping cohort of videos from 3 animals consisting of 47 individual cilia were analyzed. Each cilium’s length was measured in three randomly selected frames per video and averaged. CiliaIO length measurements yielded a mean absolute deviation of 0.13 μm per cilium, corresponding to a MAPE of 3.57% relative to the manual cilia length quantification, reflecting close agreement between manual measurements and CiliaIO output (Fig. 4E, F). Together, these benchmarking results demonstrated that CiliaIO quantifies cilia frequency and length with accuracy comparable to established methods.

### CiliaIO reveals spatial patterning of cilia waveforms in the zebrafish spinal cord

Prior studies of central canal cilia focused primarily on cerebrospinal fluid flow and ciliary beating during early development, spanning 24 to 32 hours post-fertilization (hpf) (Borovina et al., 2010; Kramer-Zucker et al., 2005; Sternberg et al., 2018; Thouvenin et al., 2020). These studies suggest that motile cilia lining the ventral wall of the central canal are the primary drivers of directional fluid flow (Sternberg et al., 2018; Thouvenin et al., 2020). However, these studies examined motile cilia at early developmental stages, before the spinal cord undergoes substantial structural rearrangement (i.e. dorsal collapse) (Ribeiro et al., 2017; Tait et al., 2020). Since these rearrangements may also influence motile cilia morphodynamics, we extended our analysis to motile ciliated cells at 4 dpf, capturing cilia behavior after this morphogenetic transition. Using the *Tg(Foxj1a:Arl13b-GFP)^ut22^* transgenic line, which labels Foxj1a+ motile cilia with a GPF-tagged Arl13b protein while leaving neighboring non-motile cilia unlabeled (Konjikusic et al., 2018), we performed time-lapse confocal imaging of motile monocilia extending from both the ventral roof plate and dorsal floor plate cells lining the central canal (Fig. 5A and Video S2).

**Figure 5.** Comparative *CiliaIO* analysis reveals distinct dynamics of dorsal and ventral Foxj1a+ central canal cilia. **(A)** Representative confocal image of wild-type Foxj1a⁺ central canal cilia at 4 dpf with CiliaIO bounding boxes and segmentation masks overlaid; dorsal cilia (blue) and ventral cilia (beige) are assigned unique numerical IDs. Scale bar: 2 µm. **(A′)** Temporal skeletal overlays of ciliary trajectories manually compiled together to correspond to ciliary positions and skeleton IDs in (A), color-coded across a 2-second video. Color gradient represents time. Scale bar: 2 µm. **(B)** Three-dimensional scatter plot of individual wild-type cilia showing dominant beat frequency (Hz), mean beat amplitude (µm), and spectral entropy for dorsal (blue) and ventral (beige) cilia. Numbered cilia correspond to those labeled in (A–A′). **(C)** 2D principal component analysis of 6 wild-type cilia morphodynamic features (standard deviation of straightness, max amplitude, average speed, entropy, dominant frequency, mean straightness). For each fish, the median feature was extracted from each of the dorsal and ventral cilia populations. The PCA shows distinct clustering of dorsal (blue) and ventral (beige) populations. PC1 explains 61.8% variance, and PC2 explains 20.9% of variance in cilia morphodynamics; distinct clusters assessed by PERMANOVA (p-value = 0.0001), with Paired Wilcoxon test between dorsal and ventral cilia groups (PC1 p-value= 0.00195, PC2 p-value = 0.695). **(D–G)** Violin SuperPlots comparing dorsal (blue) and ventral (beige) cilia for each fish; individual data points are color-coded by animal. **(D)** Cilia length (µm) is comparable between dorsal and ventral cilia. **(E)** Centroid beat frequency (Hz) is higher in dorsal than ventral cilia. **(F)** The mean axonemal straightness is higher for dorsal cilia. **(G)** The variation of axonemal straightness during the imaging interval is greater in ventral than dorsal cilia. Violin box represents interquartile range; white line indicates median. Paired Wilcoxon signed-rank test was used to compare the median values of dorsal and ventral cilia with each fish. p-values: ** ≤ 0.01. Experimental data from 10 wild-type animals comprising 117 dorsal and 102 ventral cilia; ≥ 3 independent replicates. Brightness and contrast adjusted for visibility in (A-A’).

Morphologically, dorsal and ventral cilia exhibited similar distributions of length and area (Fig. 5D and S2A). However, principal component analysis of dynamic features revealed that the motility in dorsal and ventral cilia was strikingly different for 6 morphodynamic metrics with different component coefficients (Fig. 5C and Fig. S2H). Three-dimensional graphs mapping individual cilia along dominant frequency, mean amplitude and spectral entropy also suggested that the dorsal and ventral cilia beating behavior cluster into distinct groups (Fig. 5B). Dorsal cilia tend to beat at a higher centroid frequency but with reduced amplitude compared with ventral cilia (Fig. 5B, E and Fig. S2). Dorsal cilia also exhibited higher relative spectral entropy, whereas ventral cilia displayed a lower-entropy beat pattern (Fig. 5B and Fig. S2).

Consistent with these differences, temporal overlays of ciliary skeletons revealed markedly distinct shape distributions between dorsal and ventral cilia (Fig. 5A’), indicating that the two populations exhibit different beating waveforms. To test this directly, we quantified axonemal shape dynamics using straightness as a proxy for ciliary bending. Dorsal cilia were significantly straighter over the time course of imaging and exhibited lower variability in straightness, while ventral cilia displayed greater bending and higher variability in straightness (Fig. 5F, G).

Corroborating the CiliaIO findings, the frequency difference between dorsal and ventral cilia was independently captured by pixel-based FFT quantification (Thouvenin et al., 2021) (Fig. S2G). Together, these findings demonstrate that Foxj1a+ central canal motile cilia display a robust spatial segregation of beating characteristics at 4 dpf, with consistent differences across multiple dynamic metrics. These findings reveal that positional identity along the dorsal-ventral axis is a determinant of ciliary beating behavior in the zebrafish spinal cord central canal.

### Late-onset scoliosis in *bbs2^stl438^* mutants is associated with dorsal-specific defects in the dynamics of central canal cilia and heterogeneous defects of the Reissner fiber

Motile cilia defects in the central canal are associated with whole-body scoliosis in zebrafish (Grimes et al., 2016; Cantaut-Belarif et al., 2018; Wang et al., 2022). To test the utility of CiliaIO in a real-world experimental setting, we examined cilia morphodynamics in a previously uncharacterized scoliosis mutant displaying a recessive scoliosis phenotype that presents substantially later than other previously characterized scoliosis mutants (Troutwine et al., 2020; Wang et al., 2022) (Fig. 6A-E). Genomic sequencing, mapping, and cloning of full-length mutant cDNA revealed a 3’ splice acceptor site mutation in the zebrafish homolog of the Bardet-Biedl syndrome 2 gene (*bbs2^stl438^*), resulting in a 21 base pair deletion of exon 14 in *bbs2^stl438^* transcript (Fig. 6F-G and Fig. S3A-E). This mutation produces a predicted in-frame 7-amino acid deletion within the highly conserved platform domain of the Bbs2 protein (Chou et al., 2019; Singh et al., 2020; Traub et al., 1999) (Fig. 6G, H and Fig. S3F). To confirm that the *bbs2^stl438^* splice acceptor allele was deleterious, we performed a complementation test with a published *bbs2* allele (Song et al., 2020). Transheterozygous animals failed to complement the scoliosis phenotype, confirming both alleles disrupt *bbs2* function (Fig. S3G-I).

**Figure 6.** An intronic splice-disrupting mutation in *bbs2* (*stl438*) causes late-onset scoliosis in zebrafish. Brightfield **(A–B′)** and microCT **(C–D′)** images of wild-type (A, A′, C, C′) and *bbs2^stl438^* mutant (B, B′, D, D′) zebrafish at 60 dpf, shown in lateral (A–D) and dorsal (A′– D′) orientations. Mutants display overt axial curvature in both planes. Brightfield images stitched from multiple panels. **(E)** Longitudinal survey of body-curvature incidence across multiple *bbs2^stl438^* crosses, demonstrating late-onset penetrance that is not influenced by maternal effects. *bbs2^stl438^*^/+^ incrosses (n = 226 at 5 dpf, n = 114 at 60 dpf); MZ*bbs2^stl438^* crosses (n = 243 at 5 dpf, n = 96 at 60 dpf); male *bbs2^stl438^*^/+^ × female *bbs2^stl438^* crosses (n = 162 at 5 dpf, n = 102 at 60 dpf). Compiled from ≥ 2 crosses per condition. **(F)** Sequence alignment of *bbs2* cDNA from wild-type and *bbs2^stl438^* mutants showing a 21-base-pair deletion in the mutant. **(G)** Predicted Bbs2 protein alignment showing a 7-amino acid deletion (ENSDART00000182692.1:r.[508_514del]) in the mutant. **(H)** Bbs2 protein structure (UniProt: Q98SP7) and domain schematic highlighting the region deleted in *bbs2^stl438^* (arrow) within the platform domain (pf); domains shown are β-Propeller, hx (heterodimerization), GAE (γ-adaptin ear), pf (platform), and CC (coiled coil). **(I–L″)** Confocal imaging of wild-type (I–J″) and *bbs2^stl438^* (K–L″) central canal cilia at 4 dpf. Foxj1a⁺ motile cilia visualized using *Tg(Foxj1a:Arl13b-GFP)^ut22^* (green; I, K) and counterstained with a zebrafish-specific Arl13b antibody (magenta; I′, K′) to label all Arl13b-expressing cilia. Merged channels shown in I″ and K″; insets (J–J″, L–L″) show higher-magnification views of the dashed boxed regions. *bbs2^stl438^* mutants display axonemal accumulation of both transgenic and endogenous Arl13b (K–L″), concentrated at cilia tips and midpoints (L–L″). Analysis from 5 wild-type and 3 *bbs2^stl438^* animals. **(M–P)** Confocal imaging of Reissner fiber at 7 dpf using the *scospondin-GFP^ut24^* transgenic line. (M) Wild-type Reissner fiber forms a continuous structure. (N–P) *bbs2^stl438^* mutants display a spectrum of Reissner fiber defects. **(Q)** Categorization of Reissner fiber phenotypes across brain, hindbrain ventricle, and tail regions in wild-type and *bbs2^stl438^* animals at 7 dpf. Each column represents one animal; each row represents one anatomical region. Analysis from 6 wild-type and 15 *bbs2^stl438^* animals; ≥ 3 independent replicates. Scale bars: 5 mm in A-D’; 5 µm in I-I’’, K-K’’; 1 µm in J-J’’and L-L’’; and 5 µm in M-P.

Scoliosis in zebrafish is linked to the Reissner fiber, a continuous glycoprotein fiber in the central canal (Cantaut-Belarif et al., 2018; Troutwine et al., 2020; Wang et al., 2022). We therefore compared Reissner fiber integrity in curved and normal *bbs2^stl438^* mutant fish at 4 and 7 dpf. We found that mutants with straight axes displayed a typical Reissner fiber, while mutants with curved axes showed a range of defective phenotypes at both developmental stages (Fig. 6M-Q and Fig. S3K).

Formation of the Reissner fiber requires cilia-mediated fluid flow (Troutwine et al., 2020; Cantaut-Belarif et al., 2018; Wang et al., 2022), so we next imaged Foxj1+ motile cilia using the *Tg(Foxj1a:Arl13b-GFP)^ut22^* transgene in *bbs2^stl438^* mutants. Cilia were largely normal in overall morphology, though a subset displayed bulges of accumulated Arl13b-GFP protein, similar to the dysmorphic cilia reported in the airway epithelium *of BBS2* mutant mice (Shah et al., 2008) (13.31%; n= 266) (Fig. 6I-K and Fig. S3J). Immunostaining against endogenous zebrafish Arl13b protein (Duldulao et al., 2009) confirmed this phenotype (Fig. 6I-L”).

Given the relatively mild morphological cilia phenotype in *bbs2^stl438^* mutants, we next examined motile cilia dynamics but we observed no overt, qualitative defects. However, CiliaIO analysis in our mutant fish (Videos S3-S6, Fig. 7, Fig. S4 and Fig. S5) showed dorsoventral separation of ciliary behavior in *bbs2^stl438^* comparable to wild-type animals based on principal component analysis (Fig. 7C). The effects of *bbs2* loss were region-specific, predominantly affecting a subset of dorsal cilia (Fig. 7A-B’). Dorsal cilia in *bbs2^stl438^* mutants displayed increased centroid beating frequency and an increased variability of axonemal straightness relative to wild type (Fig. 7D-E) which were unaltered in ventral cilia (Fig. 7D’-E’). This selective disruption of dorsal, but not ventral, cilia dynamics highlights the dorsoventral asymmetry we observed in normal cilia and suggests a region-specific role for Bbs2 in maintaining the dorsal ciliary waveform.

**Figure 7.** *bbs2^stl438^* mutants display subtle dorsal-specific defects in central canal cilia dynamics at 4 dpf. **(A–B’)** Representative CiliaIO skeletal overlays for Foxj1a⁺ central canal cilia in wild-type (A, A’) and *bbs2^stl438^* (B, B’) animals at 4 dpf. Color gradient represents time. Dorsal cilia (A, B): skeletal overlays reveal altered beat trajectories in a subset of *bbs2^stl438^* cilia (B) compared with wild type (A). Ventral cilia (A’, B’): beat patterns are comparable between genotypes. **(C)** 2D principal component analysis of 6 morphodynamic features (change in straightness, max amplitude, average speed, entropy, dominant frequency, mean straightness) for both wild type (represented as circle) and *bbs2^stl438^* (represented as cross) mutants. *bbs2^stl438^* projected on wild-type PCA. **(D–E′)** Violin SuperPlots comparing wild-type and *bbs2^stl438^* cilia within dorsal (D–E) and ventral (D′–E′) populations; individual data points are color-coded by animal. (D, D′) Centroid beat frequency (Hz) is increased in *bbs2^stl438^* dorsal cilia (D) but unaffected in ventral cilia (D′). (E, E′) Variability in axonemal straightness is increased in *bbs2^stl438^* dorsal cilia (E) but ventral cilia are unaffected (E′). Mann-Whitney U tests performed between wild-type and *bbs2^stl438^* medians for each fish within each dorsoventral group. Violin box represents interquartile range; white line indicates median. p-values: * ≤ 0.05. Wild-type: 10 animals, 117 dorsal and 102 ventral cilia. *bbs2^stl438^*: 10 animals, 132 dorsal and 117 ventral cilia. ≥ 3 independent replicates. Scale bars: 1 μm in A-B’.

## DISCUSSION

The development of CiliaIO was motivated by the need for both automated and accurate cilia quantification. To achieve this, we leveraged state-of-the-art machine learning architectures and fine-tuned a detection model capable of automatically recognizing cilia in the zebrafish central canal and generating segmentation masks. From these masks, we skeletonize individual cilia and compute metrics that capture motile cilia morphology and dynamics. In its current form, CiliaIO was trained specifically for spinal cord mono-motile cilia imaged by confocal microscopy and performs optimally in this context. We anticipate that the CiliaIO workflow can be extended to other tissue types and imaging modalities through additional training, which we aim to pursue in future work, alongside releasing CiliaIO as open-source software so that it can be adapted for diverse research applications in the cilia community.

CiliaIO resolves fine scale, biologically meaningful differences in ciliary morphodynamics that are not easily assessed by conventional qualitative assessments of ciliary behavior. Using CiliaIO, we observed spatial differences in ciliary behavior between dorsal and ventral populations in both wild-type and *bbs2^stl438^* animals at early larval stages. These differences may arise from several contributing factors. First, dorsal and ventral cilia may harbor distinct axonemal architectures. Dorsal cilia may lack a central pair and adopt a 9+0 configuration, while the waveform characteristics of ventral cilia are consistent with the presence of a central pair. Second, the composition and axonemal positioning of proteins that fine-tune the beating waveform may differ between the two populations. Third, the geometry of the central canal may differentially orient cilia, giving rise to distinct beating behaviors. Future structural and molecular analysis, including transmission electron microscopy and immunostaining of ciliary components, will be needed to directly assess whether differences in axonemal architecture or protein composition underlie the distinct ciliary waveforms.

We further showed that *bbs2^stl438^* mutants harbor subtle but quantifiable alterations in ciliary motility restricted to the dorsal central canal, with the loss of *bbs2* function selectively disrupting the unique waveform-generating mechanisms of dorsal cilia without substantially affecting ventral cilia dynamics. Although these changes are subtle and therefore difficult to capture, CiliaIO successfully detected quantitative differences that may have biologically meaningful effects on Reissner fiber assembly and, subsequently, for spine morphogenesis. These findings further underscore the call to characterize motile cilia not only by beat frequency but through fine-scale waveform analysis (Chioccioli et al., 2019), which can expose mechanistic defects that would otherwise go undetected.

An open problem is the handling of missing or discontinuous tracking data. At our standard confocal imaging rate (49.9 Hz), a cilium that transiently leaves the imaging plane or falls below detection confidence is assigned a new track ID upon reacquisition. We identify and remove these duplicate-ID fragments during quality control (∼14% of tracks in this dataset), retaining only the identity with the greatest number of consecutively tracked frames to maximize the time-domain signal and minimize truncation error in the Fourier transform. While most our cilia tracks length captured the full 249-frame acquisition, there are some track-length variations across our datasets (mean 195, median 249 frames), which could affect downstream frequency-domain measures.

Future iterations of CiliaIO could incorporate a deep learning-based imputation method such as DISK (Rose et al., 2026) to recover missing ciliary track segments. Because the cilium skeleton is represented as a set of spatially ordered, physically continuous points, it is well suited to DISK’s approach, offering a ready-to-implement imputation step to reconnect fragmented tracks rather than discarding them, ultimately increasing usable data per cilium and reducing track-length variability as a confound in comparative frequency and entropy analyses. We view integration of gap-imputation methods as a promising direction for future development of CiliaIO.

Like other frequency-based ciliary quantification tools, CiliaIO uses Fourier analysis to quantify the frequency of ciliary beat signals (Hansen et al., 2018; Jeong et al., 2022; Smith et al., 2012; Thouvenin et al., 2021). However, the Fourier transform assumes that signals approximate a pure sinusoid, which can introduce bias in certain metrics. For example, non-sinusodial sawtooth-like beating waveforms that integrate harmonic frequency components in the frequency domain (Saggiorato et al., 2017), could artificially inflate spectral entropy independent of true signal irregularity. For this reason, spectral entropy is best interpreted as a comparative measure between cilia populations rather than as a standalone absolute measure of irregularity.

The current implementation of CiliaIO does not yet compute the full range of quantitative descriptors reported in prior cilia studies. Metrics such as fluorescence intensity profiles along the axoneme, arc length, and cross-correlations between neighboring cilia could provide complementary information about ciliary morphology and coordinated beating behavior that CiliaIO does not currently report. Crucially, these and other metrics could be readily integrated using data already captured by CiliaIO. Additionally, PCA-based motility features derived from the entire tracked skeleton over time might yield interpretable morphodynamic descriptors that may offer greater explanatory power than summary statistics such as dominant frequency alone. These measurements have been shown to capture aspects of ciliary dynamics, including axonemal structural integrity and inter-cilia synchrony, that are not accessible through frequency- and single-point skeleton-based metrics alone (Walker et al., 2020).

Finally, we note that the absence of these computations in the current release reflects scope rather than a fundamental constraint of the pipeline. CiliaIO generates structured outputs, including per-frame segmentation masks, tracked skeletons, and bounding box metadata, which are directly compatible with downstream analyses that implement such metrics, meaning that additional descriptors can be incorporated without requiring new imaging data. We view this as a natural opportunity for community-driven development: researchers can either integrate CiliaIO outputs into existing analytical software such as CiliaQ (Hansen et al., 2021) and SpermQ (Hansen et al., 2018) or contribute new computational modules directly to the CiliaIO codebase. We warmly invite open-source collaboration to broaden the use of the analytical repertoire of CiliaIO and look forward to the tool evolving through community engagement over time.

## MATERIALS AND METHODS

### EXPERIMENTAL MODEL AND STUDY PARTICIPANT DETAILS

#### Zebrafish Husbandry

All experiments were performed according to University of Texas at Austin IACUC standards. Wild-type AB strains were used for experiments unless otherwise stated. Embryos were raised at 28.5℃ in egg water (0.15% Instant Ocean in reverse osmosis water) and then transferred to standard system water at 5 dpf. Animals were raised at 28.5℃ under a 14/10 light/dark cycle until the start of experiments. The age of animals used for each experiment is specified in the respective figure legends. Both female and male zebrafish were used for all experiments. As zebrafish do not exhibit sexual dimorphism until the early adult stages of development, sex is not a factor in any experiments described in this manuscript.

### ADDITIONAL RESOURCES

#### Technical description of the *CiliaIO* software

To get CiliaIO and obtain information on the raw data used for this analysis, user guide and source code, please visit the Zenodo page (https://doi.org/10.5281/zenodo.19572444). We also provide a CiliaIO User guide and CiliaIO fine tuning guide in the Supplementary Data. The most up-to-date information will be available on GitHub (https://github.com/chekfung/cilia_io).

### METHOD DETAILS

Supplementary Resources - Table S3

#### WGS / WES analysis

To map a mutation, a single F4 homozygous *stl438* mutant was outcrossed to WIK to generate a mapping cross family. The mapping cross family was intercrossed and F5 progeny were screened for spine defects. 20 phenotypic *stl438* mutant zebrafish were pooled. 20 non-phenotypic wild-type and heterozygous siblings were pooled together. Equimolar concentration of DNA was taken from each pool and submitted to the Genome Technology Access Center (GTAC, Washington School of Medicine) for whole genome sequencing. In house pipeline was used to determine the region of homozygosity segregating with the mutant pool in contrast to WT sibling pool (1 wild type : 2 heterozygous). A SNP subtraction analysis using other whole genome sequencing (WGS) datasets from this screen was used to narrow the number of candidate SNPs.

#### *bbs2^stl438^* genotyping method

For genotyping, adult fish were fin clipped and DNA was extracted. The tissue was lysed in 50 mM NaOH and heated to 95℃ for 20 minutes. After lysing, the DNA sample was neutralized with a 1:4 volume of 1M Tris-HCI, pH 8.0. Samples were immediately used for PCR reaction or were stored at -20℃ until further processing. The *bbs2^stl438^* locus was amplified by PCR with Promega GoTaq Polymerase Enzyme using the following primers: 5’-TTCTCAGCAGACCTCCTCCTCG-3’ and 5’-CGGTTCATTCCACTGTTGCGAC-3’ and the following settings: an initial denaturation at 95℃ for 3 minutes, followed by 3 cycles of denaturation at 95℃ for 30 seconds, annealing at 65℃ for 30 seconds, and extension at 72℃ for 30 seconds, followed by a final extension at 72℃ for 5 minutes. PCR samples were screened on a 3% agarose gel to confirm the successful PCR reaction and sent to Sanger Sequencing. The base substitution was confirmed using electropherograms.

#### RNA Isolation and Cloning

Total RNA was isolated from both wild-type and *bbs2^stl438^* embryo clutches using Trizol based RNA extraction following with cDNA synthesized using reverse transcription enzyme. The reverse transcription protocol followed with high fidelity amplification reaction using the following primers that targets full length *bbs2* coding region: 5’-ATG CTA GTG CCC ATC TTC ACA CTT AAA CTG AAC CAT AAA ATA AAT CC-3’ and 5’ TCA GGA GGA TGT AGT ACCAGC TCT CAT GAT CTT GAA AAG AGC-3’. The PCR samples used for ligation protocol. The ligation reaction was incubated at 4℃ overnight and transferred to ampicillin selective plates. Individual colonies were selected and enriched in selective grow medium. The following day, the plasmids were isolated using a MiniPrep Kit. Plasmids were cut with EcoRI restriction enzyme for sample confirmation and whole-plasmid sequencing was performed by Plasmidsaurus using Oxford Nanopore Technology with custom analysis and annotation.

#### Immunohistochemistry

Larval fish were fixed using a paraformaldehyde solution (4% PFA in 1X PBS). Animals were washed in PBSTr (1X PBS with 0.1 Triton X-100) and after, MilliQ water. The samples were permeabilized by incubating in ice cold acetone at -20℃ for 20 minutes and washed consecutively with MilliQ water following PBSTr. Animals were incubated with Wyart Blocking Buffer (0.5% Triton X-100, 1% DMSO, 10% NGS in 1X PBS) for 2 hours at room temperature. The samples were incubated with the 1:100 anti-zebrafish Arl13b(Duldulao et al., 2009) rabbit primary antibody at 4℃, overnight. The following day, samples were washed with PBSTr and incubated with anti-rabbit antibody for overnight at 4℃. Samples were transferred into glycerol solution and imaged using a Nikon CSU-W1 Yokogawa Confocal Microscope .

#### MicroCT imaging and analysis

2 months post fertilization wild-type and *bbs2^stl438^* fish were euthanized and fixed in 10% neutral buffered formalin overnight. Animals were mounted in 8% agarose for imaging. Bruker SkyScan 1276 micro-computed tomography (MicroCT) machine was used with following settings: 80 kV, 165 mA and10.5-micron resolution. Raw data was reconstructed using the Bruker nRecon software (version 3.0) and reconstructed scans were rotated in the Bruker Data Viewer software (v1.6.0.0). Reconstructed and rotated samples were loaded into 3D Slicer (v5.6.2) (Fedorov et al., 2012) for image processing.

#### Cilia Confocal Microscopy Dataset Collection and Manual Labeling

Larval fish at 4 dpf and 7 dpf were anesthetized in 0.0016% Tricaine and mounted in 2% low melt agarose dissolved in egg water before imaging with the Nikon CSU-W1 Yokogawa Confocal Microscope. Transgenic zebrafish strains *Tg(foxj1:Arl13b-GFP)^ut22^* for wild type and *bbs2^stl438^* mutants were generated as previously described (Konjikusic et al., 2018). To confirm the sampling rate needed to accurately define cilia frequency, we first oversampled at 198.8 Hz at both 4 and 7 dpf and determined that the maximum and average beating frequency of central canal cilia to be ±21 Hz and ±7 Hz, respectively (Video S7). From this, we standardized all imaging of cilia dynamics in the central canal to collect 5 second movies at 49.9 Hz, representing an ∼2.38X oversampling rate, surpassing the Nyquist rate for perfect reconstruction of sampled signals, without sacrificing resolution needed for higher-fps confocal videos. To take cilia images, 5 second videos were acquired at 60X magnification with following settings: 25 fps, 40 ms exposure; 49.9 fps, 20 ms exposure; or 198.8 fps, 5 ms exposure.

#### Fine-Tuning of YOLO for Cilia Identification

*CiliaIO* fine-tunes the Ultralytics YOLOv11m model (Jocher and Qiu, 2024; Redmon et al., 2016) to detect cilia. 32,167 live confocal images of central canal motile cilia were hand-annotated. Contrast and brightness of the confocal images were adjusted using FIJI (Schindelin et al., 2012). 3 wild-type and 4 *bbs2^stl438^* mutant central canal confocal microscopy videos were imaged (25 and 49.9 fps) and labeled frame by frame using Label Studio (Tkachenko et al., 2020) to assist with annotation of bounding boxes. An 80/20 video-wise train-validation split was performed after manual labeling of the bounding boxes, consisting of 918 training frames (19,087 cilia), and 474 validation frames (13,394 cilia). Additionally, a separate test set consisting of 53 frames (1507 cilia) from an unseen confocal microscopy video was used for evaluating final performance of the fine-tuned model. By splitting the dataset at the video level, dataset leakage is minimized, ensuring that the validation and test set only consists of complete microscopy videos not present in training. For the stratified 3-fold cross validation, we keep the same test set, but randomly permute the other 6 videos with the constraint that the validation set contain 1 wild type and 1 mutant animal. The full dataset details are provided in Table S1. The dataset size was increased using mosaic augmentation, which combines four separate images into one. For each mosaic tile, there is a 50% probability for horizontal flipping, up to 50% random scaling, and up to 10% image translation. Additionally, each image had a 40% chance for random erasing, meaning that a randomly selected region in the training image is replaced with a constant value, effectively increasing robustness by forcing the network not to rely on single regions for accuracy.

All training runs were performed on an x86 machine (AMD 5800x3D, GTX 1080Ti, 32GB DDR4), in Linux Ubuntu 22.04.05 using PyTorch 2.2.2 and Ultralytics 8.3.1. Fine-tuning of the Ultralytics YoloV11m model (Jocher and Qiu, 2024; Redmon et al., 2016), with 20.1 million trainable parameters, was performed using the previously mentioned dataset for 300 epochs. The input images were resized to 640x640 pixels, with a mini-batch size of 4. The network was optimized using stochastic gradient descent with a momentum of 0.937, weight decay of 0.0005, and an initial learning rate of 0.01 with a warmup time of 3 epochs. The loss function used the typical box, classification, and distribution focal loss scaling of 7.5, 0.5, and 1.5, respectively. Overall training took 7.49 hours. On the test set of 1,507 cilia, the fine-tuned YOLO model achieved an F1 score of 86% with a confidence threshold of 0.63 and a mean average precision of 91% with a 0.5 overlap.

#### CiliaIO Pipeline

In this section, each step of the *CiliaIO* pipeline is described in detail. From a raw video, *CiliaIO* computes the following on each frame: 1) YOLO-based cilia identification, 2) Augmented ByteTrack for tracking of cilia IDs between frames, 3) SAM-based cilia segmentation, 4) skeletonization via morphological thinning, and a 5) spline curve fit of the skeleton. To preprocess the raw file, provided a .tif file, *CiliaIO* reads the metadata of the file to get the pixels to micron conversion rate and fps. It then reads through every frame of the .tiff file using the *tifffile* Python package (Gohlke, 2025), normalizes the pixel intensities of each frame to 0 to 255 using *OpenCV* (Bradski, 2000), and converts the image to RGB. The resulting frames are saved as an .mp4 file that is then opened.

For each frame of the video, **1)** the fine-tuned *Ultralytics* YOLOv11m model tracks the frame using the native track function with the tracker set to the “bytetrack.yaml” file, and a matching confidence of 0.4. The inference call on the frame returns the x1, y1, x2, y2 (xyxy) coordinates of the detected cilia bounding boxes along with their IDs. **2)** If this is the first frame, each box is reassigned an ID from 0 to N cilia. If this frame is not the first frame, the cilia bounding box centers are compared against the historical centers, where the new frame’s bounding boxes are greedily matched based on which historical bounding box center has the minimum distance to the new frame’s bounding box. If the bounding box center is beyond the max matching distance (30 pixels or 3.21 microns for our analysis), it is assigned a new ID. If a cilia ID is not matched to a new frame’s bounding box, the historical box and ID are added into the valid bounding boxes for up to MAX_AGE consecutive missing frames (MAX_AGE=5 for our analysis). After that, it is removed from the tracking dictionary and is no longer tracked. This hyperparameter is akin to the patience of *CiliaIO*. To get rid of duplicate IDs that have identified the same cilia, boxes are coalesced with an Intersection over Union (IOU) threshold greater than 50% into a single ID. 3) Using the *segment_anything* (Kirillov et al., 2023) Python package with the out-of-the-box VIT_H_4b8939 SAM checkpoint, the frame is loaded, and the image embedding is computed using the .set_image function. Afterwards, all bounding boxes in the frame are converted into a (N, 4) tensor, where the columns represent the x1, y1, x2, and y2 values of each bounding box.

Using transform.apply_boxes_torch for the SAM model, the input box tensor is converted into a prompt embedding. SAM is run with the transformed boxes embedding and a single mask output. To address cases where the mask contains multiple disconnected regions, the largest connected component is identified and retained for subsequent processing. 4) To get the skeleton, the mask is dilated by 5 pixels on each side to stretch more of the skeleton to the edges of the true mask. The padded mask is skeletonized using *scikit-image*’s (Walt et al., 2014) skeletonize function, which performs morphological thinning on the binary mask. This identifies the center pixels, providing the centerline coordinates in order. Since skeletonization stops short of the actual end of the cilia mask, the skeleton is grown at each end by applying the Sobel operator to the distance transform of the mask. The Sobel filter computes the spatial gradient of the distance field, which produces vectors that point from the interior toward the nearest mask boundary. The skeleton is grown by following the minimum gradient of the neighboring pixels until the mask boundary is reached. The skeleton is finally cleaned up by removing sharp turns between two adjacent points (greater than 120 degrees) from one end of the skeleton to the other end. For each pair of connected points in the skeleton, any point where the local angle exceeds a specified threshold is removed to eliminate sharp or spurious branches. 5) The pixel-based skeleton is fit to an N-point (100 points in our case) B-spline to smooth out the skeleton. At each frame, the morphology metrics like length, straightness, and area are computed. Otherwise, the center point of the skeleton is tracked per cilia ID across the entire video, and then the dynamics are calculated for each of the unique cilia IDs in our quantification engine if the number of consecutive frames of the cilia ID appearing is greater than a user-set minimum number (42 in our case). All tuned hyperparameters can be found in Table S2.

#### Pixel-based FFT method parameters

The scripts are adapted from the available MATLAB resources of Thouvenin et al., 2021 and adjusted to Python. Local average filter of size 4 was applied and the first maximal peak of the Fourier spectrum is extracted. Regions of interest containing more than 40 pixels were segmented and analyzed. In cases where there are multiple frequencies in a single cilium’s path, the frequency from the largest region of interest was recorded.

### QUANTIFICATION AND STATISTICAL ANALYSIS

Statistical analysis was performed using Python 3.10, with packages: statsmodel 0.14.2 (Seabold and Perktold, 2010), scikit-image 0.25.2(Walt et al., 2014), scikit-learn 1.5.0 (Pedregosa et al., 2011) and Scikit-bio 0.7.2(Aton et al., 2026). Statistical details are given in figure legends.

#### Quantification of Morphodynamic Metrics

The quantification of each cilium morphology follows after each frame goes through the ML pipeline. The skeleton of the cilium is used to quantify cilia length and straightness (ratio of cilium length to the minimum distance between the basal body and tip), while the segmentation mask is used for the area. Motility metrics are similarly obtained after every frame. A point is chosen on the skeleton as a user-set parameter and is used as the tracked point of interest over time at which motility metrics will be computed. The middle point of the skeleton was chosen for cilia motility quantification in our studies since it minimizes the error that may come from the tip. After obtaining the time-domain signal, we performed baseline correction using symmetric least squares smoothing (Eilers and Boelens, 2005) to remove slow drifts and background trends. Metrics such as path length, average speed of the cilia, mean angular velocity, and cumulative angular velocity are measured from the tracked point of interest. The point cloud of the tracked point of interest is then quantified using PCA to get the major and minor axes of variation of the cilia movement. Examining the major axis of movement provides a 1D time-series signal to evaluate amplitude and frequency. Amplitude is evaluated directly from the time domain. For frequency, the major axis of movement over time is converted into a power spectrum using the Fourier transform. To obtain the power spectrum, the detrended time domain signal is passed through a Hamming window (Hamming, 1977) of the same length for spectral smoothing, and then zero padded by a factor of 4x to increase the fidelity of the discrete Fourier transform. From the power spectrum, the dominant frequency and centroid frequency are extracted. Spectral entropy, which measures the disorder or complexity of a signal is described in Eqn 1. *S*(*f_i_*) is the power spectrum which is obtained from the squared magnitude of each Fourier coefficient. N represent the number of discrete Fourier coefficient bins in the analysis. N is also associated with the sample size of the time domain signal.

*Equation 1*.

Normalized entropy

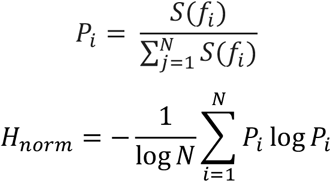

#### Definition and Quantification of Different Morphodynamic Metrics

1. **Mean Cilia Length:** The averaged length of a cilium, in microns, across all frames linked to that cilia ID.
2. **Mean Cilia Area:** The averaged area of a cilium, in microns, across all frames linked to that cilia ID.
3. **Mean Straightness:** Straightness is calculated as a dimensionless ratio between the arc length of a cilium versus the minimum distance between the two ends of the skeleton, with the arc length in the denominator. Straightness is calculated at every identified frame for a cilia ID and then averaged together.
4. **Std Straightness:** Straightness calculated at every identified frame for a cilia ID, but the standard deviation is computed with 1 degree of freedom.
5. **Max Straightness:** The maximum straightness, calculated from the set straightness calculated from the identified frames for a cilia ID.
6. **Min Straightness:** The minimum straightness, calculated from the set of straightness calculated from the identified frames for a cilia ID. The following metrics are motility-based and track one point (center) of each cilium skeleton:
7. **Total Traveled Length (Path Length):** Total length, in microns, of the entire path across all consecutive tracked frames.
8. **Average Speed:** The speed, in microns/second, is calculated distance between each consecutive pair of time steps in the tracked point and divided by the sample interval of the microscope and averaged across all consecutive frames.
9. **Mean Angular Velocity:** The angular velocity, in radians / second, is calculated by taking the arctan between each consecutive pair of timestep paths across all consecutive frames and then averaged across all frames.
10. **Cumulative Angular Displacement:** The absolute value of the sum of the change in angle of the consecutive pairs of timestep paths across all consecutive frames.
11. **Semi Major Axis Length:** The square root of the largest eigenvalue of the covariance matrix taken from the point cloud of cilia movement across all consecutive frames. It represents the length of the spread, in microns, of the variance in the major axis from PCA.
12. **Semi Minor Axis Length:** The square root of the second largest eigenvalue of the covariance matrix taken from the point cloud of cilia movement across all consecutive frames. It represents the length of the spread, in microns, of the variance in the minor axis from PCA.
13. **Radius of Gyration:** Measures the root mean squared (RMS) distance, in microns, of the point cloud movement from the center of mass of the point cloud. The point cloud is the tracked point on the skeleton across all consecutive frames.
14. **Mean Amplitude:** The mean of the absolute deviation of a tracked point on the skeleton across all consecutive frames from the PCA major axis origin, in microns.
15. **Max Amplitude:** The maximum of the absolute deviation of a tracked point on the skeleton across all consecutive frames from the PCA major axis origin, in microns.
16. **RMS Amplitude:** The root mean squared absolute deviation of a tracked point on the skeleton across all consecutive frames from the PCA major axis origin, in microns.
17. **Dominant Frequency:** The dominant frequency, in Hz, is derived from the frequency with the maximum magnitude power in the Fourier transform of the tracked point on the skeleton movement along the PCA major axis.
18. **Centroid Frequency:** The centroid frequency, in Hz, is derived from the center of mass of the power spectrum in the Fourier transform of the tracked point on the skeleton movement along the PCA major axis.
19. **Entropy:** Spectral entropy of the power spectrum computed from the Fourier transform of the tracked point on the skeleton movement along the PCA major axis (Eqn 1).

## ACKNOWLEDGEMENTS

We thank Brian Perkins for sharing the *bbs2^Iri82^* line and Zhaoxia Sun for sharing the zebrafish specific Arl13b antibody. We thank members of the Gray and Wallingford Lab for help with fish husbandry and helpful discussion.

## AUTHOR CONTRIBUTIONS

A. conducted all the wet lab experiments, generated all the figures, and wrote the original draft of the manuscript. J.H. conducted *in-silico* design of the ML program, performed statistical analysis, and wrote the original draft of the manuscript. T.G.B. and I.B.J assisted E.A. in crossing, genotyping, and phenotyping. They also assisted J.H. in generating the ML dataset for downstream applications. B.Z. preformed the genetic mapping of the *stl438* zebrafish mutant found by ENU screen by R.S.G. in the L.S.K. lab. A.G. provided ML computational resources and reviewed/edited the manuscript. J.B.W. supervised the experiment and reviewed/edited the manuscript. R.S.G. provided initial conceptualization for the project and supervised all experiments and reviewed/edited the manuscript.

## SUPPLEMENTARY FIGURE LEGENDS

**Figure S1.**
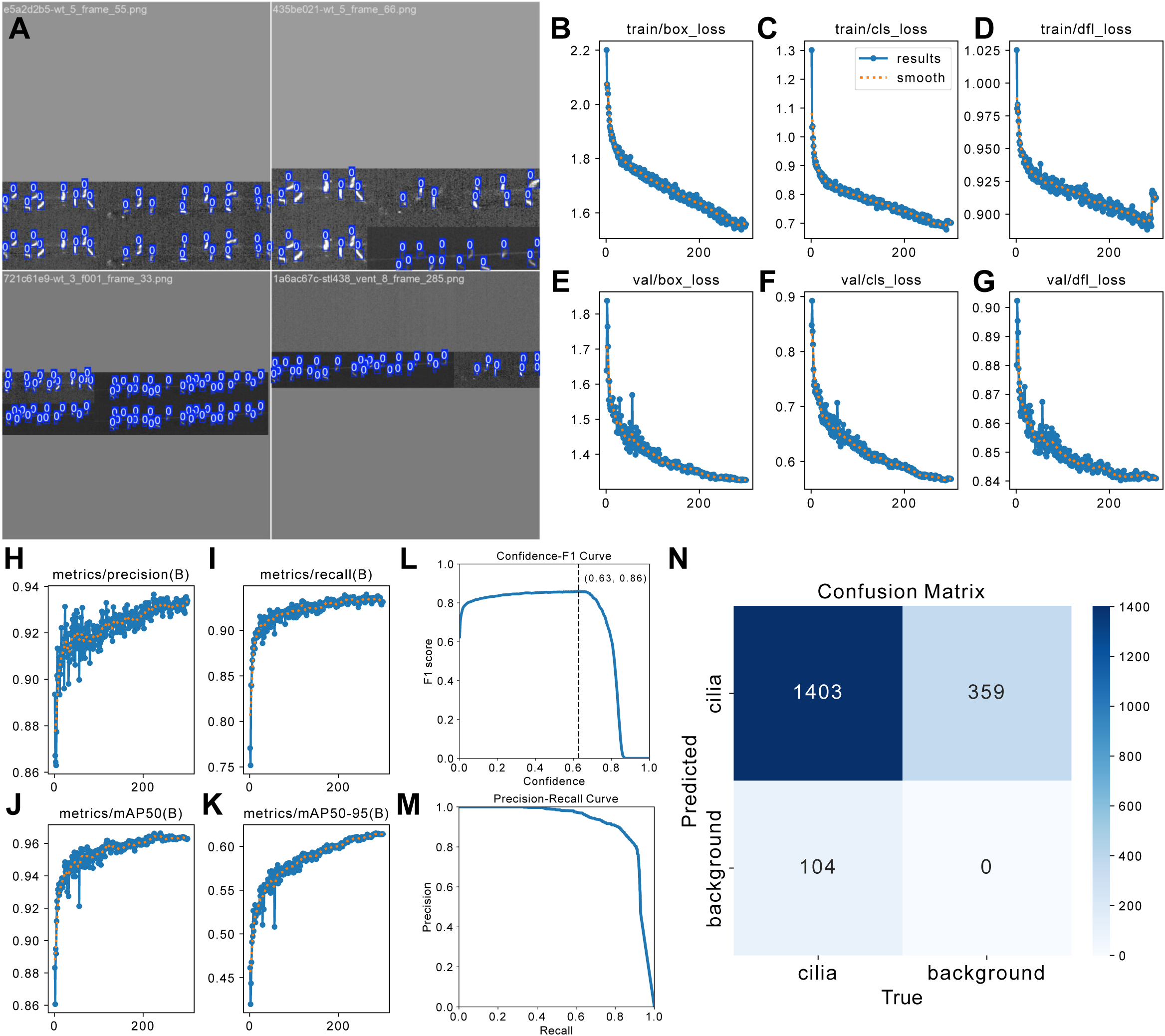
YOLOv11m fine-tuning performance and training metrics, related to Figure 1. Example mosaic augmentation of dataset stitching 4 separate training frames together with augmented scaling, flipping and translation. **(B)** Training box loss over 300 training epochs, **(C)** Training classification loss over 300 training epochs, **(D)** Training distribution focal loss over 300 training epochs, **(E)** Validation box loss over 300 training epochs, **(F)** Validation classification loss over 300 training epochs, **(G)** Validation distribution focal loss over 300 training epochs, **(H)** Cilia object classification precision on validation set over 300 training epochs, **(I)** Cilia object classification recall on validation set over 300 training epochs, **(J)** Validation mAP50 for cilia: mean average precision with intersection over union of 50% (predicted bounding box overlaps at least 50% area with the hand labeled bounding box) over 300 training epochs, **(K)** Validation mAP50-95 for cilia: mean average precision with intersection over union greater than 50% up to 95% area over 300 training epochs **(L)** Test F1-Confidence curve generated with the best-performing model saved at epoch 239. The F1 score is 86% with at 0.63 confidence. **(M)** Test precision-recall curve generated with the best performing model saved at epoch 239. The mAP50 is 91%. **(N)** Confusion matrix on the cilia test set generated with the best-performing model saved at epoch 239 with confidence threshold of 0.25 showing true positives (cilia correctly detected), false negatives (cilia missed), and false positives (background regions incorrectly classified as cilia).

**Figure S2.**
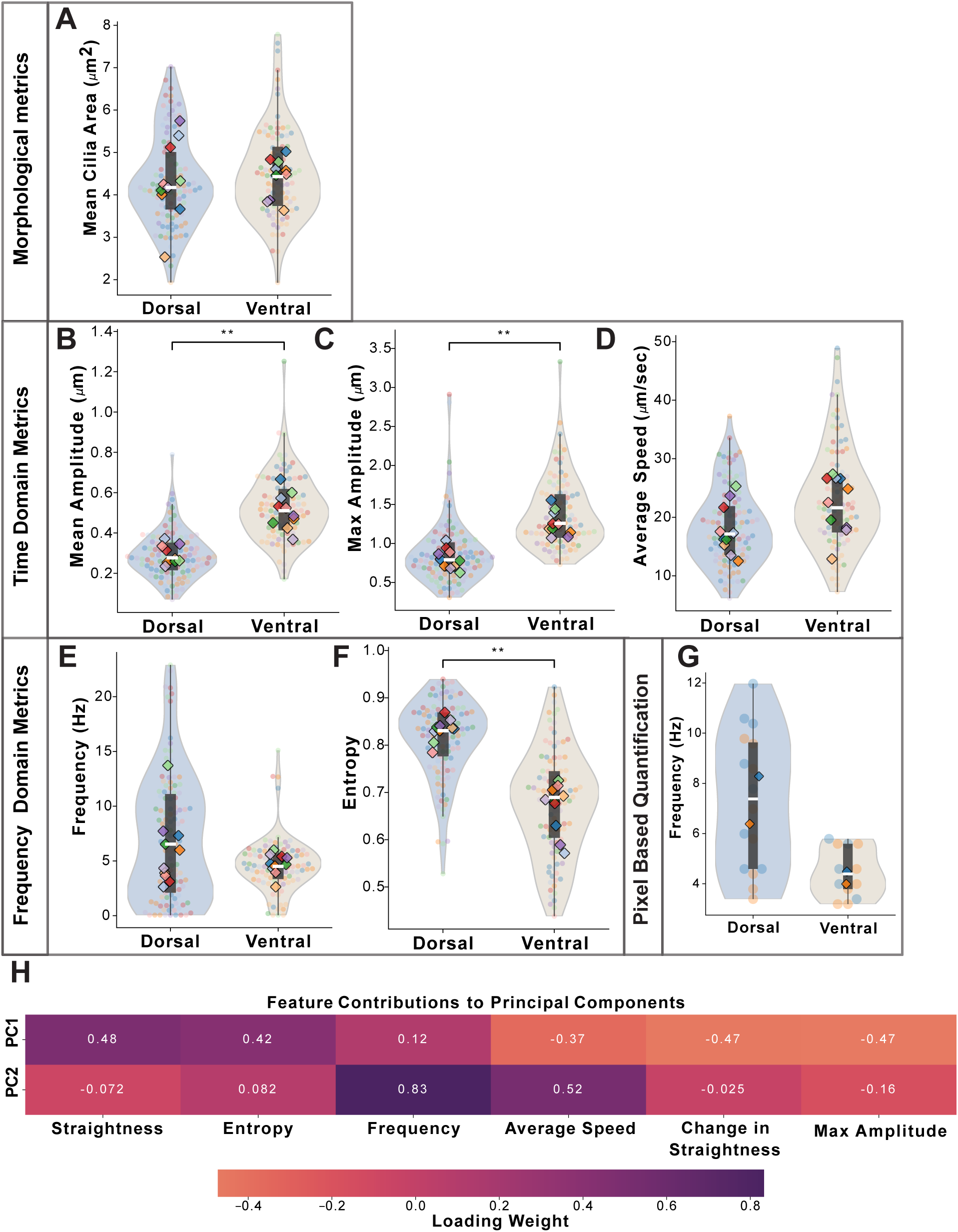
Wild-type dorsal and ventral cilia exhibit differences in beating dynamics while maintaining similar static morphological features, related to Figure 5. **(A–G)** CiliaIO Violin SuperPlots comparing dorsal (blue) and ventral (beige) cilia; individual data points are color-coded by animal. (A) No significant differences in mean cilia area (µm^2^) between dorsal and ventral populations. (B, C) Dorsal cilia show significantly smaller (B) mean and (C) maximum beating amplitude (µm). No significant differences in (D) average speed (µm/s) and (E) dominant frequency (Hz) between dorsal and ventral cilia populations. Dorsal cilia display higher (F) spectral entropy. Wild-type: 10 animals comprising 117 dorsal and 102 ventral cilia; ≥ 3 independent replicates. (G) Pixel-based FFT analysis (Thouvenin et al., 2021) showing a similar dorsal–ventral frequency trend as observed with CiliaIO; analysis from 2 wild-type animals comprising 30 cilia**. (H)** PCA component coefficient heatmap derived from wild-type cilia showing the contribution of each morphodynamic feature to PC1 and PC2 for all 2D PCA projections. For (A-G), paired Wilcoxon signed-rank test was used to compare the median values of dorsal and ventral cilia within each fish. Violin box represents interquartile range; white line indicates median. p-value: ** ≤ 0.01.

**Figure S3.**
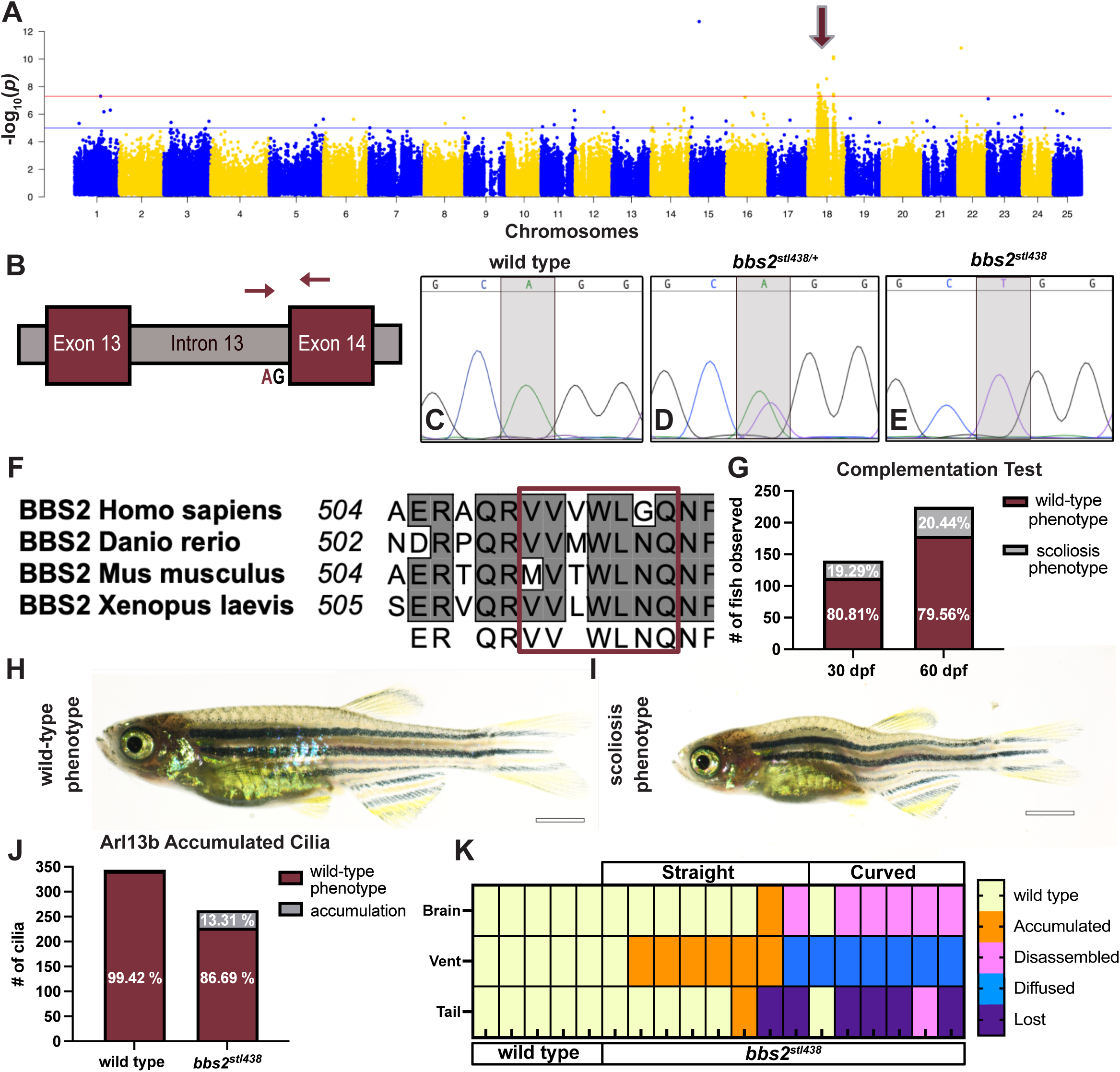
The *stl438* allele harbors a single-nucleotide variant at the conserved splice acceptor site of intron 13 in *bbs2*, related to Figure 6. **(A)** Manhattan plot of −log₁₀(*p*) from SNP-based linkage and homozygosity mapping using whole-exome sequencing data from *stl438* mutants (n = 20) and wild-type siblings (n = 20). Horizontal lines denote suggestive (blue) and genome-wide (red) significance thresholds. Arrow indicates the peak region of homozygosity on chromosome 18. **(B)** Schematic of the *stl438* mutation site in *bbs2*, showing the A>T transversion that disrupts the conserved splice acceptor site of intron 13 (red). Red arrows indicate primer positions used to amplify the intron–exon junction. **(C–E)** Representative Sanger sequencing chromatograms from (C) wild-type (n = 8/8), (D) *bbs2^stl438/+^* heterozygous (n = 8/8), and (E) *bbs2^stl438/stl438^* homozygous animals (n = 14/14) confirming the A>T substitution at the intron 13 splice acceptor locus. **(F)** Clustal alignment of the Bbs2 platform domain across species; the predicted deletion site is highlighted with a red box. **(G)** Bar graph showing phenotypic counts from combined complementation tests at 30 dpf and 60 dpf (30 dpf, n = 140; 60 dpf, n = 225). **(H, I)** Brightfield images of 60 dpf fish from a complementation cross (*stl438/+* × *bbs2^iri82/+^*) showing (H) wild-type body axis and (I) scoliosis in *trans*-heterozygous *bbs2^iri82/stl438^* animals. Brightfield images stitched from multiple panels. Scale bar: 2 mm. **(J)** Bar graph showing Arl13b axonemal accumulation frequency in wild-type and *bbs2^stl438^* cilia at 4 dpf (Fisher’s exact test; p ≤ 0.0001). Wild-type: 15 animals, 344 cilia; *bbs2^stl438^* : 11 animals, 263 cilia. **(K)** Categorization of Reissner fiber phenotypes in the brain, vent region, and tail region of wild-type and *bbs2^stl438^* animals at 4 dpf. Each column represents one animal; each row represents one anatomical region. Analysis from 5 wild-type and 14 *bbs2^stl438^* animals; ≥ 3 independent replicates.

**Figure S4.**
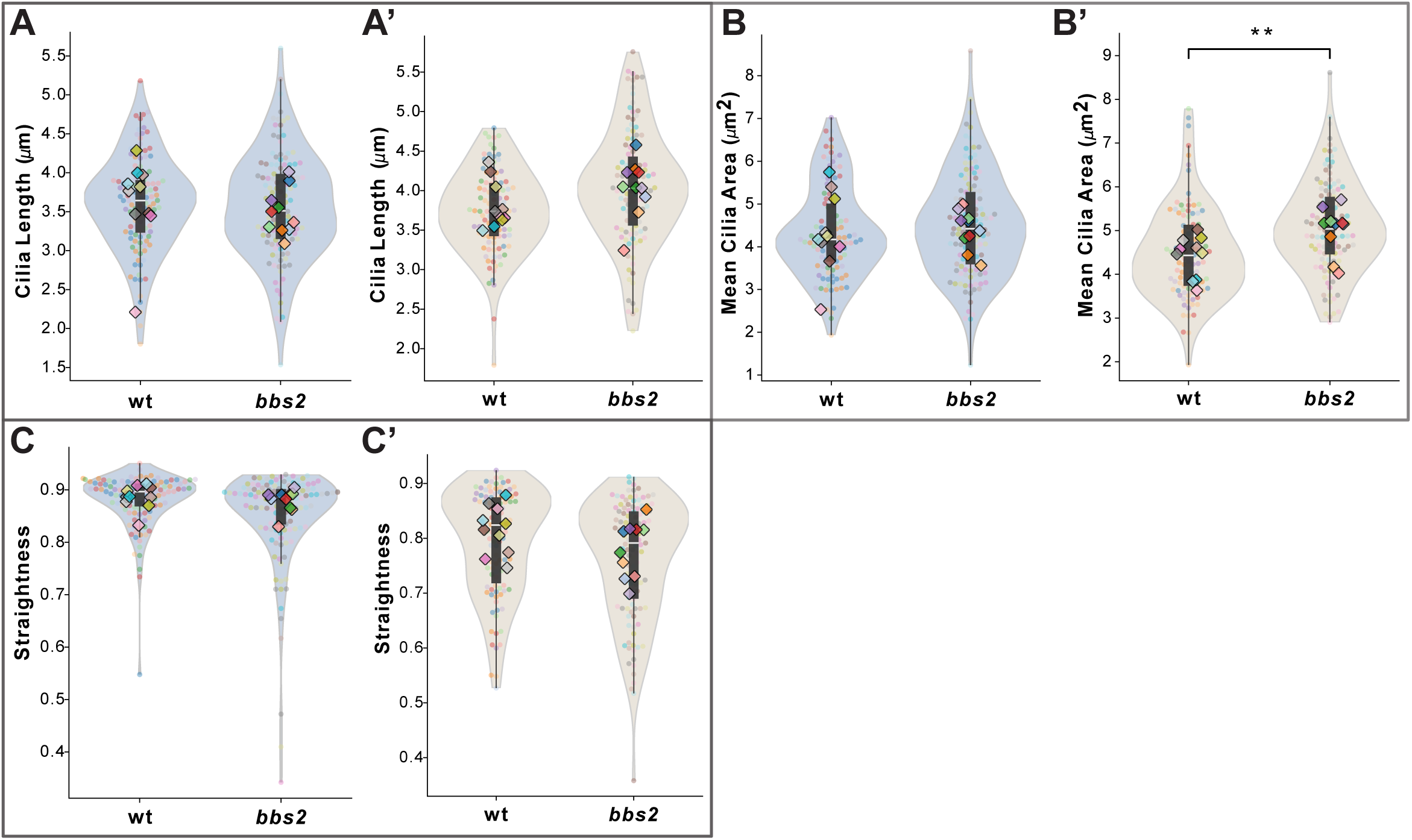
*bbs2^stl438^* mutants show altered dynamic morphological metrics in dorsal cilia with largely unaffected ventral cilia, related to Figure 7. **(A–C′)** CiliaIO Violin SuperPlots comparing wild-type and *bbs2^stl438^* animals within dorsal and ventral populations; individual data points are color-coded by animal. Mean cilia length (µm) displayed no changes in both dorsal (A) and ventral cilia (A’) in *bbs2^stl438^*. Mean cilia area (µm²) is increased only in ventral cilia (B′), with no change in dorsal cilia (B). Mean axonemal straightness metrics (C, C’) displayed no changes in *bbs2^stl438^* animals. Mann-Whitney U tests performed between wild-type and *bbs2^stl438^* medians for each fish within each dorsoventral group. Violin box represents interquartile range; white line indicates median. p-value:** ≤ 0.01. Wild type: 10 animals, 117 dorsal and 102 ventral cilia. *bbs2^stl438^* : 10 animals, 132 dorsal and 117 ventral cilia. ≥ 3 independent replicates.

**Figure S5.**
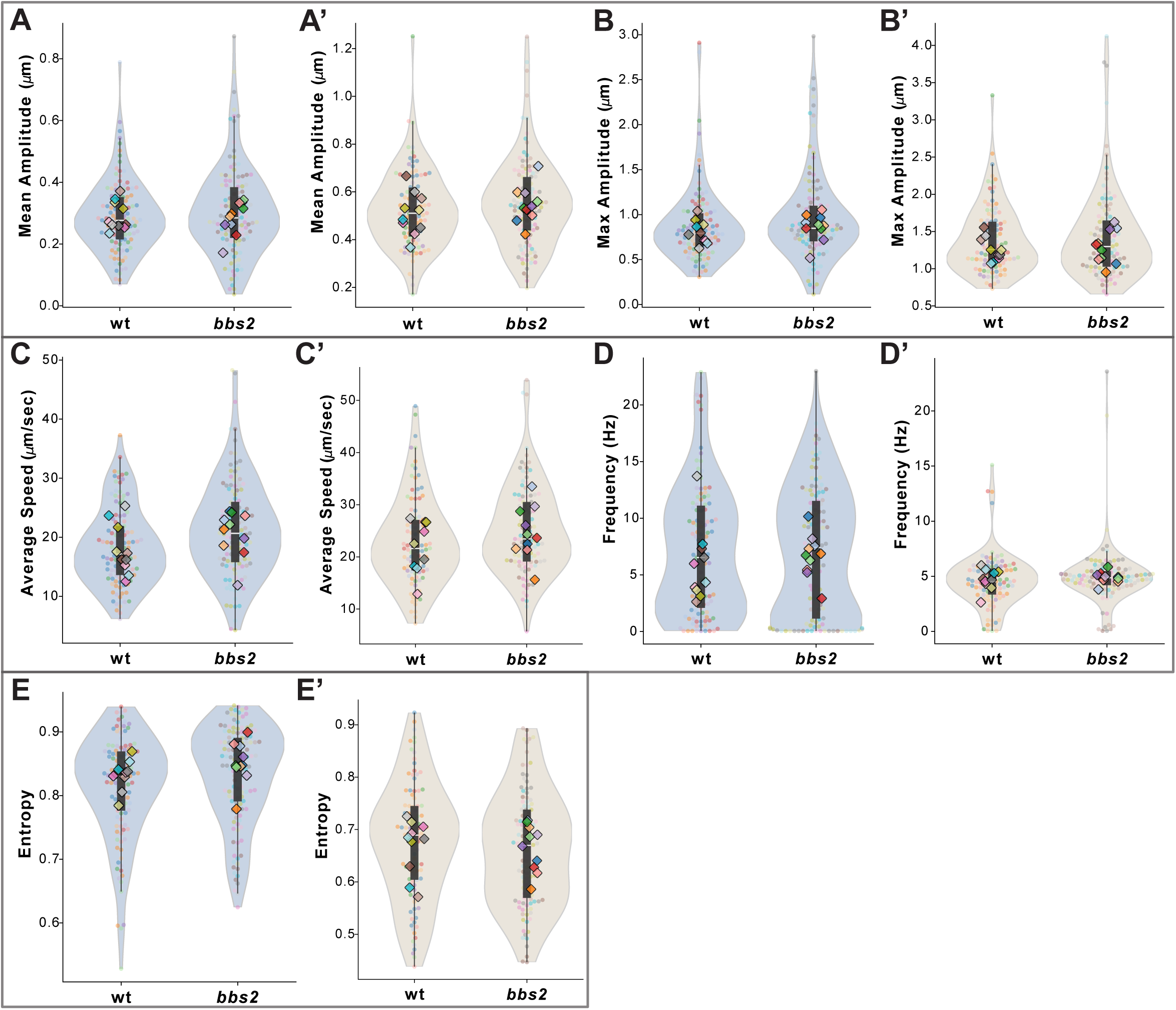
*bbs2^stl438^* mutants display selective changes in dorsal cilia dynamic metrics, related to Figure 7. **(A–E′)** CiliaIO Violin SuperPlots comparing wild-type and *bbs2^stl438^* animals within dorsal and ventral populations; individual data points are color-coded by animal. Mean (A, A′), maximum (B, B′) beating amplitude (µm), average speed (µm/sec) (C, C’) are comparable between genotypes in both dorsal and ventral cilia. Dominant beat frequency is comparable between genotypes in both dorsal (D) and ventral (D′) cilia. Spectral entropy is unaffected in both dorsal (E) and ventral (E′) cilia. Mann-Whitney U tests performed between wild-type and *bbs2^stl438^* medians for each fish within each dorsoventral group. Violin box represents interquartile range; white line indicates median. p-values: * ≤ 0.05. Wild-type: 10 animals, 117 dorsal and 102 ventral cilia. *bbs2^stl438^*: 10 animals, 132 dorsal and 117 ventral cilia. ≥ 3 independent replicates.

## SUPPLEMENTARY TABLES

**Table S1.** Dataset for fine tuning YOLO.

| **Genotype** | **FPS** | **Frames** | **Total Cilia** | **Max Cilia / Frame** |
| --- | --- | --- | --- | --- |
| WT | 25 | 373 | 11573 | 35 |
| bbs2 | 25 | 314 | 6152 | 24 |
| bbs2 | 25 | 374 | 7634 | 25 |
| WT | 49.9 | 100 | 2640 | 29 |
| WT | 49.9 | 130 | 2661 | 23 |
| bbs2 | 49.9 | 101 | 1821 | 22 |
| bbs2 (test) | 49.9 | 53 | 1507 | 32 |

**Table S2.** CiliaIO Hyperparameters.

| Parameter Name | Value | Description |
| --- | --- | --- |
| DORSAL_VENTRAL_THRESHOLD_GLOBAL | Video Dependent<br>(Default = 80) | Vertical coordinate (y-axis) defining the anatomical boundary for dorsal and ventral categorization within the frame. |
| NUMBER_INTERPOLATION_POINTS | 100 | Spline resolution; defines the number of nodes used to smooth and standardize the raw pixel-based skeleton. |
| MIN_NUMBER_POINTS_TO_EXAMINE_MOTILITY | 42 | Minimum temporal window (consecutive frames) required for a cilium to calculate motility metrics. |
| MATCHING_CONF | 0.4 | Confidence score threshold for the YOLO detector; filters low-probability candidates prior to segmentation mask generation. |
| MAX_DIST | 30 | Distance in pixels defining the maximum allowable centroid displacement of a bounding box for identity preservation between consecutive frames. |
| MAX_AGE | 5 | Maximum occlusion tolerance; specifies the temporal window (in frames) for maintaining object identity during periods of signal loss before track termination. |
| POST_IOU_THRESH | 0.5 | Spatial overlap threshold used to coalesce redundant bounding boxes; prevents over-segmentation by merging detections with a high Intersection over Union (IoU) ratio. |

**Table S3.**
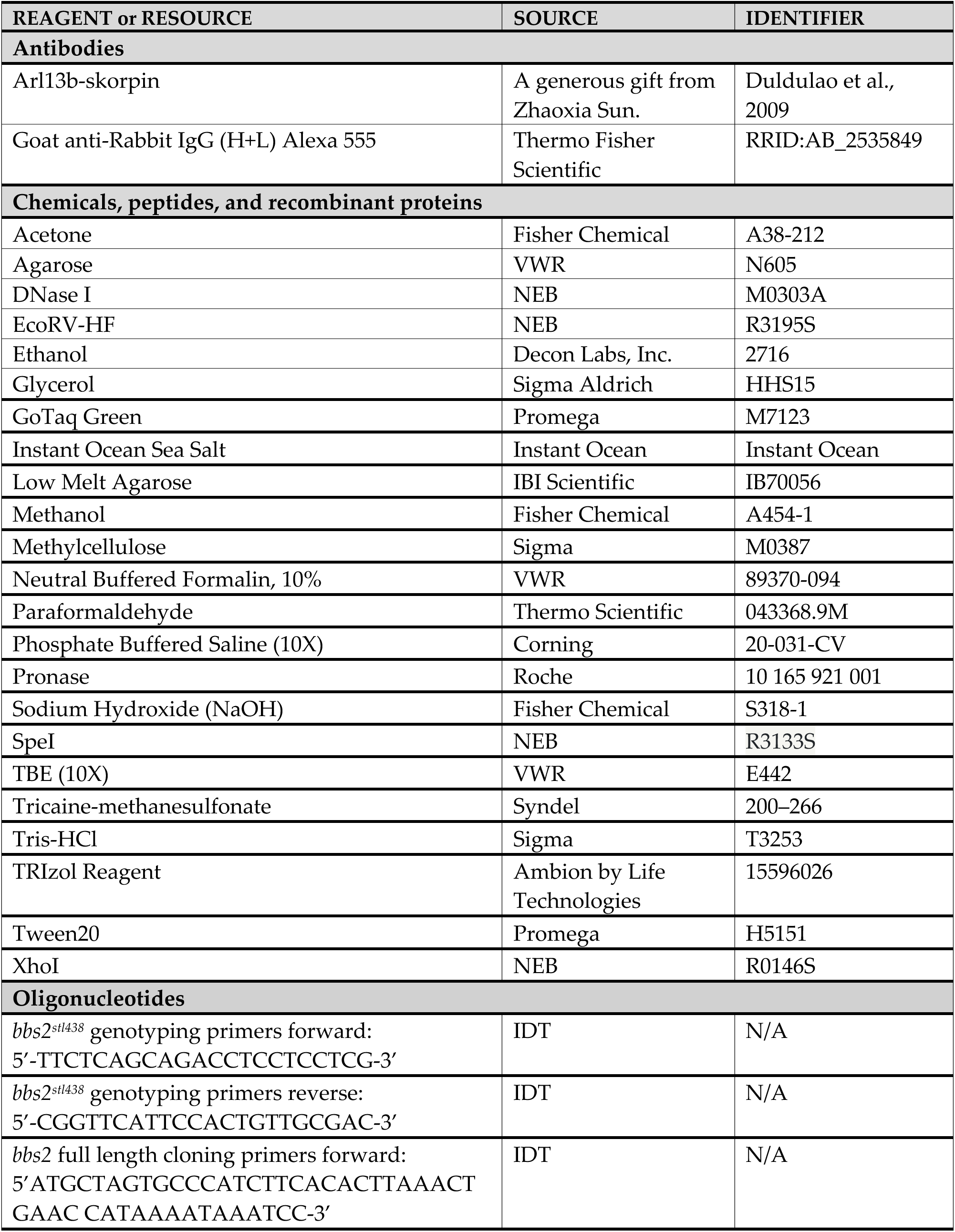
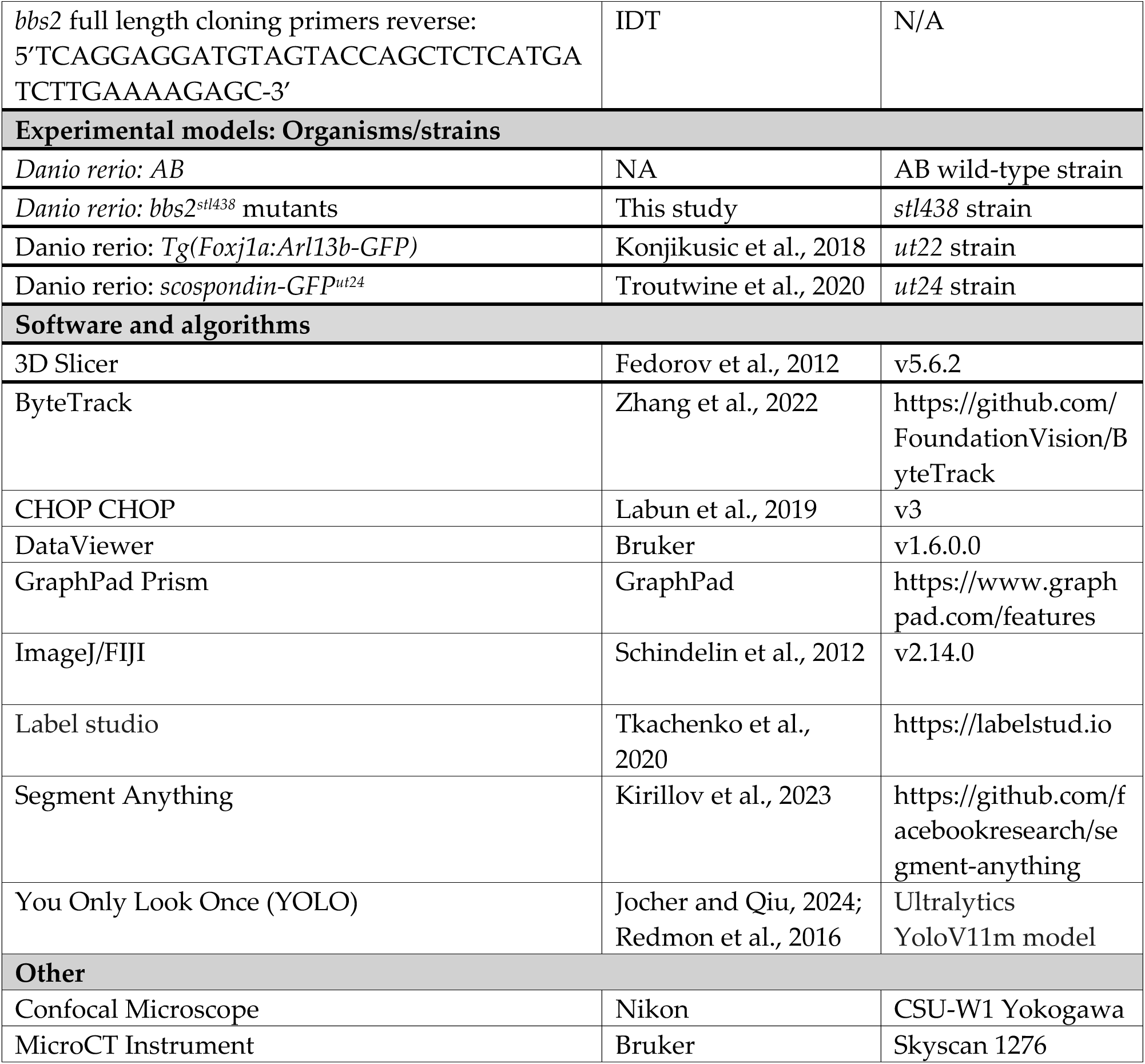
Key Resource Table.

## SUPPLEMENTARY VIDEO LEGENDS

**Video S1:** CiliaIO analysis of a *Tg(Foxj1:Arl13b-GFP)*^ut22^ wild-type, 4 dpf animal. Related to Figure 2. Dorsal cilia are marked with blue and ventral cilia are marked with beige masks, accompanied with cilia IDs. In gray, the YOLO bounding box is displayed. The white line tracks the trajectory for each cilia’s midpoint over time. The video is played at 5 fps.

**Video S2:** Representative live confocal video of Foxj1a+ central canal cilia at the vent region of a transgenic *Tg(Foxj1:Arl13b-GFP)*^ut22^ wild-type, 4 dpf animal. Related to Figure 2. Scale bar = 5 mm. Time stamp is seconds:milliseconds. Video was taken and played at 49.9 fps.

**Video S3:** Representative live confocal video of Foxj1a+ central canal cilia at the vent region of a transgenic *Tg(Foxj1:Arl13b-GFP)*^ut22^ wild-type, 4 dpf animal. Related to Figure 4. Scale bar = 5 mm. Time stamp is seconds:milliseconds. Video was taken and played at 49.9 fps.

**Video S4:** CiliaIO analysis of a *Tg(Foxj1:Arl13b-GFP)*^ut22^ wild-type, 4 dpf animal. Related to Figure 4. Dorsal cilia are marked with blue and ventral cilia is marked with beige masks, accompanied with cilia IDs. In gray, the YOLO bounding box is displayed. The white line tracks the trajectory for each cilia’s midpoint over time. The video is played at 5 fps.

**Video S5:** Representative live confocal video of Foxj1a+ central canal cilia at the vent region of a transgenic *Tg(Foxj1:Arl13b-GFP)*^ut22^ wild-type, 4 dpf animal. Related to Figure 4. Scale bar = 5 mm. Time stamp is seconds:milliseconds. Video was taken and played at 49.9 fps.

**Video S6:** CiliaIO analysis of a *Tg(Foxj1:Arl13b-GFP)*^ut22^ wild-type, 4 dpf animal. Related to Figure 4. Dorsal cilia are marked with blue and ventral cilia is marked with beige masks, accompanied with cilia IDs. In gray, the YOLO bounding box is displayed. The white line tracks the trajectory for each cilia’s midpoint over time. The video is played at 5 fps.

**Video S7:** High speed confocal imaging of a *Tg(Foxj1:Arl13b-GFP)*^ut22^ wild-type, 4 dpf animal. Related to Methods. Scale bar = 5 mm. Time stamp is seconds:milliseconds. Image was taken with 198 fps and played at 20 fps.

## Supplementary Materials and Methods - CiliaIO Fine Tuning Guide

### CiliaIO User Guide

CiliaIO is a state-of-the-art ML-based quantification methodology for cilia morphodynamics. This repository contains a complete multi-modal pipeline for quantifying ciliary motion from confocal microscopy videos. This workflow relies on single-shot cilia detection using a fine-tuned YOLO v11 model which identifies bounding box regions of interest (ROI) to pass to Meta’s Segment Anything (SAM) vision transformer models, removing the need for computationally expensive iterative masking and tracking. Unlike prior work which use classical computer vision approaches like binary thresholding and Canny detection, CiliaIO identifies where cilia are, enabling high quality masks even when cilia overlap. The goal of this project is to enable fast, robust, automated analysis of cilia dynamics overtime across diverse experimental conditions. CiliaIO can detect subtle changes in cilia morphodynamics that may not be detected by the human eye, but are pivotal to developing our understanding of ciliary function overall.

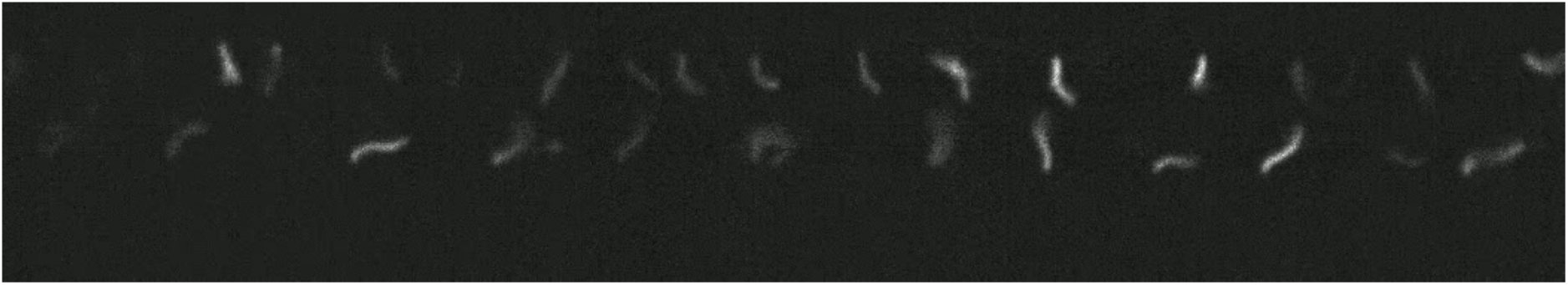

**Raw Frame:** 4-day-post-fertilization Zebrafish mono-cilia raw confocal image.

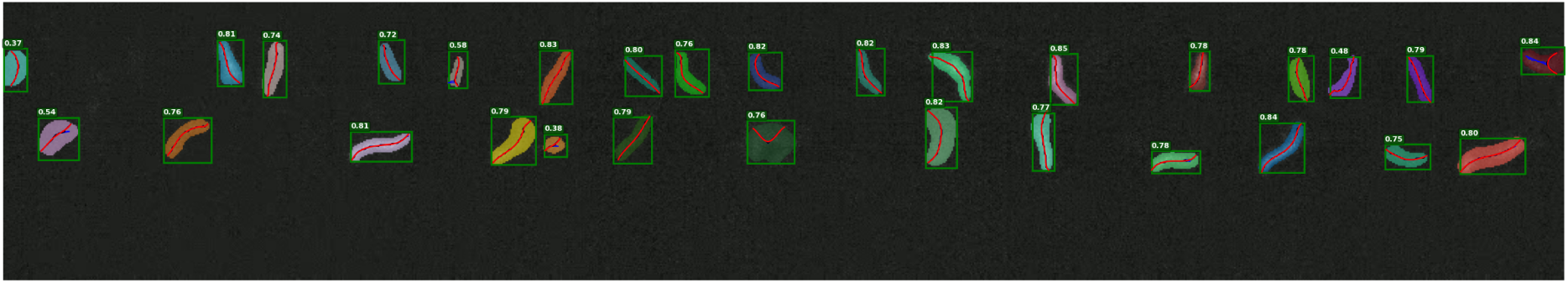

**CiliaIO Frame:** High quality detection, segmentation, skeletonization, and quantification of cilia.

### CiliaIO’s Core Pipeline

1. **Detection** Cilia are detected frame-by-frame using a fine-tuned YOLOv11m model from Ultralytics.
2. **Tracking** Object tracks are initialized and maintained using ByteTrack with additional logic for center-distance matching and overlap-based ID merging. Fallback heuristics handle missed detections and ensure temporal ID consistency across frames.
3. **Segmentation** Boxes are segmented using the Segment-Anything Model (SAM).
4. **Skeletonization** Segmentation masks are skeletonized using the morphological thinning algorithm, which iteratively removes pixels on each side of the mask until one pixel remains.
5. **Quantification** Tracked trajectories are analyzed to extract per-cilium motion features, including spatial amplitude, beating frequency, centroid frequency, and more. Summary statistics are aggregated across videos for downstream analysis.
6. **Visualization** The pipeline supports trajectory overlays, segmentation mask overlays on the raw confocal videos, skeleton tracking, and bounding box tracking.

### Directory Structure

The following text details the directory structure of all files found in CiliaIO. These are not instructions, but instead serve as a reference for every single file included in CiliaIO.

- **data/**: This is where the data goes
- **sam/**: If coming from github, download the SAM ViT-H checkpoint 4b8939.pth file from Meta. See the section labeled: Downloading SAM for GitHub for more details.
- **src/**: Contains all of the source files pertaining to CiliaIO’

- **cilia_io.py**: The top main file for the entire repository.
- **ml_helpers.py**: Helper functions used by CiliaIO for machine learning.
- **quantification_help.py**: Helper functions used by CiliaIO for quantification.
- **benchmarking_cilia_tools.py**: Benchmarking cilia_io versus binary thresholding, canny thresholding, and Thouvenin et al. pixel-based frequency segmentation
- **src/analysis/**: Contains all of the source files pertaining to downstream analysis with cilia_io

- **analyze_csv_data_violin.py**: Uses the cilia_io output CSVs to perform statistical tests, 3D graphics, Cohen’s power analysis, PCA, and violin super plots.
- **analyze_k_fold_cross_val_plots.py**: Contains utilities for coalescing k-fold cross validation plots together
- **benchmark_segment_anything.py**: Benchmarks zero-shot segment-anything on manual labeled data.
- **compare_csv_for_accuracy.py**: Script used for calculating the MAPE between manual / Thouvenin et al. and CiliaIO quantification outputs.
- **visualize_skeleton_over_time.py**: Script to take skeleton pickles out of cilia_io output and produce .PNG temporal waveform plots
- **src/dataset_creation**: Create datasets for YOLO fine-tuning and k-fold cross validation

- **create_yolo_training_dataset.py**: From raw label-studio projects, generates the combined train, validation, and test splits for the fine-tuning of the YOLO model
- **create_k_fold_cross_val_dataset.py**: Generate 3-fold stratified cross validation dataset for YOLO validation analysis
- **preprocess_nd2.py**: For dataset fine-tuning. Takes .ND2 files and converts them into individual frames for bounding-box labeling in label-studio
- **preprocess_tif.py**: For dataset fine-tuning. Takes .TIF files and converts them into individual frames for bounding-box labeling in label-studio
- **yolo/**: Contains the original Ultralytics Yolov11m model, the train and test scripts, as well as the final, fine-tuned model

- **configs.yaml**: YAML file for configuring the fine-tuning of the YOLO model.
- **save_training_curves.py**: Saves training curves from the YOLO training flow as .csv files
- **yolo_test.py**: Runs the test inference on the trained best YOLO model.
- **yolo_train.py**: Runs training and validation curves for 300 epochs on the dataset.
- **yolo11m.pt**: The actual pre-trained YOLOv11m model from Ultralytics.
- **k_fold_cross_val.py**: Runs k_fold cross validation using the dataset created by ’../src/dataset_creation/create_k_fold_cross_val_dataset.py’

### 0. System Requirements

Before you begin, ensure your workstation meets the following criteria:

#### Hardware

- **GPU (Required):** This software uses AI models that require high-performance graphics processing to function. Specifically, they call Torch libraries and use large ML models that typically require a GPU

- Windows/Linux: Requires a dedicated NVIDIA GPU.
- **Mac:** Supports **Apple Silicon (M1, M2, M3 chips)** using the integrated GPU via Apple’s Metal framework.
- **RAM:** At least 16GB is recommended for handling large .TIF stacks without system lag.

#### Software

- **Python 3.10 or higher:** The programming language used to run the scripts. (The code was specifically tested in Python 3.10.12)
- **Drivers:**

- **Windows/Linux:** CUDA Toolkit (compatible with your NVIDIA GPU).
- **Mac:** Metal Performance Shaders (MPS), which is natively supported on macOS for Apple Silicon.

#### Operating System & Environment

- **Windows:** The code has **not** been tested in the native Windows environment. You must use **WSL2 (Windows Subsystem for Linux)** to run the analysis.
- **Mac / Linux:** Standard Terminal is sufficient. The scripts are fully tested and supported on macOS and Linux distributions.

### 1. Acquiring the Software

You can obtain the CiliaIO files by downloading a stable release for publication citation (Zenodo) or by syncing the live development code (GitHub).

#### Option A: Stable Release (Zenodo)

Best for citing in papers or ensuring you are using the exact version used in a specific study.

1. Visit the CiliaIO Zenodo repository.
2. Download the `Source code (zip)` file from the latest version.
3. Extract the zip folder to your desired location on your computer. *Note: This version is a "snapshot" and will not receive automatic updates*.

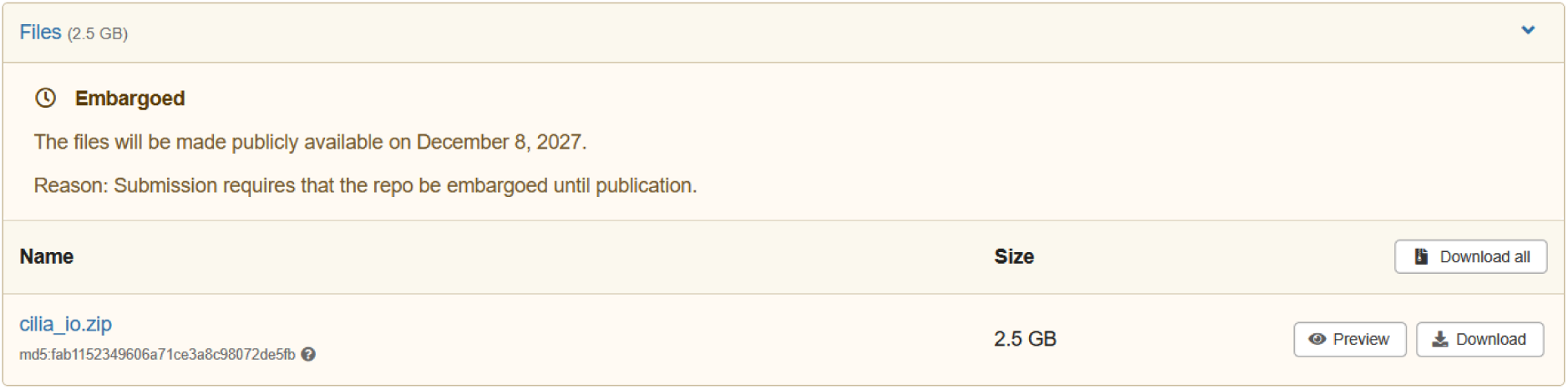

#### Option B: Graphical Method (GitHub Desktop)

Best for users who prefer a visual interface and want to keep the software updated with the latest bug fixes. This works on **Windows and Mac**.

1. **Download:** Go to desktop.github.com and install the app.
2. **Clone:** Click **File > Clone Repository**, select the **URL** tab, and paste the URL for cilia_io (https://github.com/chekfung/cilia_io.git).
3. **Update:** To get the latest code, open the app and click **Pull/Fetch Origin**. This "refreshes" your local folder with the newest lab updates.

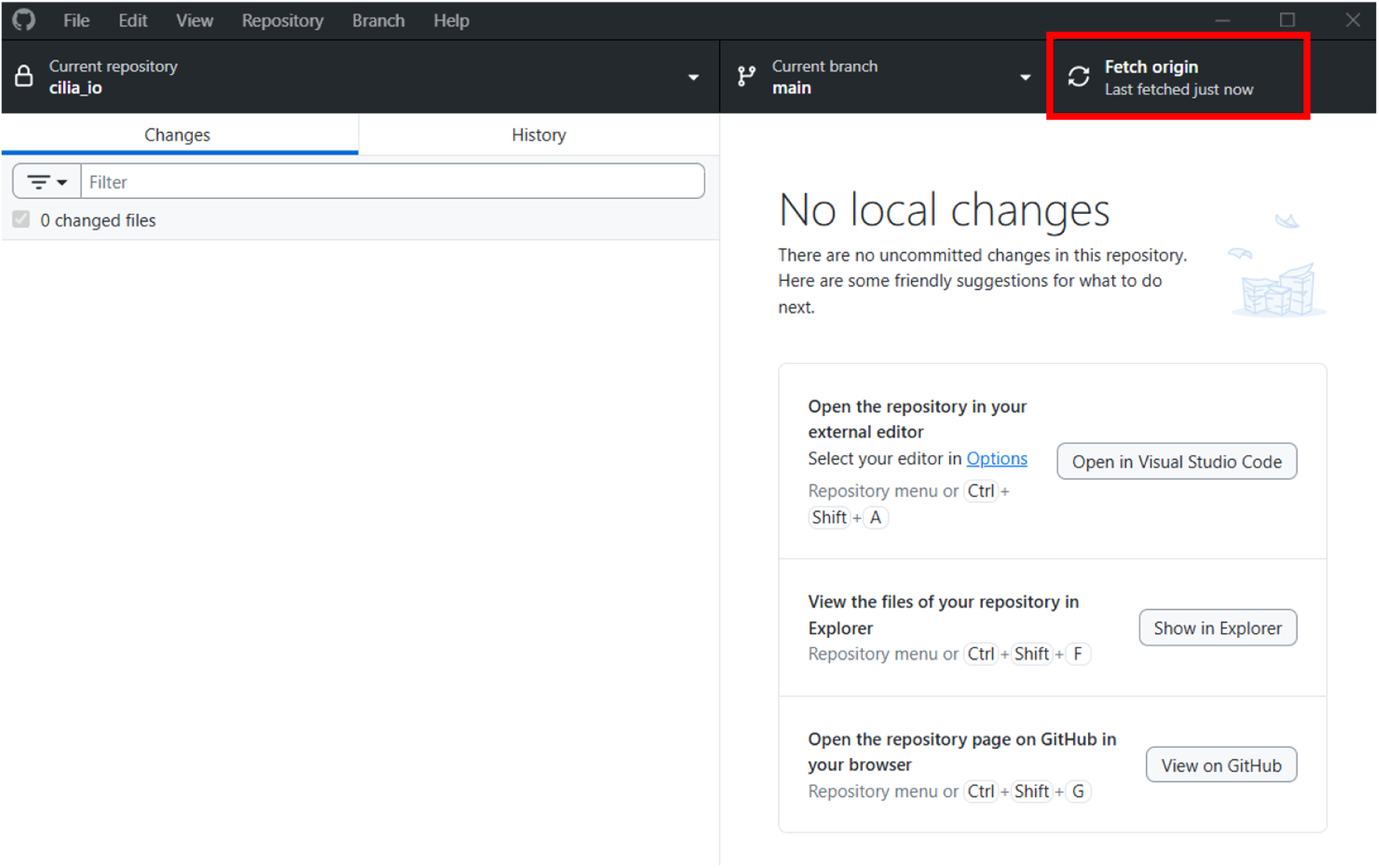

#### Option C: Command Line Method (Terminal)

Best for users comfortable with the command line (Windows/Mac/Linux) who already have Git installed.

1. Open your terminal (Ubuntu on Windows; Terminal on Mac/Linux).
2. Run the following command:

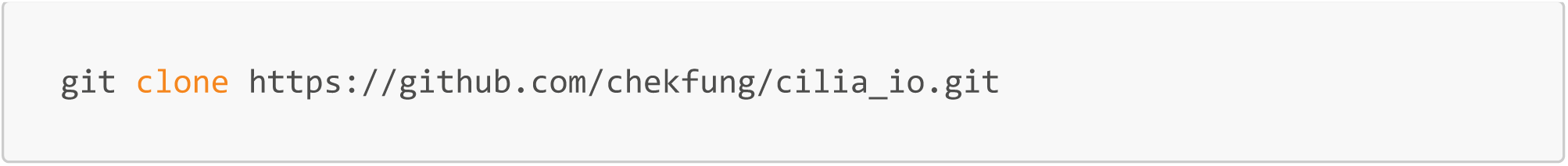

**NOTE: IF INSTALLING FROM ANYWHERE EXCEPT ZENODO, YOU NEED TO DOWNLOAD SAM AS WELL**

Downloading SAM for GitHub

The Segment-Anything Model is available at this link: https://dl.fbaipublicfiles.com/segment_anything/sam_vit_h_4b8939.pth

Download the model and then inside of the cilia_io repository, create a new folder titled sam in the root directory and move the model into that folder. The fine-tuned YOLO model is already installed in both Zenodo and GitHub, but the SAM model is too large. An example of the file structure is shown below.

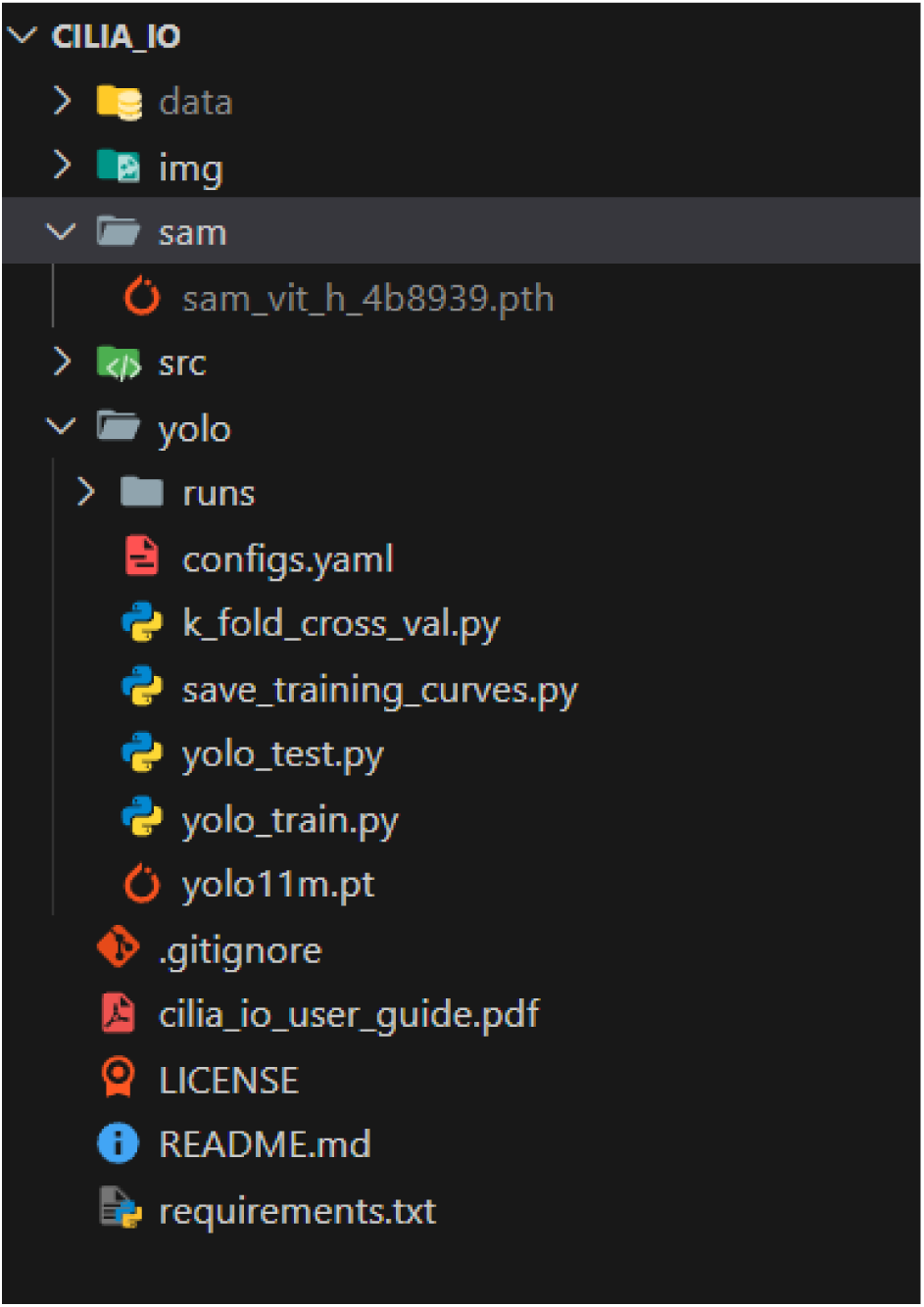

### 2. Installing Python and Instantiating the Virtual Environment

**NOTE:** This step only needs to be done once 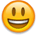

Note: While it is possible to already have Python completely installed and ready to go on your computer, to make sure that we do not run into dependency issues,werecommend following this section to install a virtual environment of Python with all the package dependencies needed for dataset creation.

#### Step 1: Open the Terminal (Command Prompt)

- On Windows:

1. Press the Windows key + R
2. Type "cmd" and press Enter
- On Mac:

1. Press Command + Space
2. Type "Terminal" and press Enter
- On Linux:

1. Press Ctrl + Alt + T

#### Step 2: Check if Python is Already Installed

1. In the terminal, type the following and press Enter:

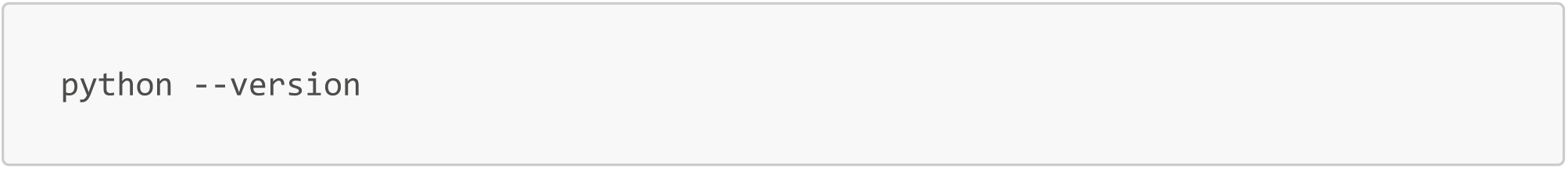 **Note that the keyword for python may not be python and could be python3, or py!**
2. If you see a version number (like "Python 3.10.0"), Python is already installed. Make sure that you have python3 (denoted by the first number before the decimal). You can skip to Step 4.
3. If you see an error or no version number, continue to Step 3.

#### Step 3: Install Python

- On Windows:

1. Go to https://www.python.org/downloads/ in your web browser
2. Click the "Download Python" button (it will download the latest version for Windows)
3. Run the downloaded installer
4. On the first screen, check the box that says "Add Python x.x to PATH" (very important!)
5. Click "Install Now"
6. When the installation is complete, click "Close"
- On Mac:

1. Go to https://www.python.org/downloads/ in your web browser
2. Click the "Download Python" button (it will download the latest version for Mac)
3. Run the downloaded installer package (.pkg file)
4. Follow the installation wizard, accepting the default options
5. When the installation is complete, click "Close"
- On Linux (Ubuntu/Debian):

1. In the terminal, type the following commands one by one, pressing Enter after each:

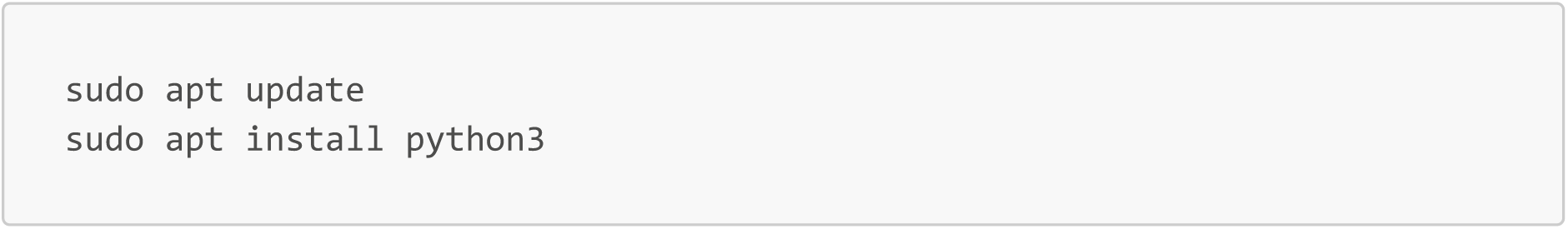
2. When prompted, enter your password and press Enter
3. When asked if you want to continue, type ’y’ and press Enter

In some cases, you won’t have sudo / superuser / admin access to a computing resource. In this case, please contact your respective IT department on installing python for additional help.

#### Step 4: Verify Python Installation

1. Close your current terminal window and open a new one
2. In the new terminal window, type the following and press Enter:

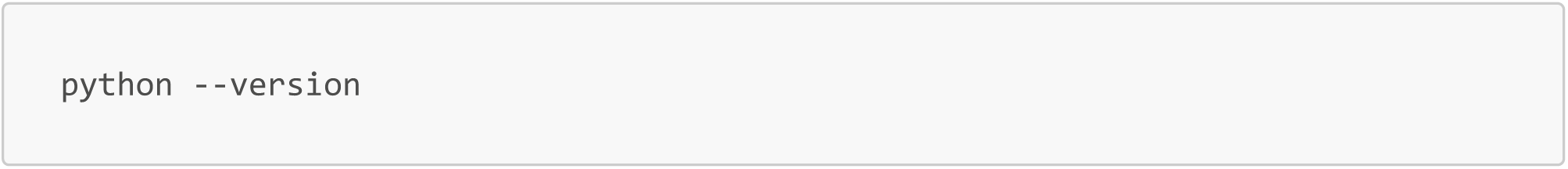 **Again, the keyword may be python, py, or python3**
3. You should now see a Python version number. If not, try restarting your computer and trying again.

#### Step 5: Install virtualenv

Virtualenv is a tool to create isolated Python environments. We’ll use it to create our virtual environment.

1. In the terminal, type the following and press Enter:

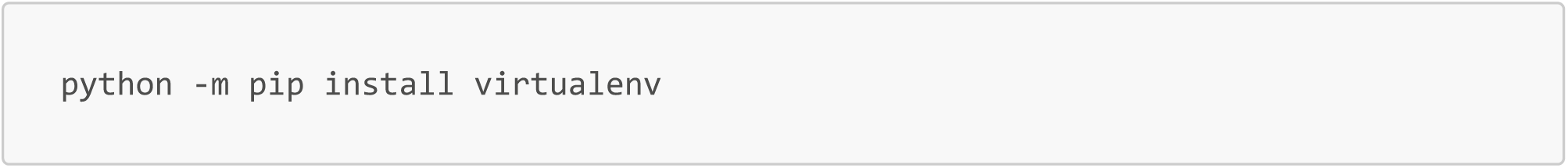
2. Wait for the installation to complete. You might see some progress information in the terminal.

#### Step 6: Navigate to an Area to Install Virtual Environment

1. Choose a location for your virtual enviornment. For this example, we’ll use the Desktop as the place in which we added the code for CiliaIO.
2. In the terminal, type the following commands one by one, pressing Enter after each:

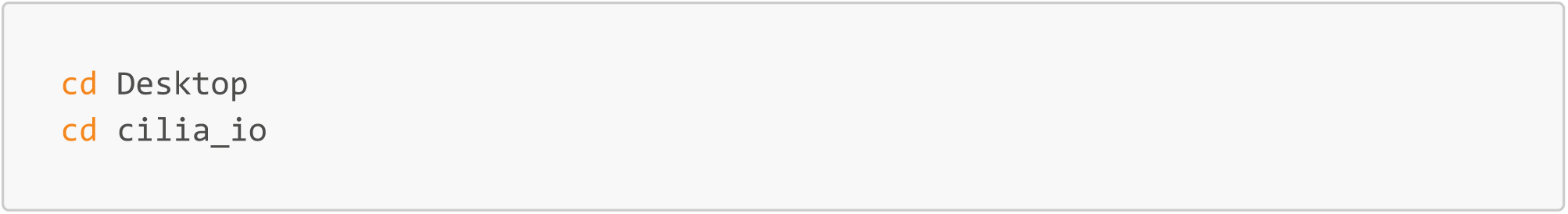 From the root directory that the terminal is opened in, this goes to the Desktop, and then to CiliaIO. If you are located anywhere other here, use the command cd ∼ which will reset your current directory to the root directory when the terminal begins.

#### Step 7: Create a Virtual Environment

1. Make sure you’re in your project directory (you should be if you followed Step 6)
2. In the terminal, type the following and press Enter:

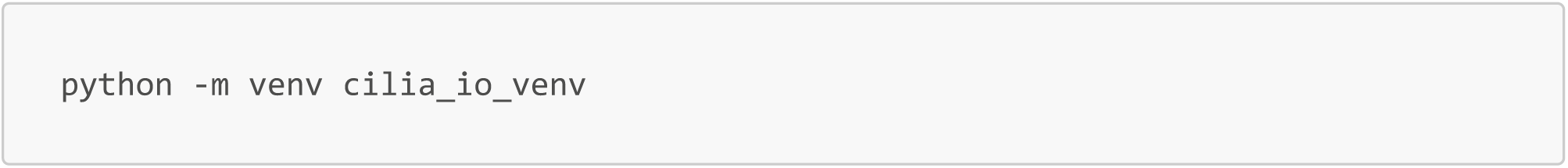 This creates a virtual environment named "cilia_io_venv". You can replace "cilia_io_venv" with any name you prefer, but just note that the keyword will be replaced in future commands with whatever you named it.

#### Step 8: Activate the Virtual Environment

1. 
  - On Windows:

1. In the terminal, type the following and press Enter:

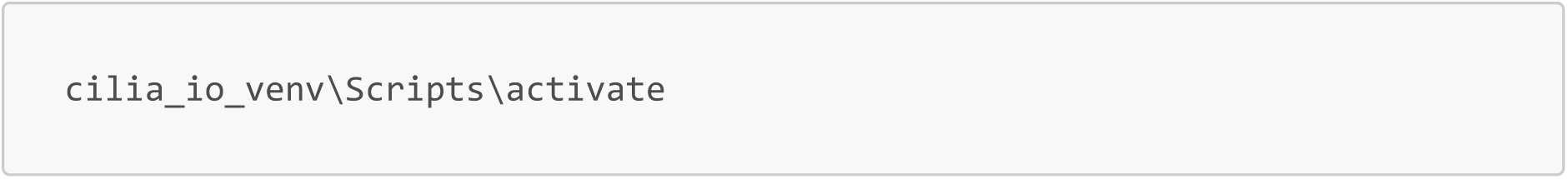
  - On Mac or Linux:

1. In the terminal, type the following and press Enter:

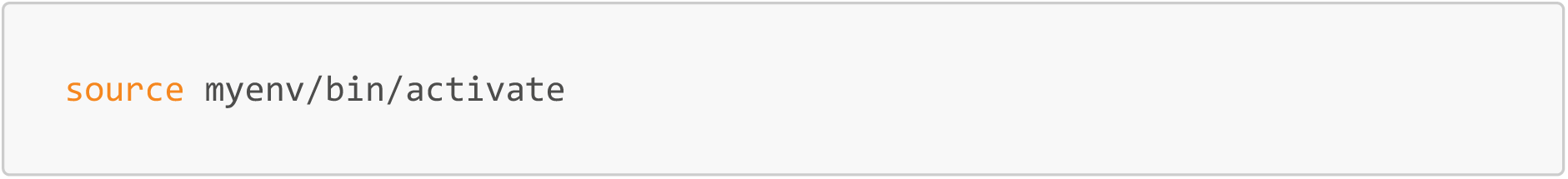
2. You should now see (cilia_io_venv) at the beginning of your terminal line. This indicates that the virtual environment is active.

#### Step 9: Using Your Virtual Environment

- To install packages in your virtual environment, use pip. For example:

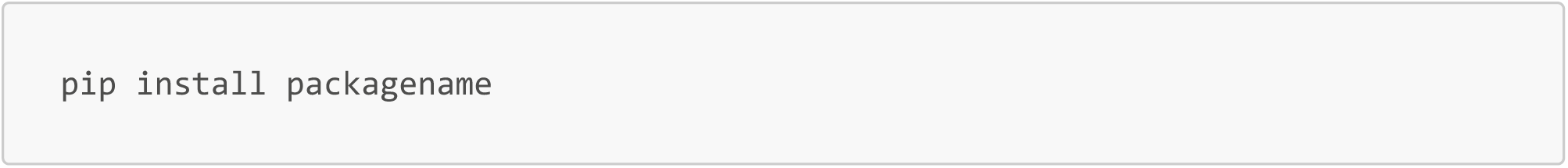 **Note: The next section will go over installation of everything, so no need to install any packages here.**
- To deactivate your virtual environment when you’re done working, simply type:

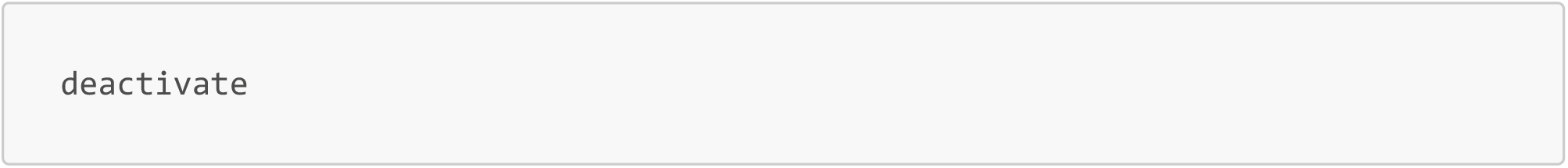
- To reactivate it later, navigate to your project directory and repeat Step 8.

**Remember to activate your virtual environment each time you work with CiliaIO!**

## 3. Installation of Requirements

**See Requirements.txt for the most up to date dependencies from Python. All results were run with Python 3.10**

To install the requirements, we again recommend a virtual environment to keep your Python package installation different than the ones required by CiliaIO. You can do this as follows.We would also recommend copy and pasting each line separately so that we can assess the errors independently.

**Also, if the terminal is unfamiliar,wehave provided a small cheatsheet at the end of the user guide on navigating the terminal and understanding some key commands.**

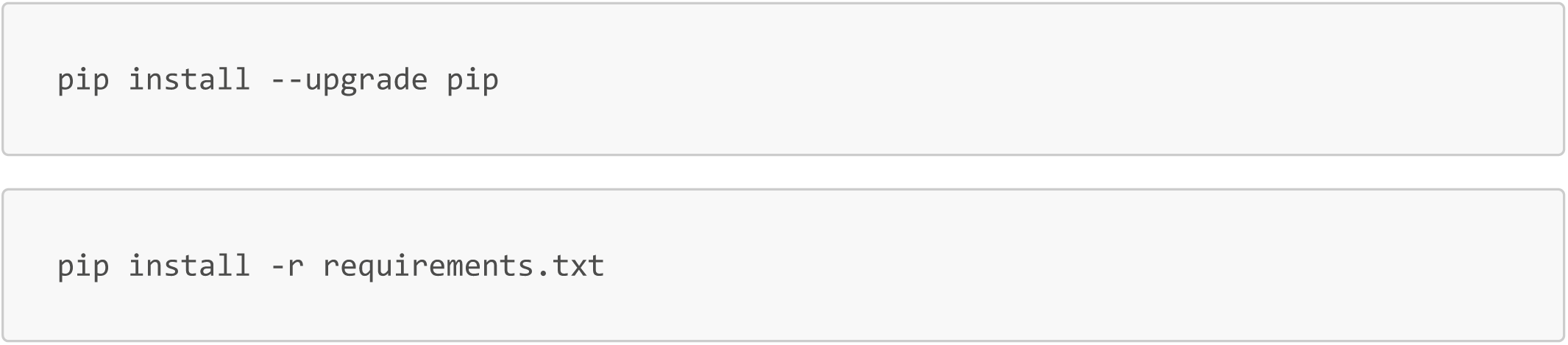

**Note:** This will take some time because it installs all the PyTorch and CUDA binaries. ∼15-20 minutes

## 4. Creating an Experiment Folder

While CiliaIO is flexible and can be pointed to any location on your computer, keeping your data organized within the project directory is recommended for simplicity.

1. **Open the `cilia_io` folder:** Navigate to where you downloaded or cloned the repository.
2. **Access the Data Directory:** Open the folder named `data/`. If it does not exist, create a new folder and name it `data`.
3. **Create your Experiment Folder:** Inside `data/`, create a new sub-folder specifically for your current batch of videos (.TIF)
4. **Move .TIF Files into Folder:** Inside `data/FOLDER_NAME`, insert your .TIF files.

Below is an example of the file structure of CiliaIO. The data folder is in the main directory of cilia_io.

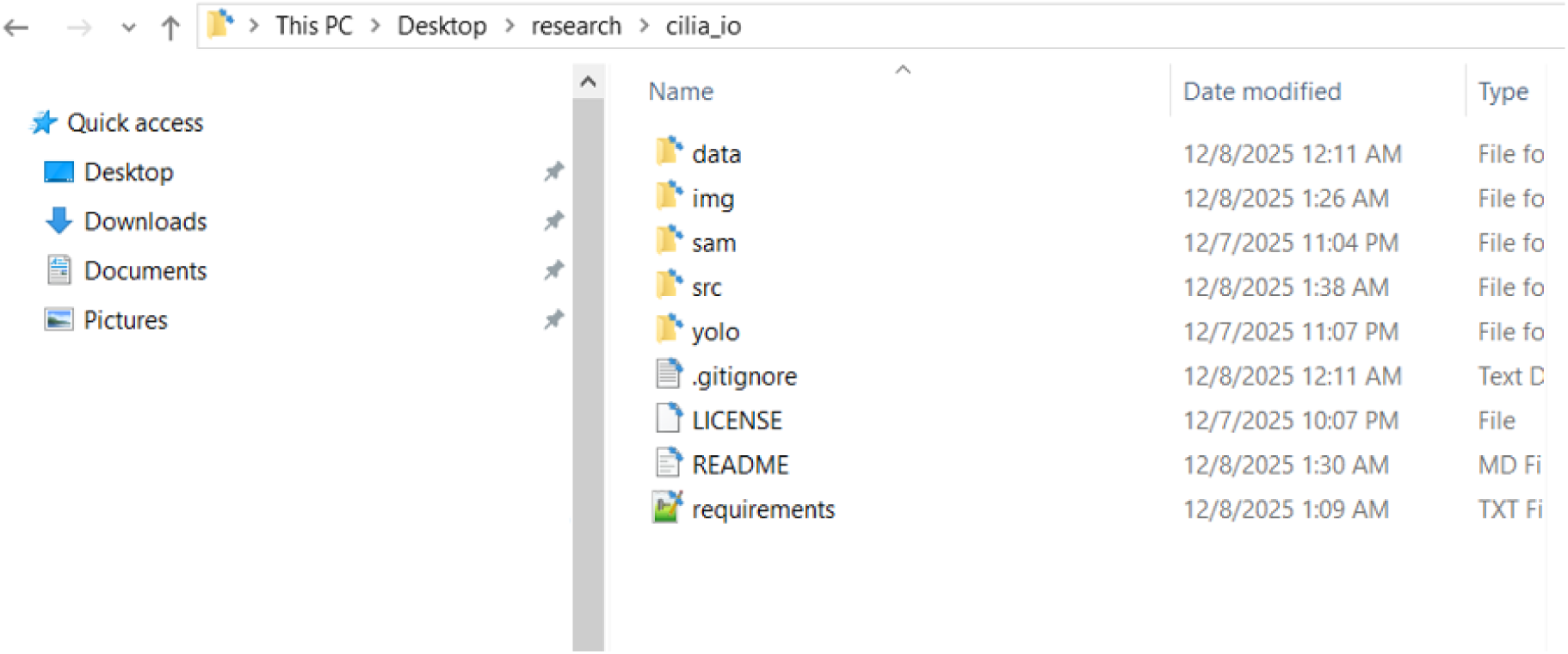

If we dive into the data folder, I created a subfolder, in this case, called `test_cilia_io`. Inside, there is a TIF file named `bbs2_21_f_vent.tif`. There is another folder, called `test_cilia_io_2` which can contain other .TIF files that I want to analyze in a separate call to CiliaIO.

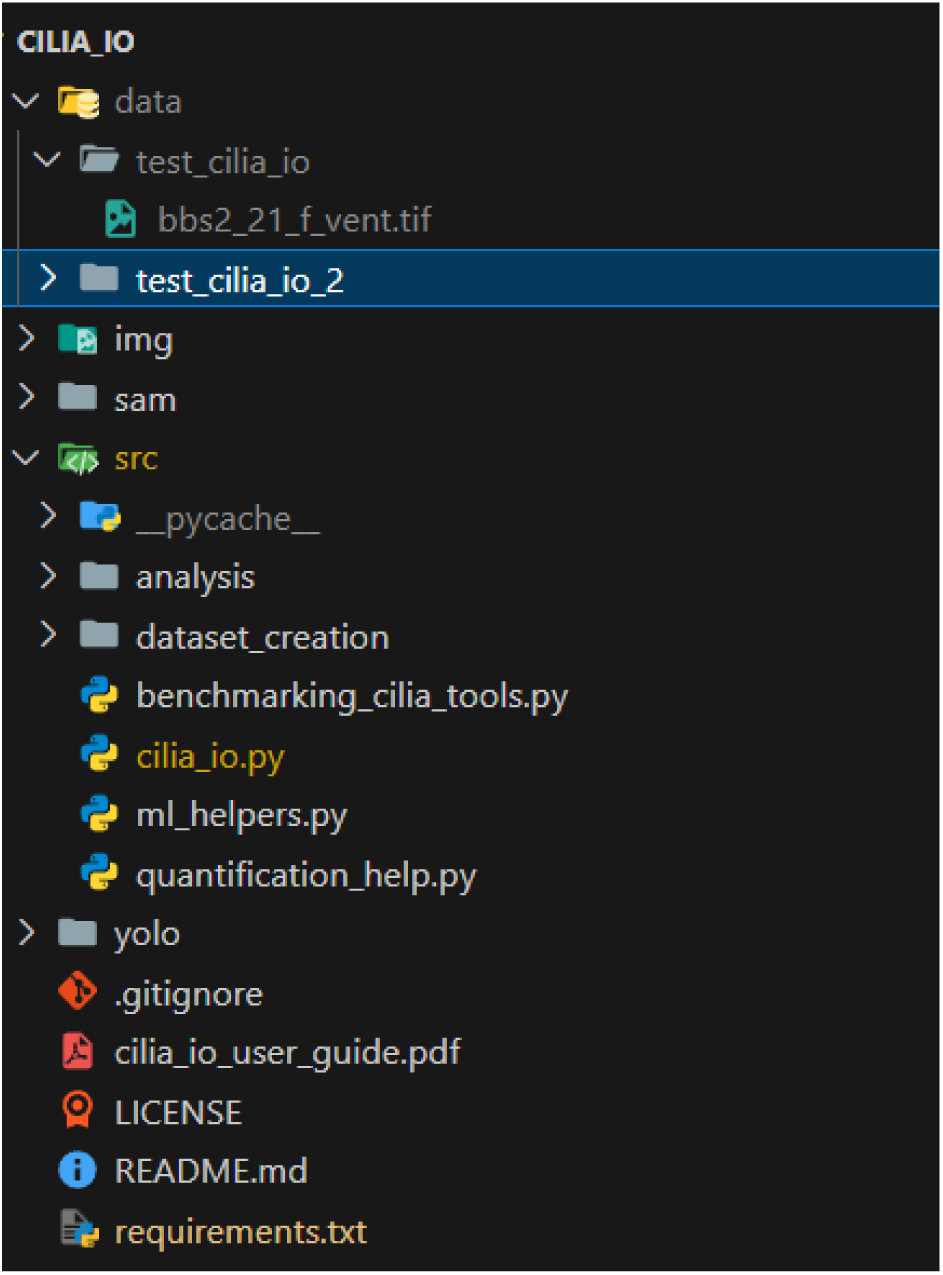

### Naming Rule (In general)

You must use underscores instead of spaces (e.g., `2026_04_30_cilia_test`). The terminal cannot easily read folder names that contain spaces.

### Note on Flexibility

If you prefer to keep your data on an external hard drive or a different local directory, you can do so. You will simply need to provide the **full file path** to that location when you run the script in the Terminal later. Using the internal `data/` folder is the easiest way to avoid path errors, but not required.

## 5. Preparing Data for CiliaIO

For this tutorial, we will put everything into the data folder. Inside of the data folder, create

1. **Copy Videos:** Place your raw `.TIF` files into the folder you just created.

## 6. Running CiliaIO

**Note: If you already installed everything, you do not need to repeat the steps above. START HERE!**

The pipeline processes all video files within a specified directory. Output results, including processed videos and data files, are saved directly into the source directory.

CiliaIO will go through every single .tif file inside of the folder in `input-dir`. This can then be used in the `analyze_csv_data_violin.py` scripts in order to join all the .csv files together and perform statistical analysis.

NOTE: To change CiliaIO to run file formats other than .TIF, convert to .TIF first.

1. Open your virtual environment from the previous steps by finding the folder where you installed it and using this command in a terminal. In our case, in this tutorial, we installed the virtual environment in the cilia_io directory:

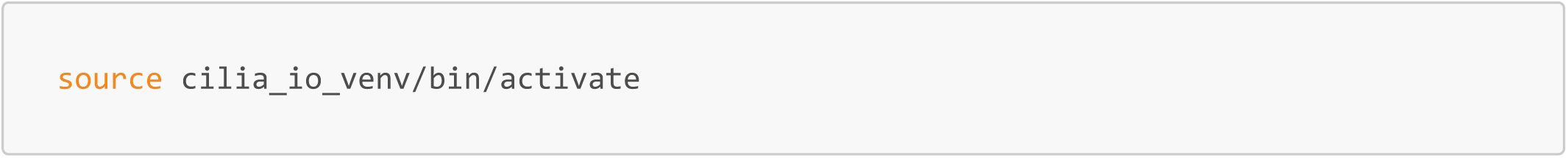

- Quick tip: The most immediate sign is the appearance of the environment name in parentheses at the start of your command line prompt.
  - Active: (cilia_io_venv) user@workstation:∼/cilia_io/src$
  - Inactive: user@workstation:∼/cilia_io/src$
- Navigate from there to the cilia_io/src directory.

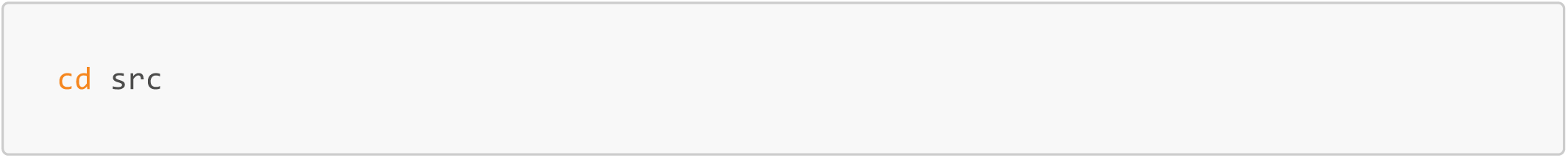

### Execution Commands

Below are some example commands to run cilia_io with different parameters. These are examples, and parameters can be combined into long command line arguments.

#### Dorsal/Ventral Static Y-Threshold Mode

This is the default mode. It applies a single horizontal Y-axis coordinate to every video in the folder to define the dorsal and ventral regions. Notably, the "/path/to/videos_directory" is a PLACEHOLDER where you will need to insert your directory that you want CiliaIO to process.

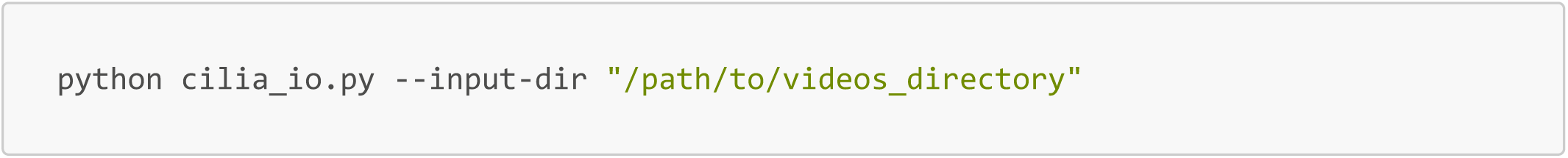

From our previous example using our folder created in data for cilia_io, we would perform the following command:

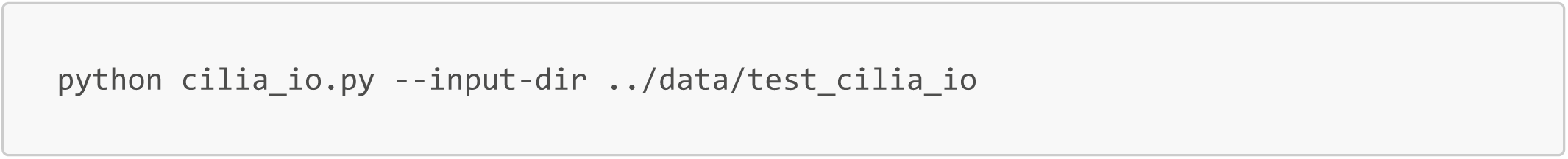

Quick explanation: In the terminal, .. is a shorthand command that means "go up one level" to the parent directory.

Because you are running the software from the src/ folder, but your videos are stored in the data/ folder, you must tell the computer how to navigate between them.

When you type ../data/test_cilia_io, you are giving the terminal a "relative path":

- ..: Exit the current src/ folder and move up to the main cilia_io/ project folder.
- /data: From that main folder, look down into the data/ directory.
- /test_cilia_io: Enter the specific folder containing your videos.

#### Batch List Mode

To apply specific, unique thresholds to videos within the same folder, disable the static toggle and provide a list. The thresholds are applied to files in alphabetical order. The command below applies a Y-axis threshold of 85, 72, and 90 to the first three videos, respectively. This is particularly useful because different videos may occur on different microscopes, have different scaling, etc.

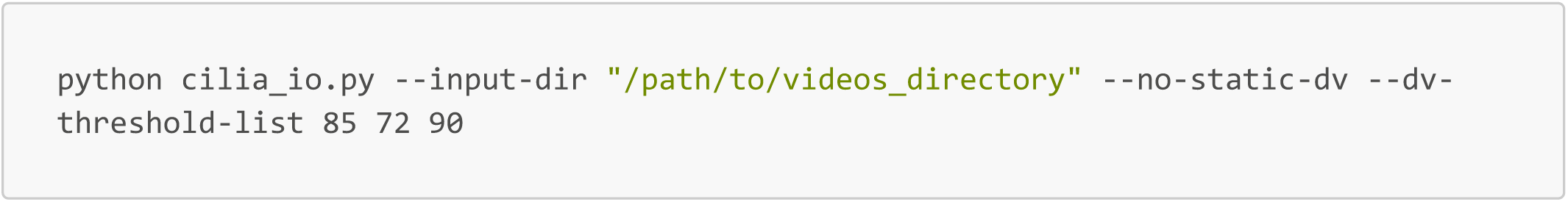

#### Manual Calibration for Metadata

If metadata extraction fails due to the specific microscope or corrupted metadata, you must manually provide the temporal and spatial scales (fps and number of microns in a pixel). The below shows an example where we disable the default metadata search, and instead provide the video-fps and pixel-to-micron conversions ourselves.

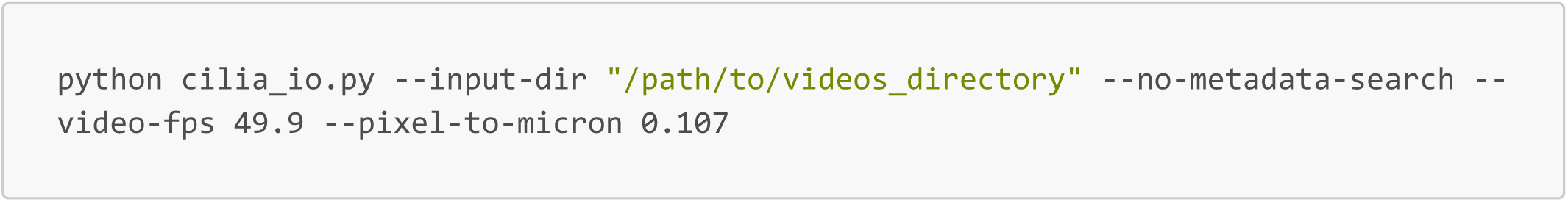

#### Another Real-World Example Usage

This is an example in which we disable the random colors for the cilia output video display, and instead make them based on the base dorsal ventral coloring scheme (can be adjusted in the source code under DORSAL_COLOR and VENTRAL_COLOR). Then, the output video fps is set to save at 1 FPS, with a minimum number of consecutively tracked frames of 1 to examine motility. The static dorsal ventral threshold has been turned off as well, and set to Y=80 for the first video, and 85 for the second. This is displayed by the blue line in the output videos. Early stop has also been turned on as a way to examine the first 5 frames to make sure that the dorsal ventral threshold is set correctly.

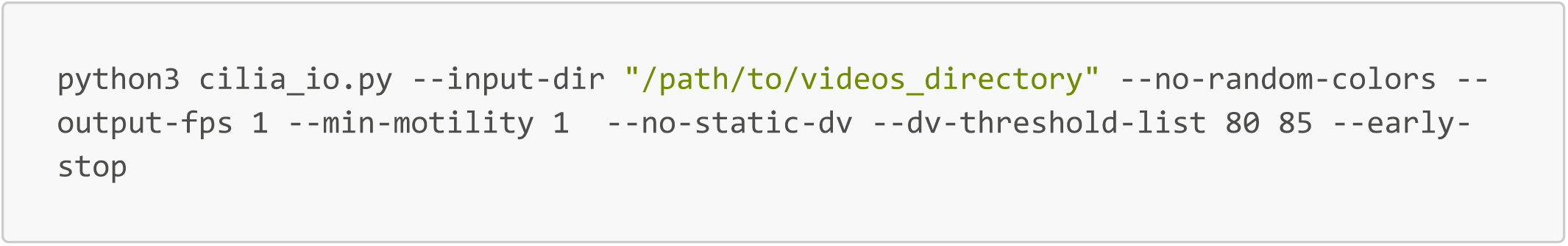

### All Parameter Definitions

These parameter definitions are also available using this command line function for reference:

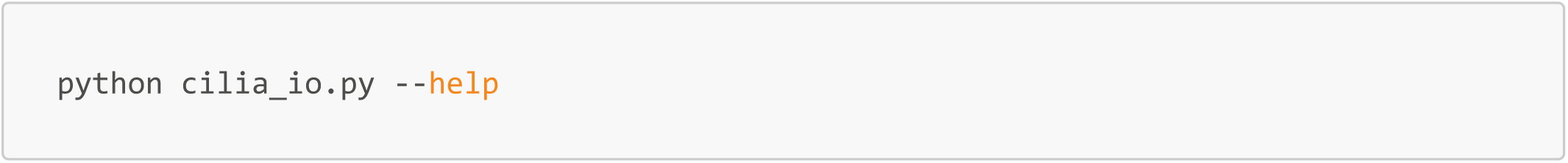

#### Required Parameters

- `--input-dir:` Path to the directory containing source videos.

##### Dorsal Ventral Y-Axis Thresholding Parameters

- `--static-dv | --no-static-dv:` Toggles between using a single threshold for the entire batch or a sequence of thresholds from the list. Default is to use one static-dv for all video files
- `--dv-threshold:` Integer value representing the Y-pixel coordinate for the horizontal cut. Default is 80.
- `--dv-threshold-list:` A space-separated list of integers used when static-dv is disabled.

##### Calibration and Metadata

- `--metadata-search | --no-metadata-search:` Toggles automated extraction of FPS and scale from file headers. Default is to search for metadata if not explicitly specified.
- `--video-fps:` Value for frames per second. Required if metadata-search is disabled.
- `--pixel-to-micron:` Value for spatial conversion (AKA. How big is a pixel in microns). Required if metadata-search is disabled.

##### Detection and Tracking

- `--matching-conf:` YOLO confidence threshold for point identification. Default is 0.4.
- `--max-dist:` Maximum pixel distance allowed for matching IDs between frames. Default is 30.
- `--max-age:` Number of frames to retain an unmatched ID before dropping it. Default is 5.
- `--post-iou-thresh:` Intersection over Union (IOU) threshold used for bounding box matching. Default is 0.5.

##### Skeletonization and Motility

- `--interpolation-pts:` Number of nodes used for spline resolution during skeletonization. Default is 100.
- `--min-motility:` Minimum consecutive frames required to calculate motility metrics. Default is 42.

##### Visualization and Debugging

- `--output-fps:` Playback speed for the generated result videos. Default is 5.
- `--random-colors | --no-random-colors:` Toggles between unique track colors or dorsal/ventral specific coloring. Default is random colors if not specified.
- `--early-stop:` If enabled, the script processes only the initial frames for debugging purposes. Default is False.

#### CiliaIO Outputs (per video file)

1. Converted mp4 file: *{file_name}.mp4*
2. Converted ciliaIO annotated output: *{file_name}_5fps_ciliaio_output.mp4*
3. Converted ciliaIO annotated output with original fps: *{file_name}_original_fps_output.mp4*
4. Center of Box .PKL: *{file_name}_box_center_points.pkl*

- This is maintained as a dictionary with key (cilia_id) and is a list of tuples consisting of (frame_id, (x_i,y_i)) for the box centroid
5. Center of Box Tracking Cilia Quantification .CSV: *{file_name}_quant_box_center.csv*
6. Center of Skeleton .PKL: *{file_name}_skel_center_points.pkl*

- This is maintained as a dictionary with key (cilia_id) and is a list of tuples consisting of (frame_id, (x_i,y_i)) for the skeleton midline
7. Center of Skeleton Tracking Cilia Quantification .CSV: *{file_name}_quant_skel_center.csv*
8. Full cilia skeleton .PKL file: *{filename}_full_skeletons.pkl*
  - This is maintained as a dictionary of dictionaries. First indexed by cilia_id, then by frame_id, and then the 2D matrix where each row represents the an (x,y) coordinate of one point on the skeleton. The skeleton is ordered, meaning that when incrementing the row index, it represents the next point on the skeleton.

## 7. Troubleshooting and Common Problems

Even with a perfect setup, you may encounter errors in the terminal. Most issues in CiliaIO are related to file naming, folder paths, or your computer’s hardware.

### Hardware & Environment Errors

| Error Message | What it Means | The Fix |
| --- | --- | --- |
| **python3: command not found** | Python is either not installed or not "mapped" in your terminal. | **Windows/WSL:** Type `sudo apt update && sudo apt install python3`.<br>**Mac:** Install Python via python.org. |
| **CUDA Out of Memory** | Your NVIDIA GPU does not have enough memory to process the current video. | 1. Close Chrome, Zoom, or other GPU-heavy apps.<br>2. Try processing a smaller video clip. |
| **ModuleNotFoundError** | A required software library is missing. | Run `pip install -r requirements.txt` again from the main project folder. |
| **NVIDIA-SMI has failed** | The terminal cannot talk to your GPU. | **Windows:** Update your NVIDIA drivers on the Windows side. **Mac:** Ensure you are on a Mac with an M1/M2/M3 chip. |

### Data & Folder Errors

| Error Message | What it Means | The Fix |
| --- | --- | --- |
| **FileNotFoundError** | The computer cannot find your data folder or metadata file. | 1. Check for typos (Linux is case-sensitive).<br>2. Ensure you are in the `src` folder.<br>3. Ensure your folder name has **no spaces**. |
| **IndexError** or **KeyError** | There is a mismatch or typo inside your `metadata.csv`. | 1. Check that the "filename" column matches your `.TIF` file names exactly.<br>2. Ensure there are no empty rows at the bottom of the spreadsheet. |
| **Permission Denied** | The computer is blocked from accessing a file. | Make sure the video file isn't currently open in Excel, ImageJ, or a video player. |

### Terminal Navigation Cheatsheet

| Command | Meaning | Description |
| --- | --- | --- |
| `pwd` | **P**rint **W**orking **D**irectory | Displays the exact path of the folder you are currently in. |
| `ls` | **L**ist | Shows all files and folders inside your current directory. |
| `cd <folder>` | **C**hange **D**irectory | Moves you into a specific folder (e.g., `cd src`). |
| `cd ..` | **C**hange **D**irectory **..** | Moves you **up** one level to the "Parent" folder. |
| `cd ~` | **C**hange to **H**ome | Returns you immediately to your primary user home directory. |

## Supplementary Materials and Methods - CiliaIO Fine Tuning Guide

For any questions, feel free to email me at to debug.

### Setting Up Your Environment

Make sure to have followed the `cilia_io_user_guide.pdf step 2` and `step 3` which creates a virtual environment for CiliaIO in Python and installs all the required libraries necessary for CiliaIO to function.

To get CiliaIO ready for fine-tuning, we need to add one more package, `label-studio`, that will do a lot of the heavy lifting for us for YOLO fine-tuning. The following section details how to install this package.

#### Installing Label Studio

**Note:** This only needs to be completed once.

1. Activate your virtual environment. See the cilia_io_user_guide.pdf for more guidance on loading into your virtual environment.
2. Run the following command:

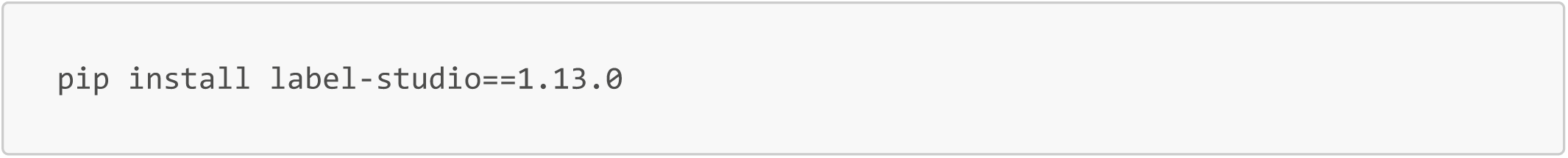

After it runs, there should be no errors in the terminal.

**Note:** We ran this with Python 3.10 with label-studio 1.13.0. There have been updates since then, but for exact replication of our flow, please use 1.13.0

### Converting Raw Videos to Individual Frames for Label Studio

CiliaIO contains two helpful scripts for converting raw .ND2 files and .TIF files into individual .PNG frames for individual annotation of each of the separate frames necessary for Label Studio.

Such files are located at `/src/dataset_creation/preprocess_nd2.py` and `/src/dataset_creation/preprocess_tif.py`, respectively.

The usage of these files is detailed below. We recommend creating a separate output directory for each video file such that the video frames are contained together and not mixed with other frames, which can be confusing in downstream analysis or future analysis.

#### preprocess_nd2.py

This file is the preprocess script used to preprocess .nd2 zstack files and produce black and white pictures of each frame of the video and save them into a directory. The usage is as follows:

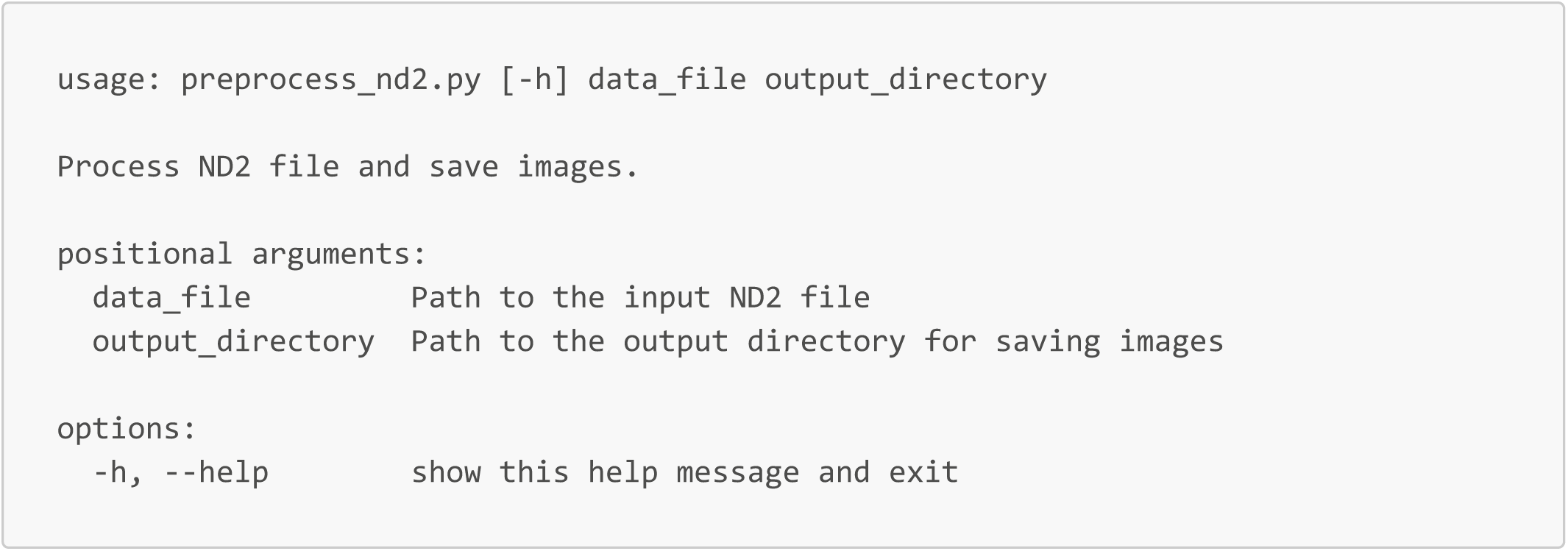

This is a command line python file that takes in two things:

1. **First argument:** The relative location of the .nd2 file
2. **Second argument:** The folder to save the frames of the specific folder to

#### preprocess_tif.py

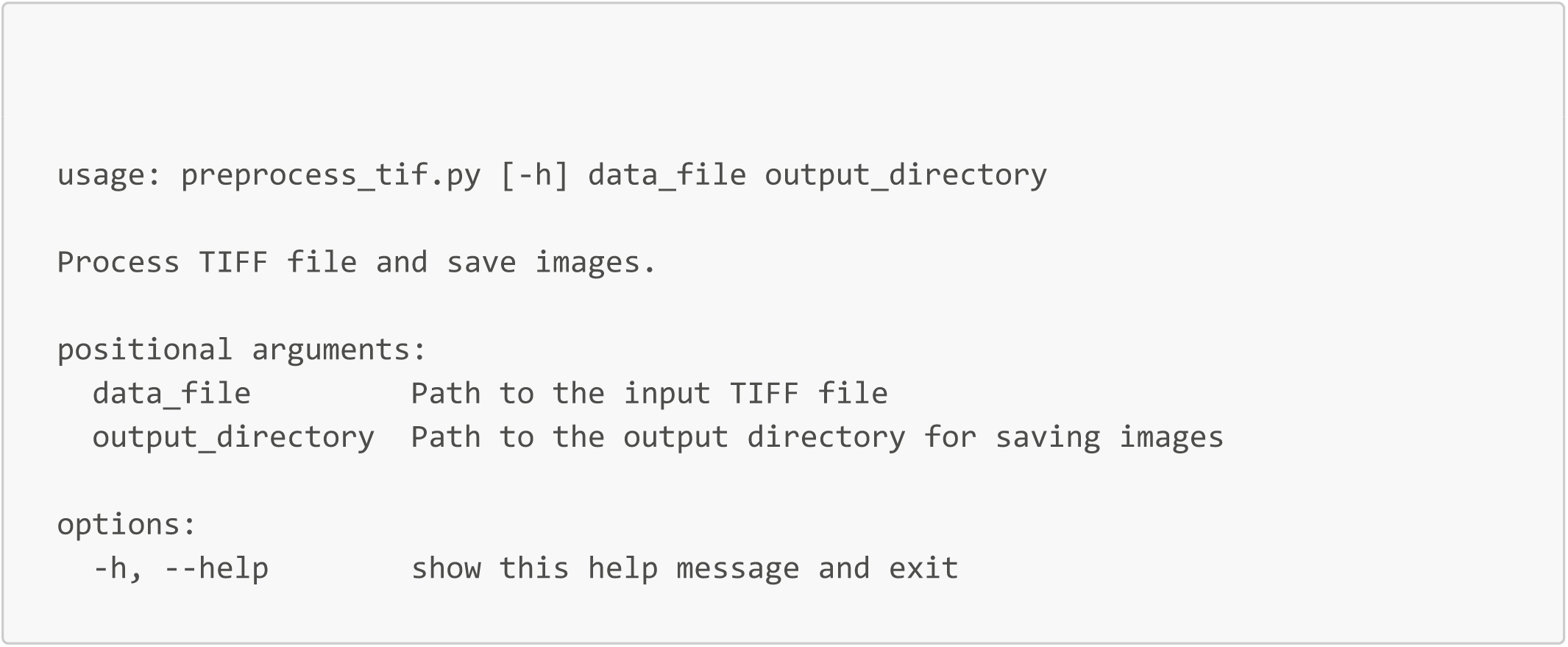

This is a command line python file that takes in two things:

1. **First argument:** The relative location of the .tif file
2. **Second argument:** The folder to save the frames of the specific folder to

##### Example Usage

I put a file named: *example_cilia_video.nd2* into my cilia project directory. This file, *preprocess_nd2.py* is in the same folder. I want to create a new folder labeled to house all of this information called: **example_cilia_frames** with the preprocessing done. I would run the following command:

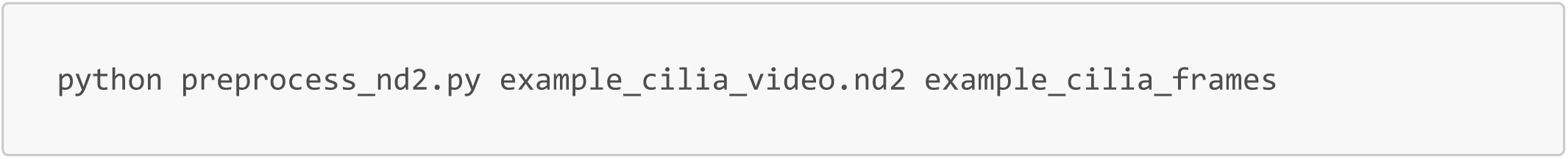

### PLEASE DO NOT MIX DIFFERENT VIDEOS OR RENAME THE FRAMES AS THIS METADATA IS USEFUL FOR FUTURE TRAINING

#### Launching Label Studio and Using Label Studio

Label-studio is the primary tool that used for the labeling of cilia. Here, there is a nice interface which should be friendly to non-code users that effectively acts as a website that is hosted locally. Here, you will create a project for each video, create the labels, and then label each of the frames in the video by hand. Additionally, you will export all of the data from each labeled video out of this program.

##### Step 1: Launching label-studio and creating an account

1. After generating a directory from the video file, launch label-studio using the following command in the terminal **while you are still in your virtual environment**:

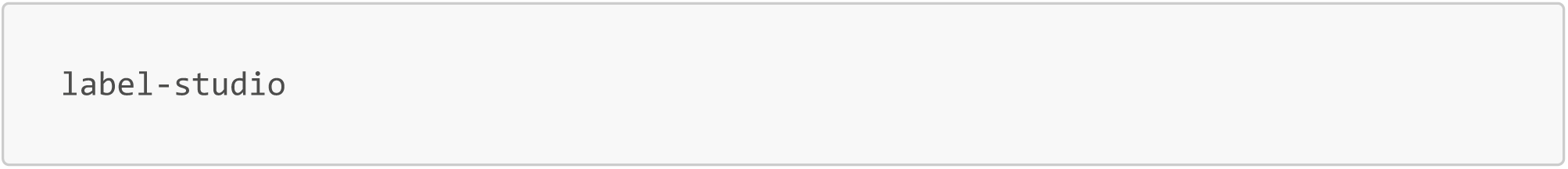 **Make sure not to close the terminal window!** The terminal should hang and state that it is starting a development local server for label studio hosted at 0.0.0.0:8080 as shown below: **Note: If this does not work, MAKE SURE YOU ARE IN YOUR VIRTUAL ENVIRONMENT.** You will see (virtual_environment_name) in the front of your terminal line. In our case, (cilia_io_venv)

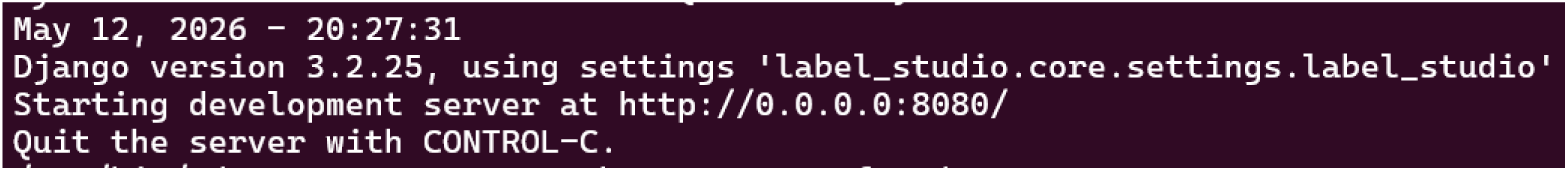
2. This call to label studio will create a locally hosted website on your computer. Navigate to your favorite web browser and in the search bar the following hyperlink to navigate to the open label-studio.

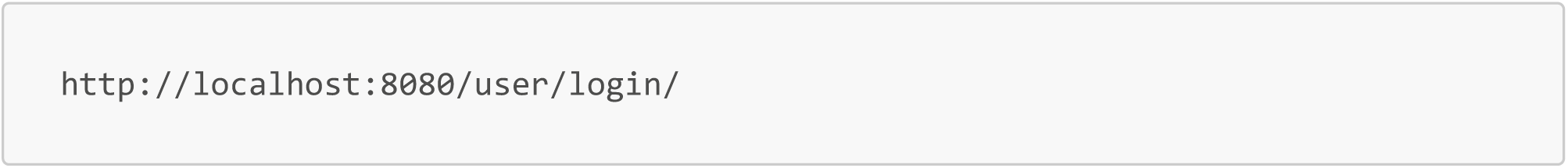
3. If you are a new user, create an account through label studio. Note that this is a local account. The application is hosted entirely on your machine. This means that no confidential data is put on the internet that should not be available.

##### Step 2: Create a New Project

1. Click on the "Create" button (usually found on the top right hand corner of the dashboard).
2. Give your project a name (e.g., "cilia_io_test_video").

##### Step 3: Import Your Data

1. In your new project, navigate to the Data Import Tab.
2. Click Upload Files
3. Select the frames that you want to upload. Note that the project only really supports loading 80 images at a time, so if there are more than 80 frames, you will have to do multiple uploads.
  - For multiple uploads, just do 80 frames at a time (0-79, 80-159, etc.) where after you upload the first 80, just click Upload More Files where “Upload Files”’ button used to be
4. Wait for the upload to complete.

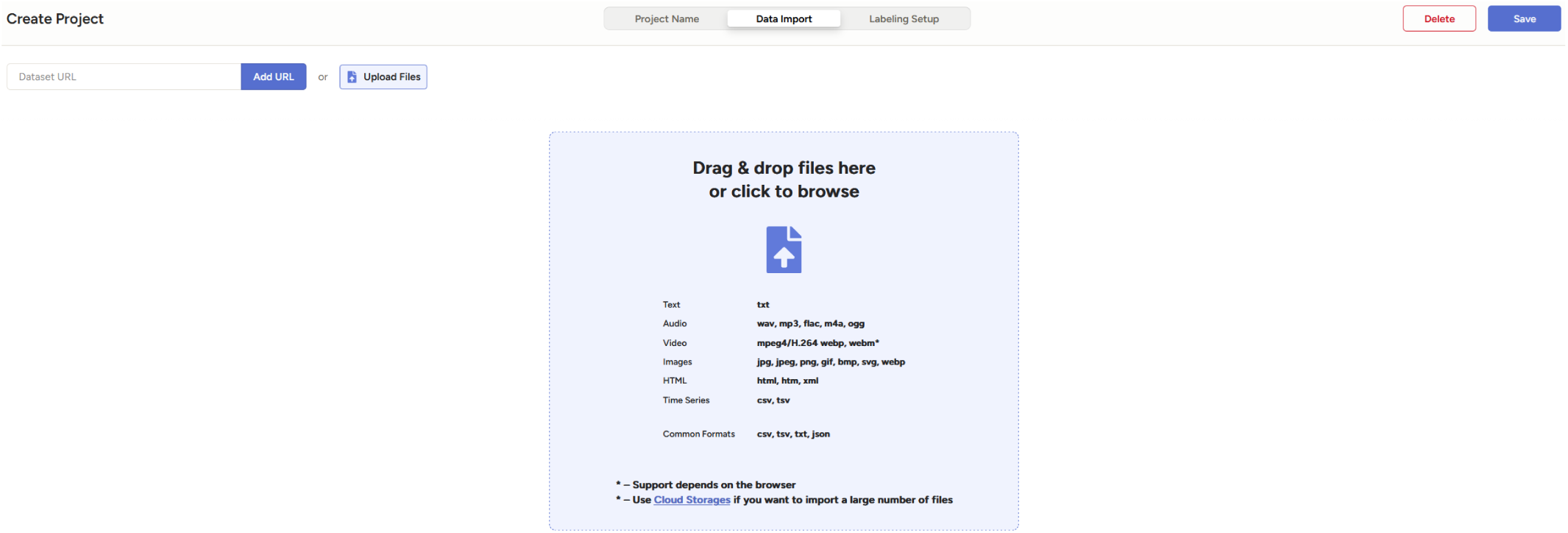

##### Step 4: Set Up Your Labeling Interface

1. Go to the `Labeling Setup` tab in your new project.
2. Navigate to the `Computer Vision` tab and click `Object Detection with Bounding Boxes`
3. This will bring up a picture of airlines with two labels (airplane and car).
4. Delete the two labels by clicking on the red x next to their names.
5. Under `Add label names` add a label named `cilia` and click add.
6. Check the box that says: `Allow image zoom (ctrl+wheel)` (not necessary, but useful to zoom in)
7. Save your changes.

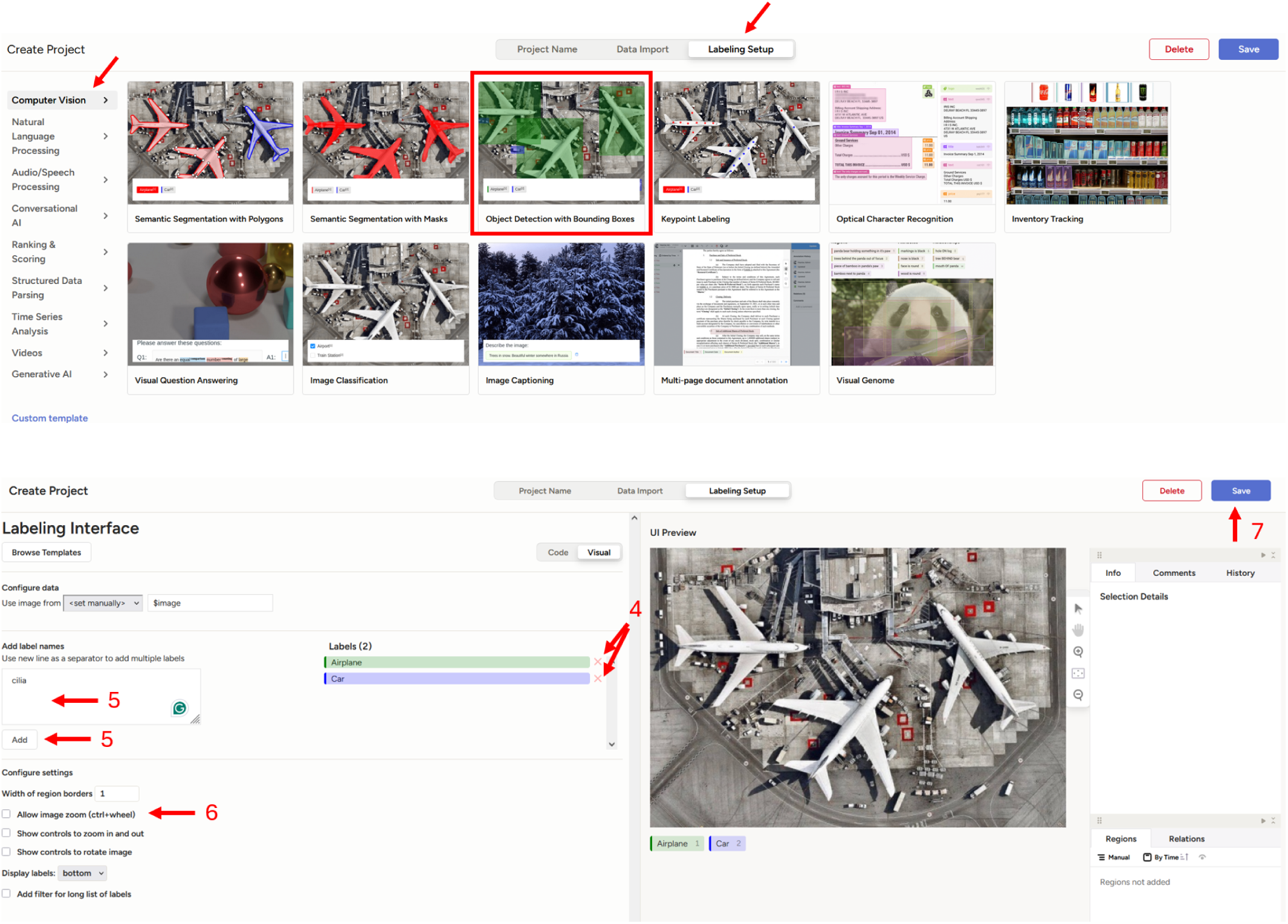

##### Step 5: Start Labeling

1. Click on `Label All Tasks` in blue
2. You’ll see your data items one by one.
3. For each item:
  - Look at the data carefully.
  - On the bottom, click the cilia button label which will highlight the cilia button.
  - Left click and drag to make a bounding box for a single cilia
4. Click "Submit" or "Next" to move to the next item.

**Note: The data is saved internally after submitting a labeled image. There is no manual saving needed.**

Below, we show what one cilia bounding box label looks like in a frame in Label Studio.

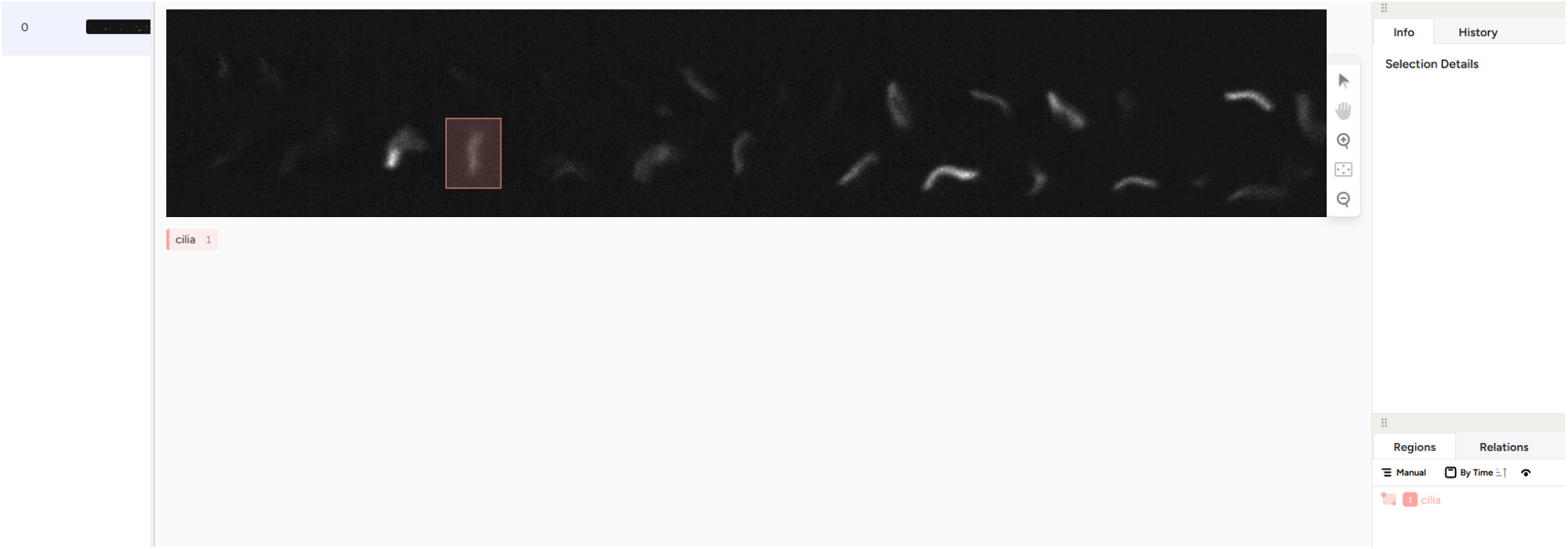

##### Step 6: Review and Adjust

1. Periodically check your work in the "Results" or "Review" section.
2. Look for any mistakes or inconsistencies in your labeling.
3. You should edit or delete incorrect labels here.

##### Step 7: Export Your Labeled Data

1. Once you’ve labeled all the data in one videos, go to the "Export" section.
2. Select `YOLO` as the export configuration
3. Click "Export" and save the file to your computer.
4. This will produce a zip file containing the images, the bounding box labels, and the number of labels in this dataset.

Here is what that screen looks like when exporting. The red arrows indicate the initial export button in the top right-hand corner, the YOLO export configuration, and the Export button.

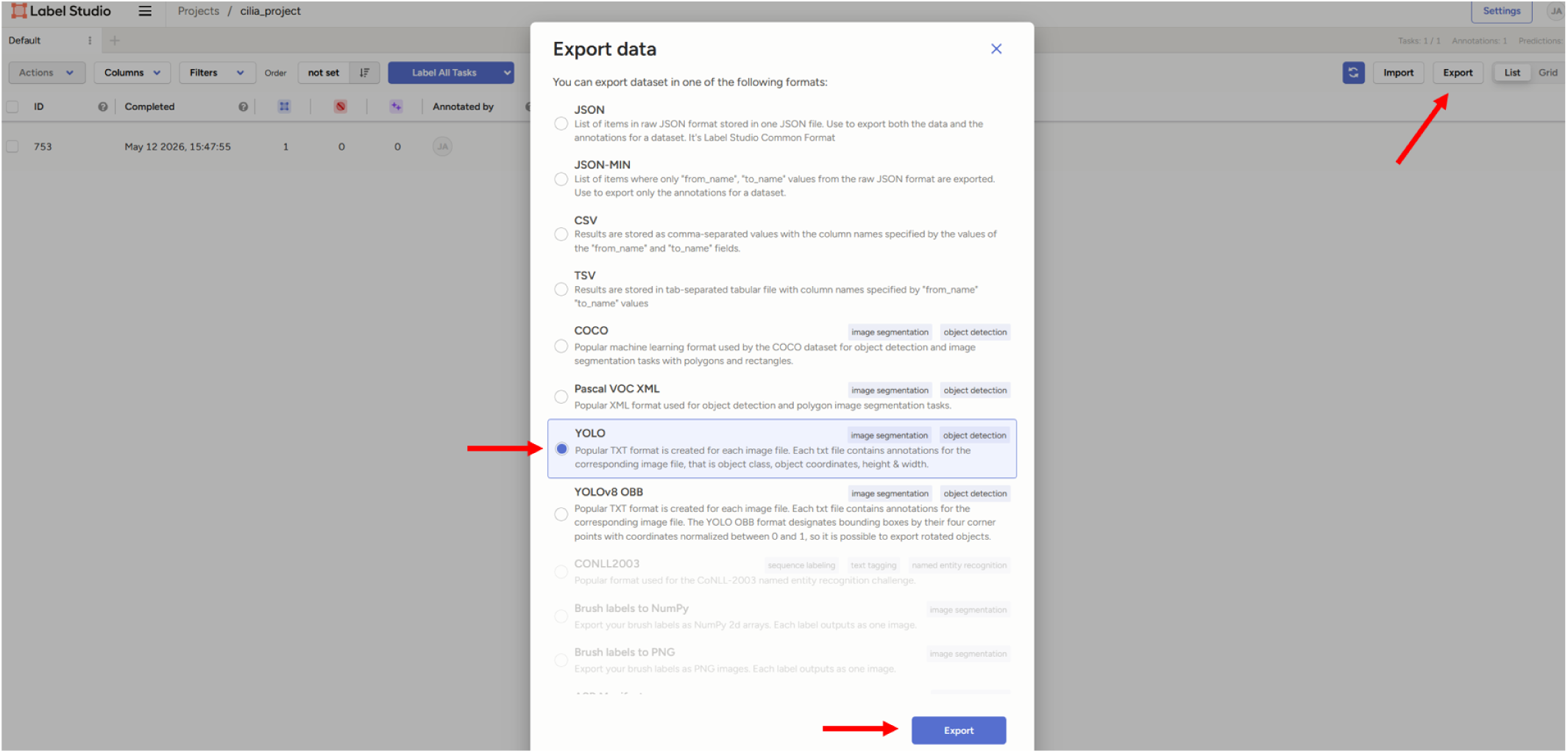

### Stopping a Label Studio Session

- After finishing labeling, you can close out the tab as everything is automatically saved.
- Go to your terminal and close the locally hosted label-studio server with CTRL + C on Windows or Linux, or CMD + C on Mac
- To open label-studio again, follow the previous instructions to open the virtual environment in the directory where you first installed your virtual environment, and then use the label-studio command to startup the server and login with the credentials that you first signed up with!

## Fine Tuning Ultralytics YOLO

### Setup Fine-Tuning Dataset from Label Studio Outputs

The following section will explain the format of each of the .zip files and how to incorporate these to create a dataset that YOLO can train, test and validate on.

#### Understanding Each Exported .ZIP from Label Studio

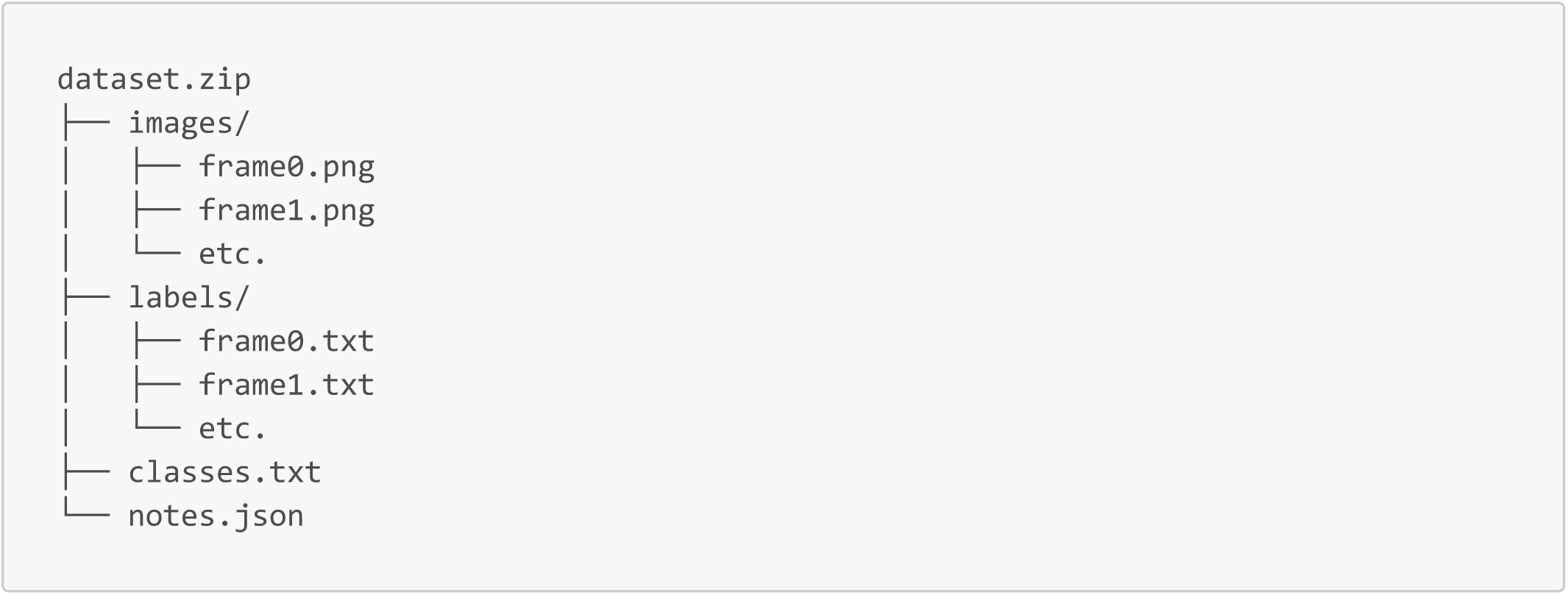

- `images/` — image files
- `labels/` — YOLO annotation .txt files
- `classes.txt` — class names
- `notes.json` — dataset metadata

Each label file corresponds to an image with the same filename.

YOLO annotation format:

~~~
<CLASS_ID> <X_CENTER> <Y_CENTER> <WIDTH> <HEIGHT>
~~~

#### Unzipping and Preparing a Dataset of Videos

If you have multiple labeled videos, unzip all exported folders and place them inside the same parent directory.

Example:

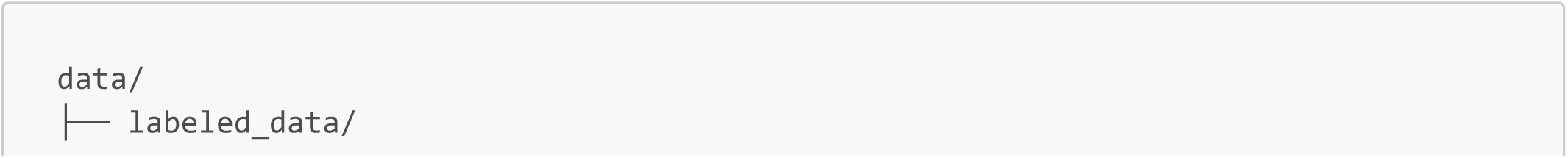
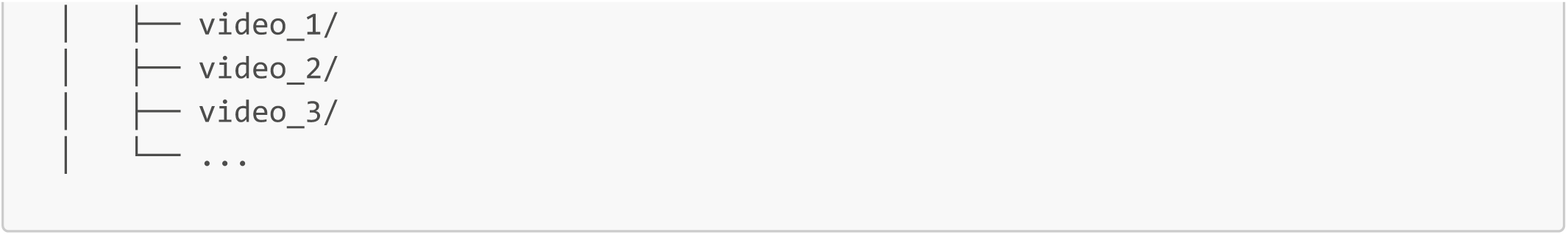

The script, `src/create_yolo_training_dataset.py` will automatically process each video folder and create the YOLO dataset.

In src/create_yolo_training_dataset.py, update lines 51–54:

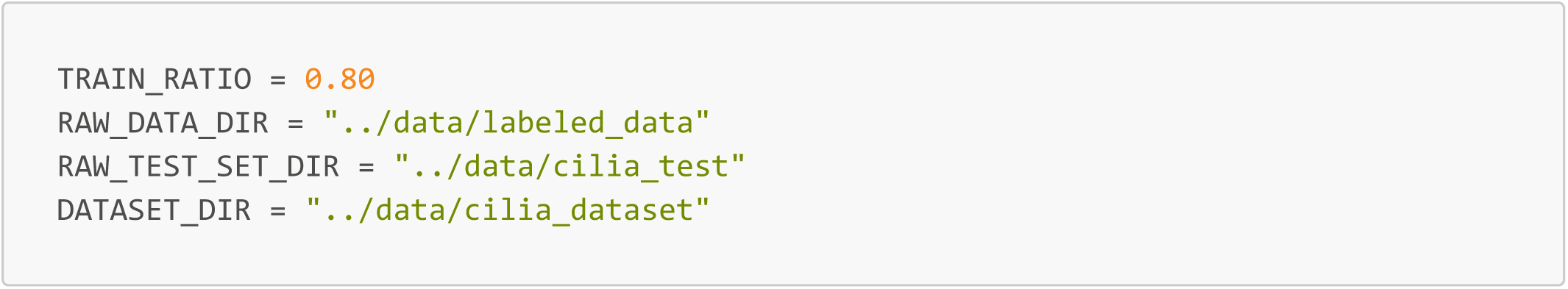

Variable Descriptions

- `TRAIN_RATIO:` Percentage of videos used for training. Here, 0.80 means 80% training and 20% validation.
- `RAW_DATA_DIR:` Folder containing all labeled video folders for training and validation.
- RAW_TEST_SET_DIR: Folder containing the separate test videos. These videos are not used during training.
- `DATASET_DIR:` Output folder where the final YOLO dataset will be created.

#### Running the YOLO Dataset Generation Script

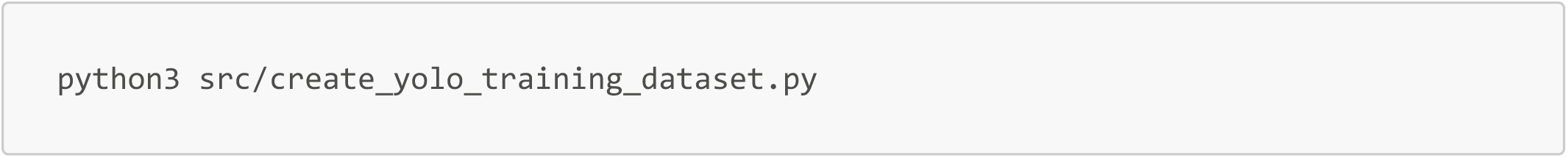

Create a YOLO-formatted dataset inside DATASET_DIR

he final dataset structure will look like:

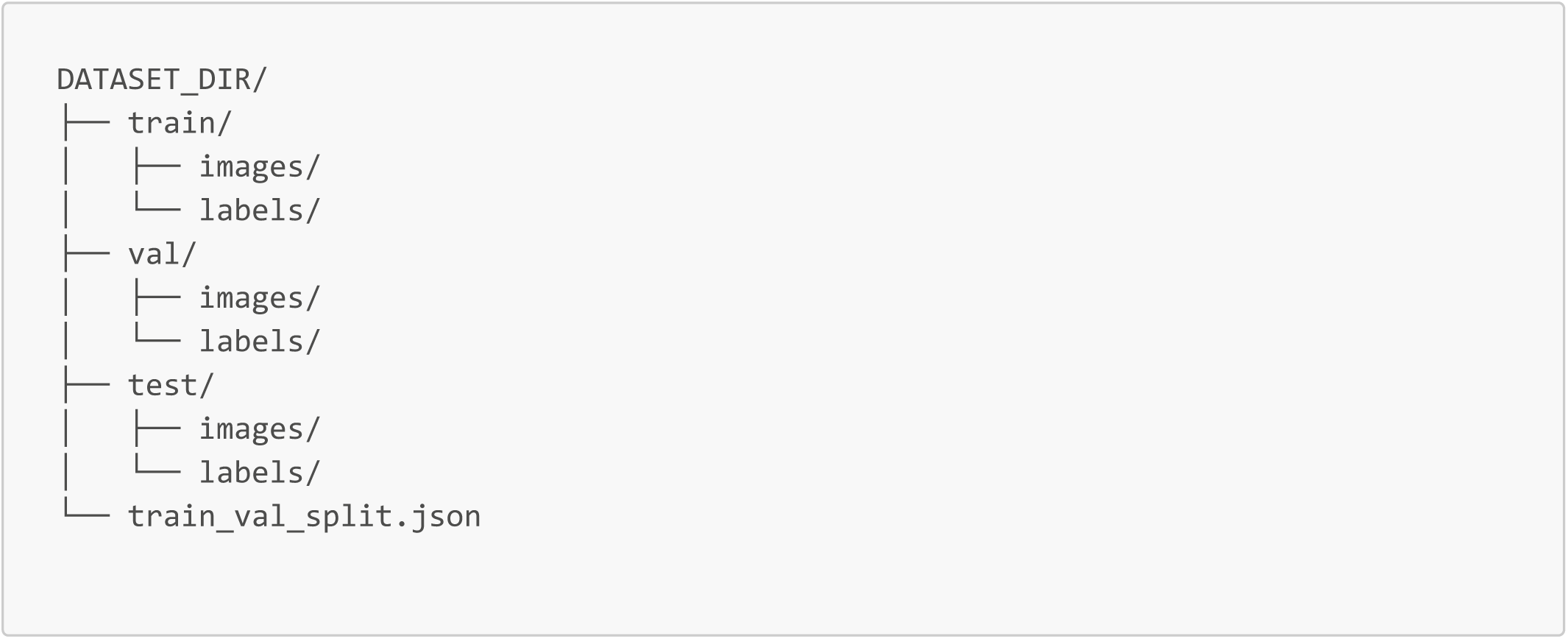

The script combines all of the training set videos together, all the validation set videos together, and test set videos together and provides a .json file explaining the exact split of everything.

### Running YOLO Fine Tuning!

The training script is located at /yolo/yolo_train.py

Before running training, update the dataset path inside /yolo/configs.yaml. This file looks like this.

Example:

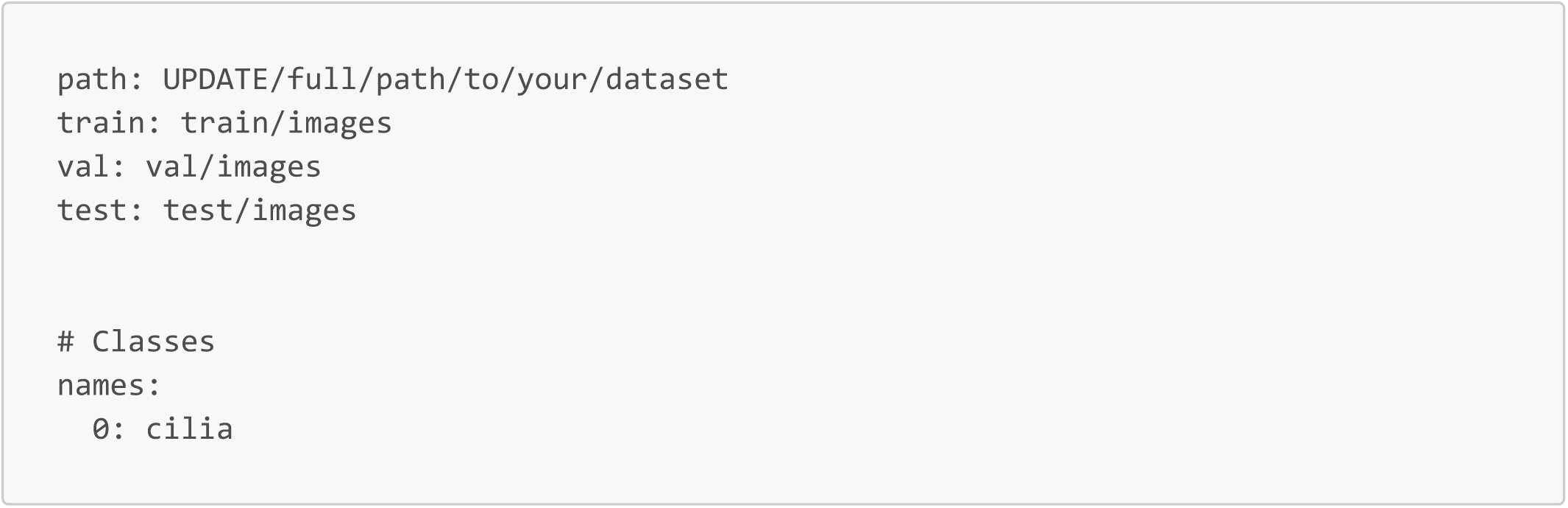

**Only update the path field so that it points to your YOLO dataset directory! This is also where we can add updated parameters to the YOLO Ultralytics model, but in order to get the most up to date documentation, check out the YOLO documentation on the Ultralytics website.**

#### Running Training

From the main project directory, navigate to the YOLO directory and run the yolo_train.py.

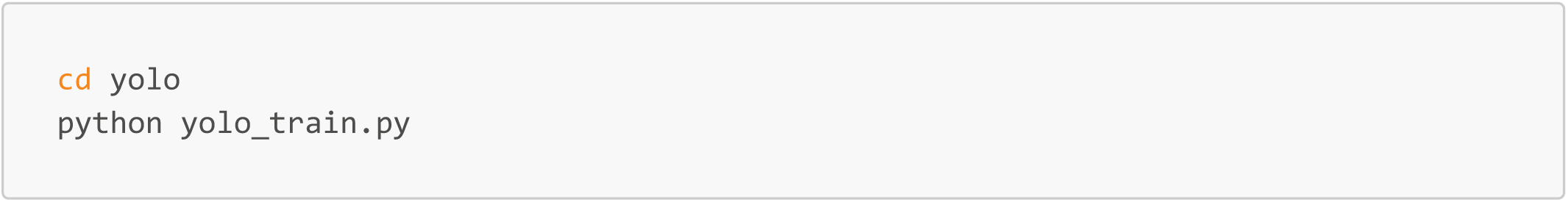

### Outputs of Ultralytics YOLO Fine-Tuning

After training finishes, Ultralytics will create an output folder containing the trained model, plots, and evaluation metrics.

The outputs are usually stored in:

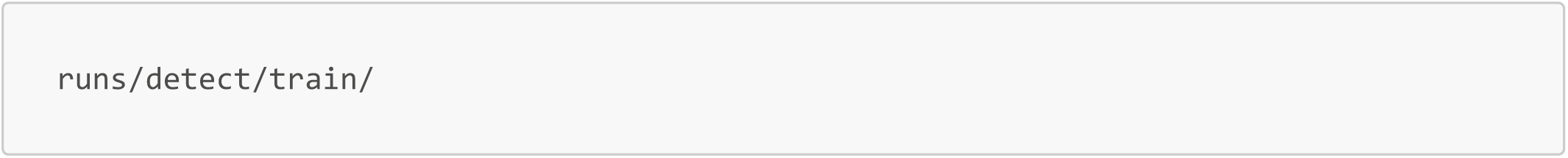

Example structure:

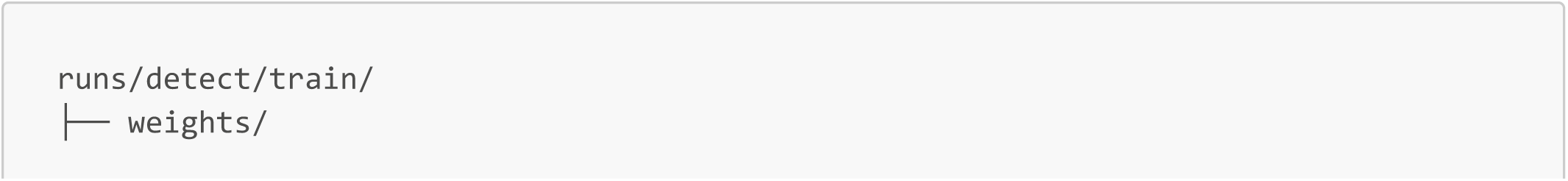
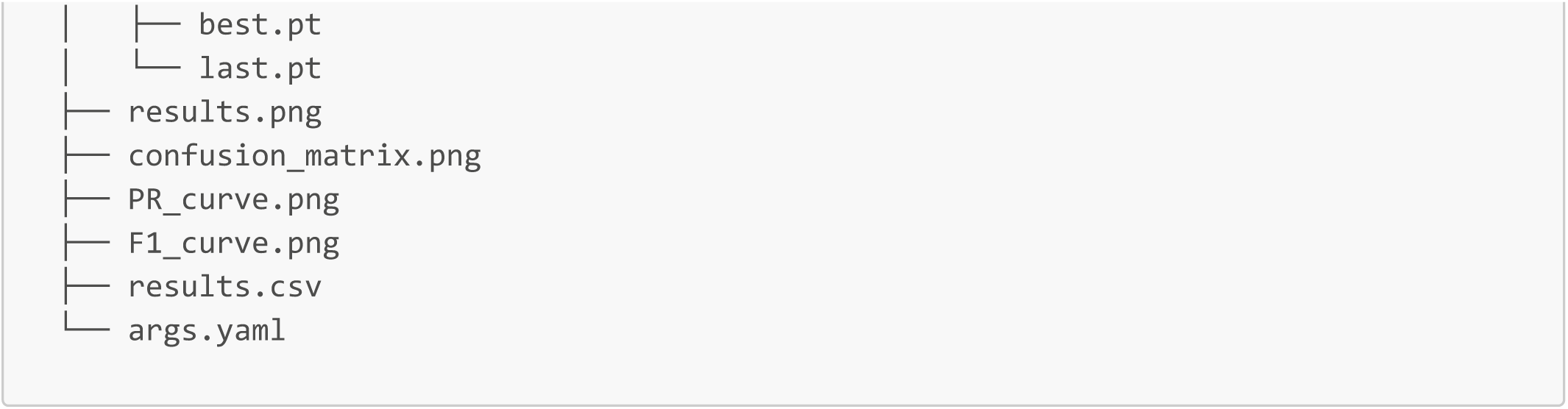

#### Important Files

- weights/best.pt: Best-performing trained model based on validation performance. This is usually the model used for inference/testing.
- weights/last.pt: Model weights from the final training epoch.
- results.png: Summary plot showing training and validation metrics over time.
- confusion_matrix.png: Shows prediction accuracy and common classification errors.
- PR_curve.png: Precision–Recall curve for model performance evaluation.
- F1_curve.png: F1 score across confidence thresholds.
- results.csv: Numerical training and validation metrics for each epoch.
- args.yaml: Copy of the training configuration and parameters used for the run.

## Using the New Fine-Tuned Models in CiliaIO

1. To incorporate the newly fine-tuned YOLO models, navigate to line 202 in cilia_io.py and update the filepath to point to the new checkpointed fine-tuned YOLO model.
2. Run CiliaIO with no other changes!

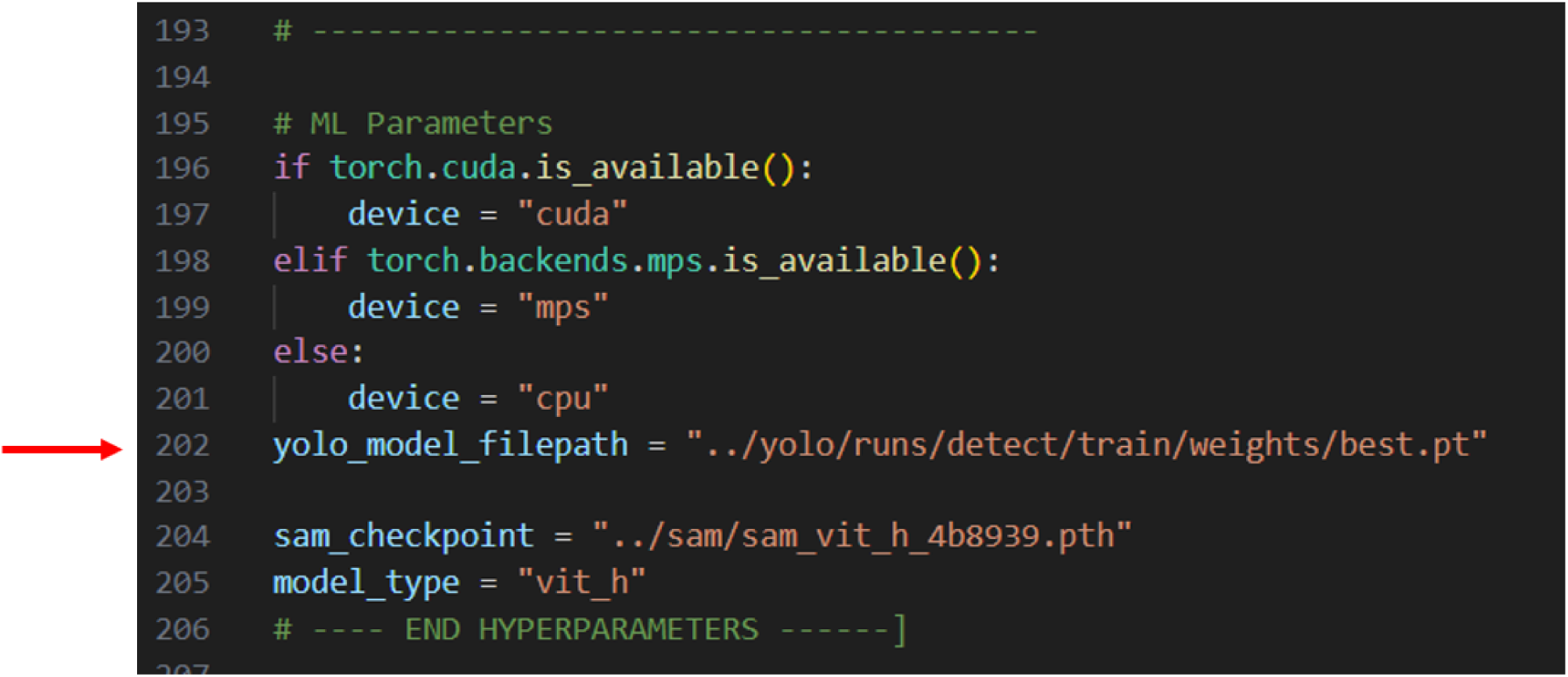

## REFERENCES

Afzelius, B.A., 1976. A Human Syndrome Caused by Immotile Cilia. Science 193, 317–319. 10.1126/science.1084576

Aton, M., McDonald, D., Cañardo Alastuey, J., Azom, R., Batra, P., Bezshapkin, V., Bolyen, E., Cagle, A., Caporaso, J.G., Debelius, J.W., Gorlick, K., Hamsanipally, N., Hunger, L., Keluskar, A., Liao, D., Lu, Y.Y., Navas-Molina, J.A., Pitman, A., Rideout, J.R., Sazonov, A., Sathappan, B., Schwarzberg Lipson, K., Sfiligoi, I., Tapo, C., Vázquez-Baeza, Y., Wu, Z., Xu, Z.Z., Ye, M.S., Zhao, J., Knight, R., Morton, J.T., Zhu, Q., 2026. Scikit-bio: a fundamental Python library for biological omic data analysis. Nat Methods 23, 274–276. 10.1038/s41592-025-02981-z

Bagnat, M., Gray, R.S., 2020. Development of a straight vertebrate body axis. Development 147, dev175794. 10.1242/dev.175794

Blanchon, S., Legendre, M., Bottier, M., Tamalet, A., Montantin, G., Collot, N., Faucon, C., Dastot, F., Copin, B., Clement, A., Filoche, M., Coste, A., Amselem, S., Escudier, E., Papon, J.-F., Louis, B., 2020. Deep phenotyping, including quantitative ciliary beating parameters, and extensive genotyping in primary ciliary dyskinesia. J Med Genet 57, 237–244. 10.1136/jmedgenet-2019-106424

Boehringer, A.S., Sanaat, A., Arabi, H., Zaidi, H., 2023. An active learning approach to train a deep learning algorithm for tumor segmentation from brain MR images. Insights Imaging 14, 141. 10.1186/s13244-023-01487-6

Boggon, A.K., Hastewell, A.D., Dunkel, J., Wan, K.Y., 2026. Embodied behavioural complexity in a ciliated microorganism. Nat Commun 17, 8445. 10.1038/s41467-026-75076-8

Borovina, A., Superina, S., Voskas, D., Ciruna, B., 2010. Vangl2 directs the posterior tilting and asymmetric localization of motile primary cilia. Nat Cell Biol 12, 407–412. 10.1038/ncb2042

Bradski, G., 2000. The OpenCV Library. Dr. Dobb’s Journal of Software Tools.

Cantaut-Belarif, Y., Sternberg, J.R., Thouvenin, O., Wyart, C., Bardet, P.-L., 2018. The Reissner Fiber in the Cerebrospinal Fluid Controls Morphogenesis of the Body Axis. Curr Biol 28, 2479-2486.e4. 10.1016/j.cub.2018.05.079

Castleman, V.H., Romio, L., Chodhari, R., Hirst, R.A., de Castro, S.C.P., Parker, K.A., Ybot-Gonzalez, P., Emes, R.D., Wilson, S.W., Wallis, C., Johnson, C.A., Herrera, R.J., Rutman, A., Dixon, M., Shoemark, A., Bush, A., Hogg, C., Gardiner, R.M., Reish, O., Greene, N.D.E., O’Callaghan, C., Purton, S., Chung, E.M.K., Mitchison, H.M., 2009. Mutations in radial spoke head protein genes RSPH9 and RSPH4A cause primary ciliary dyskinesia with central-microtubular-pair abnormalities. Am J Hum Genet 84, 197–209. 10.1016/j.ajhg.2009.01.011

Chioccioli, M., Feriani, L., Nguyen, Q., Kotar, J., Dell, S.D., Mennella, V., Amirav, I., Cicuta, P., 2019. Quantitative High-Speed Video Profiling Discriminates between DNAH11 and HYDIN Variants of Primary Ciliary Dyskinesia. Am J Respir Crit Care Med 199, 1436– 1438. 10.1164/rccm.201812-2256LE

Chou, H.-T., Apelt, L., Farrell, D.P., White, S.R., Woodsmith, J., Svetlov, V., Goldstein, J.S., Nager, A.R., Li, Z., Muller, J., Dollfus, H., Nudler, E., Stelzl, U., DiMaio, F., Nachury, M.V., Walz, T., 2019. The Molecular Architecture of Native BBSome Obtained by an Integrated Structural Approach. Structure 27, 1384–1394.e4. 10.1016/j.str.2019.06.006

Duldulao, N.A., Lee, S., Sun, Z., 2009. Cilia localization is essential for in vivo functions of the Joubert syndrome protein Arl13b/Scorpion. Development 136, 4033–4042. 10.1242/dev.036350

Dutcher, S.K., 2020. Asymmetries in the cilia of Chlamydomonas. Philos Trans R Soc Lond B Biol Sci 375, 20190153. 10.1098/rstb.2019.0153

Eilers, P.H.C., Boelens, H.F.M., 2005. Baseline Correction with Asymmetric Least Squares Smoothing.

Fedorov, A., Beichel, R., Kalpathy-Cramer, J., Finet, J., Fillion-Robin, J.-C., Pujol, S., Bauer, C., Jennings, D., Fennessy, F., Sonka, M., Buatti, J., Aylward, S., Miller, J.V., Pieper, S., Kikinis, R., 2012. 3D Slicer as an image computing platform for the Quantitative Imaging Network. Magn Reson Imaging 30, 1323–1341. 10.1016/j.mri.2012.05.001

Geyer, V.F., Howard, J., Sartori, P., 2022. Ciliary beating patterns map onto a low-dimensional behavioural space. Nat Phys 18, 332–337. 10.1038/s41567-021-01446-2

Gohlke, C., 2025. cgohlke/tifffile: v2025.10.16. 10.5281/zenodo.17373292

Grimes, D.T., Boswell, C.W., Morante, N.F.C., Henkelman, R.M., Burdine, R.D., Ciruna, B., 2016. Zebrafish models of idiopathic scoliosis link cerebrospinal fluid flow defects to spine curvature. Science 352, 1341–1344. 10.1126/science.aaf6419

Hamming, R., 1977. Digital filters. Prentice-Hall, Englewood Cliffs, N.J.

Hansen, J.N., Rassmann, S., Jikeli, J.F., Wachten, D., 2018. SpermQ–A Simple Analysis Software to Comprehensively Study Flagellar Beating and Sperm Steering. Cells 8, 10. 10.3390/cells8010010

Hansen, J.N., Rassmann, S., Stüven, B., Jurisch-Yaksi, N., Wachten, D., 2021. CiliaQ: a simple, open-source software for automated quantification of ciliary morphology and fluorescence in 2D, 3D, and 4D images. Eur Phys J E Soft Matter 44, 18. 10.1140/epje/s10189-021-00031-y

Jeong, I., Hansen, J.N., Wachten, D., Jurisch-Yaksi, N., 2022. Measurement of ciliary beating and fluid flow in the zebrafish adult telencephalon. STAR Protoc 3, 101542. 10.1016/j.xpro.2022.101542

Jocher, G., Qiu, J., 2024. Ultralytics YOLO11.

Juan, G.R.R.-S., Mathijssen, A.J.T.M., He, M., Jan, L., Marshall, W., Prakash, M., 2020. Multi-scale spatial heterogeneity enhances particle clearance in airway ciliary arrays. Nat Phys 16, 958–964. 10.1038/s41567-020-0923-8

Kempeneers, C., Seaton, C., Chilvers, M.A., 2017. Variation of Ciliary Beat Pattern in Three Different Beating Planes in Healthy Subjects. Chest 151, 993–1001. 10.1016/j.chest.2016.09.015

Kirillov, A., Mintun, E., Ravi, N., Mao, H., Rolland, C., Gustafson, L., Xiao, T., Whitehead, S., Berg, A.C., Lo, W.-Y., Dollár, P., Girshick, R., 2023. Segment Anything. 10.48550/arXiv.2304.02643

Knowles, M.R., Ostrowski, L.E., Leigh, M.W., Sears, P.R., Davis, S.D., Wolf, W.E., Hazucha, M.J., Carson, J.L., Olivier, K.N., Sagel, S.D., Rosenfeld, M., Ferkol, T.W., Dell, S.D., Milla, C.E., Randell, S.H., Yin, W., Sannuti, A., Metjian, H.M., Noone, P.G., Noone, P.J., Olson, C.A., Patrone, M.V., Dang, H., Lee, H.-S., Hurd, T.W., Gee, H.Y., Otto, E.A., Halbritter, J., Kohl, S., Kircher, M., Krischer, J., Bamshad, M.J., Nickerson, D.A., Hildebrandt, F., Shendure, J., Zariwala, M.A., 2014. Mutations in RSPH1 cause primary ciliary dyskinesia with a unique clinical and ciliary phenotype. Am J Respir Crit Care Med 189, 707–717. 10.1164/rccm.201311-2047OC

Kocak, B., Klonåas, M.E., Stanzione, A., Meddeb, A., Demircioğlu, A., Bluethgen, C., Bressem, K.K., Ugga, L., Mercaldo, N., Díaz, O., Cuocolo, R., 2025. Evaluation metrics in medical imaging AI: fundamentals, pitfalls, misapplications, and recommendations. European Journal of Radiology Artificial Intelligence 3, 100030. 10.1016/j.ejrai.2025.100030

Konjikusic, M.J., Yeetong, P., Boswell, C.W., Lee, C., Roberson, E.C., Ittiwut, R., Suphapeetiporn, K., Ciruna, B., Gurnett, C.A., Wallingford, J.B., Shotelersuk, V., Gray, R.S., 2018. Mutations in Kinesin family member 6 reveal specific role in ependymal cell ciliogenesis and human neurological development. PLOS Genetics 14, e1007817. 10.1371/journal.pgen.1007817

Kramer-Zucker, A.G., Olale, F., Haycraft, C.J., Yoder, B.K., Schier, A.F., Drummond, I.A., 2005. Cilia-driven fluid flow in the zebrafish pronephros, brain and Kupffer’s vesicle is required for normal organogenesis. Development 132, 1907–1921. 10.1242/dev.01772

Labun, K., Montague, T.G., Krause, M., Torres Cleuren, Y.N., Tjeldnes, H., Valen, E., 2019. CHOPCHOP v3: expanding the CRISPR web toolbox beyond genome editing. Nucleic Acids Res 47, W171–W174. 10.1093/nar/gkz365

Lechtreck, K.-F., Johnson, E.C., Sakai, T., Cochran, D., Ballif, B.A., Rush, J., Pazour, G.J., Ikebe, M., Witman, G.B., 2009. The Chlamydomonas reinhardtii BBSome is an IFT cargo required for export of specific signaling proteins from flagella. J Cell Biol 187, 1117–1132. 10.1083/jcb.200909183

Nishimura, D.Y., Searby, C.C., Carmi, R., Elbedour, K., Van Maldergem, L., Fulton, A.B., Lam, B.L., Powell, B.R., Swiderski, R.E., Bugge, K.E., Haider, N.B., Kwitek-Black, A.E., Ying, L., Duhl, D.M., Gorman, S.W., Heon, E., Iannaccone, A., Bonneau, D., Biesecker, L.G., Jacobson, S.G., Stone, E.M., Sheffield, V.C., 2001. Positional cloning of a novel gene on chromosome 16q causing Bardet–Biedl syndrome (BBS2). Human Molecular Genetics 10, 865–874. 10.1093/hmg/10.8.865

Nonaka, S., Tanaka, Y., Okada, Y., Takeda, S., Harada, A., Kanai, Y., Kido, M., Hirokawa, N., 1998. Randomization of Left–Right Asymmetry due to Loss of Nodal Cilia Generating Leftward Flow of Extraembryonic Fluid in Mice Lacking KIF3B Motor Protein. Cell 95, 829–837. 10.1016/S0092-8674(00)81705-5

Oltean, A., Schaffer, A.J., Bayly, P.V., Brody, S.L., 2018. Quantifying Ciliary Dynamics during Assembly Reveals Stepwise Waveform Maturation in Airway Cells. Am J Respir Cell Mol Biol 59, 511–522. 10.1165/rcmb.2017-0436OC

O’Mahony, N., Campbell, S., Carvalho, A., Harapanahalli, S., Hernandez, G.V., Krpalkova, L., Riordan, D., Walsh, J., 2020. Deep Learning vs. Traditional Computer Vision, in: Arai, K., Kapoor, S. (Eds.), Advances in Computer Vision. Springer International Publishing, Cham, pp. 128–144. 10.1007/978-3-030-17795-9_10

Pedregosa, F., Varoquaux, G., Gramfort, A., Michel, V., Thirion, B., Grisel, O., Blondel, M., Prettenhofer, P., Weiss, R., Dubourg, V., Vanderplas, J., Passos, A., Cournapeau, D., Brucher, M., Perrot, M., Duchesnay, É., 2011. Scikit-learn: Machine Learning in Python. Journal of Machine Learning Research 12, 2825–2830.

Praveen, K., Davis, E.E., Katsanis, N., 2015. Unique among ciliopathies: primary ciliary dyskinesia, a motile cilia disorder. F1000Prime Rep 7, 36. 10.12703/P7-36

Redmon, J., Divvala, S., Girshick, R., Farhadi, A., 2016. You Only Look Once: Unified, Real-Time Object Detection. 10.48550/arXiv.1506.02640

Ribeiro, A., Monteiro, J.F., Certal, A.C., Cristovão, A.M., Saúde, L., 2017. Foxj1a is expressed in ependymal precursors, controls central canal position and is activated in new ependymal cells during regeneration in zebrafish. Open Biol 7, 170139. 10.1098/rsob.170139

Rose, F., Michaluk, M., Blindauer, T., Ignatowska-Jankowska, B.M., O’Shaughnessy, L., Stephens, G.J., Pereira, T.D., Uusisaari, M.Y., Bozek, K., 2026. Deep Imputation for Skeleton data (DISK) for behavioral science. Nat Methods 23, 236–247. 10.1038/s41592-025-02893-y

Saggiorato, G., Alvarez, L., Jikeli, J.F., Kaupp, U.B., Gompper, G., Elgeti, J., 2017. Human sperm steer with second harmonics of the flagellar beat. Nat Commun 8, 1415. 10.1038/s41467-017-01462-y

Satir, P., Heuser, T., Sale, W.S., 2014. A Structural Basis for How Motile Cilia Beat. Bioscience 64, 1073–1083. 10.1093/biosci/biu180

Schindelin, J., Arganda-Carreras, I., Frise, E., Kaynig, V., Longair, M., Pieåsch, T., Preibisch, S., Rueden, C., Saalfeld, S., Schmid, B., Tinevez, J.-Y., White, D.J., Hartenstein, V., Eliceiri, K., Tomancak, P., Cardona, A., 2012. Fiji: an open-source platform for biological-image analysis. Nat Methods 9, 676–682. 10.1038/nmeth.2019

Seabold, S., Perktold, J., 2010. statsmodels: Econometric and statistical modeling with python. 9th Python in Science Conference. 10.25080/Majora-92bf1922-011

Shah, A.S., Farmen, S.L., Moninger, T.O., Businga, T.R., Andrews, M.P., Bugge, K., Searby, C.C., Nishimura, D., Brogden, K.A., Kline, J.N., Sheffield, V.C., Welsh, M.J., 2008. Loss of Bardet–Biedl syndrome proteins alters the morphology and function of motile cilia in airway epithelia. Proc. Natl. Acad. Sci. U.S.A. 105, 3380–3385. 10.1073/pnas.0712327105

Singh, S.K., Gui, M., Koh, F., Yip, M.C., Brown, A., 2020. Structure and activation mechanism of the BBSome membrane protein trafficking complex. eLife 9, e53322. 10.7554/eLife.53322

Smith, C.M., Djakow, J., Free, R.C., Djakow, P., Lonnen, R., Williams, G., Pohunek, P., Hirst, R.A., Easton, A.J., Andrew, P.W., O’Callaghan, C., 2012. ciliaFA: a research tool for automated, high-throughput measurement of ciliary beat frequency using freely available software | Cilia | Springer Nature Link. 10.1186/2046-2530-1-14

Song, P., Fogerty, J., Cianciolo, L.T., Stupay, R., Perkins, B.D., 2020. Cone Photoreceptor Degeneration and Neuroinflammation in the Zebrafish Bardet-Biedl Syndrome 2 (bbs2) Mutant Does Not Lead to Retinal Regeneration. Front. Cell Dev. Biol. 8. 10.3389/fcell.2020.578528

Sternberg, J.R., Prendergast, A.E., Brosse, L., Cantaut-Belarif, Y., Thouvenin, O., Orts-Del’Immagine, A., Castillo, L., Djenoune, L., Kurisu, S., McDearmid, J.R., Bardet, P.-L., Boccara, C., Okamoto, H., Delmas, P., Wyart, C., 2018. Pkd2l1 is required for mechanoception in cerebrospinal fluid-contacting neurons and maintenance of spine curvature. Nat Commun 9, 3804. 10.1038/s41467-018-06225-x

Tait, C.M., Chinnaiya, K., Manning, E., Murtaza, M., Ashton, J.-P., Furley, N., Hill, C.J., Alves, C.H., Wijnholds, J., Erdmann, K.S., Furley, A., Rashbass, P., Das, R.M., Storey, K.G., Placzek, M., 2020. Crumbs2 mediates ventricular layer remodelling to form the spinal cord central canal. PLOS Biology 18, e3000470. 10.1371/journal.pbio.3000470

Thouvenin, O., Cantaut-Belarif, Y., Keiser, L., Gallaire, F., Wyart, C., 2021. Automated Analysis of Cerebrospinal Fluid Flow and Motile Cilia Properties in The Central Canal of Zebrafish Embryos [WWW Document]. URL https://bio-protocol.org/en/bpdetail?id=3932&type=0 (accessed 4.15.25).

Thouvenin, O., Keiser, L., Cantaut-Belarif, Y., Carbo-Tano, M., Verweij, F., Jurisch-Yaksi, N., Bardet, P.-L., van Niel, G., Gallaire, F., Wyart, C., 2020. Origin and role of the cerebrospinal fluid bidirectional flow in the central canal. eLife 9, e47699. 10.7554/eLife.47699

Tkachenko, M., Malyuk, M., Holmanyuk, A., Liubimov, N., 2020. Label studio: Data labeling software. Open source software available from https://github.com/HumanSignal/label-studio 2022.

Traub, L.M., Downs, M.A., Westrich, J.L., Fremont, D.H., 1999. Crystal structure of the α appendage of AP-2 reveals a recruitment platform for clathrin-coat assembly. Proceedings of the National Academy of Sciences 96, 8907–8912. 10.1073/pnas.96.16.8907

Troutwine, B.R., Gontarz, P., Konjikusic, M.J., Minowa, R., Monstad-Rios, A., Sepich, D.S., Kwon, R.Y., Solnica-Krezel, L., Gray, R.S., 2020. The Reissner Fiber Is Highly Dynamic In Vivo and Controls Morphogenesis of the Spine. Current Biology 30, 2353–2362.e3. 10.1016/j.cub.2020.04.015

Walker, B.J., Phuyal, S., Ishimoto, K., Tung, C.-K., Gaffney, E.A., 2020. Computer-assisted beat-pattern analysis and the flagellar waveforms of bovine spermatozoa. R Soc Open Sci 7, 200769. 10.1098/rsos.200769

Walt, S. van der, Schönberger, J.L., Nunez-Iglesias, J., Boulogne, F., Warner, J.D., Yager, N., Gouillart, E., Yu, T., 2014. scikit-image: image processing in Python. PeerJ 2, e453. 10.7717/peerj.453

Wang, Y., Troutwine, B.R., Zhang, H., Gray, R.S., 2022. The axonemal dynein heavy chain 10 gene is essential for monocilia motility and spine alignment in zebrafish | Elsevier Enhanced Reader. 10.1016/j.ydbio.2021.12.001

Wohlgemuth, K., Hoersting, N., Koenig, J., Loges, N.T., Raidt, J., George, S., Cindrić, S., Schramm, A., Biebach, L., Lay, S., Dougherty, G.W., Olbrich, H., Pennekamp, P., Dworniczak, B., Omran, H., 2024. Pathogenic variants in CFAP46, CFAP54, CFAP74 and CFAP221 cause primary ciliary dyskinesia with a defective C1d projection of the central apparatus. European Respiratory Journal 64. 10.1183/13993003.00790-2024

Xu, Y., Quan, R., Xu, W., Huang, Y., Chen, X., Liu, F., 2024. Advances in Medical Image Segmentation: A Comprehensive Review of Traditional, Deep Learning and Hybrid Approaches. Bioengineering (Basel) 11, 1034. 10.3390/bioengineering11101034

Zhang, Y., Sun, P., Jiang, Y., Yu, D., Weng, F., Yuan, Z., Luo, P., Liu, W., Wang, X., 2022. ByteTrack: Multi-object Tracking by Associating Every Detection Box, in: Avidan, S., Brostow, G., Cissé, M., Farinella, G.M., Hassner, T. (Eds.), Computer Vision – ECCV 2022, Lecture Notes in Computer Science. Springer Nature Swiåerland, Cham, pp. 1–21. 10.1007/978-3-031-20047-2_1

Zijdenbos, A.P., Dawant, B.M., Margolin, R.A., Palmer, A.C., 1994. Morphometric analysis of white matter lesions in MR images: method and validation. IEEE Transactions on Medical Imaging 13, 716–724. 10.1109/42.363096

